# NTBC dosing and outcomes in hereditary tyrosinemia type 1: insights from a representative human model and 99 patients

**DOI:** 10.1101/2025.10.06.680797

**Authors:** QT Pham, F Tamnanloo, MA M’Callum, E Beaulieu, R Mghabghab, Y Théorêt, G Mitchell, the Quebec NTBC Study Group, D Cyr, PJ Waters, Y Doyon, U Halac, C Raggi, M Paganelli

**Affiliations:** Liver Tissue Engineering and Cell Therapy Laboratory, Azrieli Research Center, CHU Sainte-Justine, Montreal, Quebec, Canada; Department of Pediatrics, University of Montreal, Montreal, Canada; Hôtel-Dieu-de-Lévis, Department of Medicine and Faculty of Medicine, Laval University, Quebec City, Quebec, Canada; Pharmacology Research Unit, Azrieli Research Center, CHU Sainte-Justine, Montreal, Quebec, Canada; Genetics, CHU Sainte-Justine, Montreal, Quebec, Canada; The Quebec NTBC Study Group: Fernando Alvarez, Suzanne Atkinson, Manon Bouchard, Catherine Brunel-Guitton, Daniela Buhas, Jean-François Bussières, Josée Dubois, Daphna Fenyves, Paul Goodyer, Martyne Gosselin, Ugur Halac, Patrick Labbé, Rachel Laframboise, Bruno Maranda, Serge Melançon, Aicha Merouani, Grant A Mitchell, John Mitchell, Guy Parizeault, Luc Pelletier, Véronique Phan, Jean-François Turcotte; Medical Genetics Service, Dept. Laboratory Medicine, CHU of Sherbrooke (CHUS) and Dept. Pediatrics, University of Sherbrooke, Sherbrooke, Quebec, Canada; CHUS Research Centre (CRCHUS), Sherbrooke, Quebec, Canada; Centre Hospitalier Universitaire de Québec Research Center and Faculty of Medicine, Laval University, Québec City, Quebec, Canada; Pediatric Gastroenterology, Hepatology and Nutrition, CHU Sainte-Justine, Montreal, Quebec, Canada; Morphocell Technologies Inc., Laval, Quebec, Canada

**Keywords:** genetic liver disease, tyrosinemia type 1, nitisinone, NTBC, induced pluripotent stem cells, hepatocyte-like cells, disease modeling, isogeneic controls

## Abstract

Hereditary tyrosinemia type 1 (HT1) is a rare and severe metabolic liver disorder caused by fumarylacetoacetate hydrolase (FAH) deficiency. The optimal dose and long-term effects of the only available treatment, nitisinone (NTBC), remain unclear due to the absence of clinical trial data. Here, we generated a representative human in vitro model of HT1 using iPSC-derived hepatocytes, which faithfully recapitulated key disease features. We investigated the mechanisms of FAH deficiency-induced hepatocellular injury and evaluated the effects of NTBC treatment. We confirmed treatment efficacy and identified 50 µmol/L as the minimal effective NTBC concentration to prevent cellular damage. This protective dose was subsequently validated in a large cohort of 99 HT1 patients, providing compelling evidence for establishing minimal therapeutic NTBC levels. Notably, approximately 10% of disease-associated genes, many implicated in hepatocellular carcinoma, remained dysregulated despite treatment, raising concerns that NTBC may not fully eliminate long-term oncogenic risk.

## Introduction

Hereditary tyrosinemia type 1 (HT1, OMIM #276700) is a severe inborn error of liver metabolism that, if untreated, leads to mortality before 2 years of age in over 90% of affected children.^1,2^ HT1 originates from the deficiency of fumarylacetoacetate hydrolase (FAH), the enzyme carrying out the last step of the tyrosine degradation (Fig. 1A).^3^ FAH deficiency causes the accumulation of toxic metabolites, notably mutagenic fumarylacetoacetate (FAA) and maleylacetoacetate (MAA) and their derivative succinylacetone (SA). The latter is stable and therefore easier to measure, and is considered a specific diagnostic biomarker of the disease.^4,5^ Untreated, HT1 results in severe liver injury with impairment of hepatocyte synthetic functions and rapidly progressive liver failure in most patients.^2,6^ Liver disease develops with progressive fibrosis and cirrhosis, occasional episodes of acute liver failure or hepatic porphyria-like neurologic crises and renal Fanconi syndrome. HT1 patients are also at high risk of rapidly developing hepatocellular carcinoma (HCC). FAA, but not MAA or SA, was proven to be mutagenic *in vitro* and *in vivo.*^4,7,8^ Whereas FAA tumorigenicity was well proven and several mechanistic hypotheses involving endoplasmic reticulum stress response, heat shock response and resistance to apoptosis, among others, were formulated,^9^ very little is known about how the accumulation of toxic tyrosine metabolites leads to the severe hepatocellular injury observed in patients and animal models.

**Figure 1.**
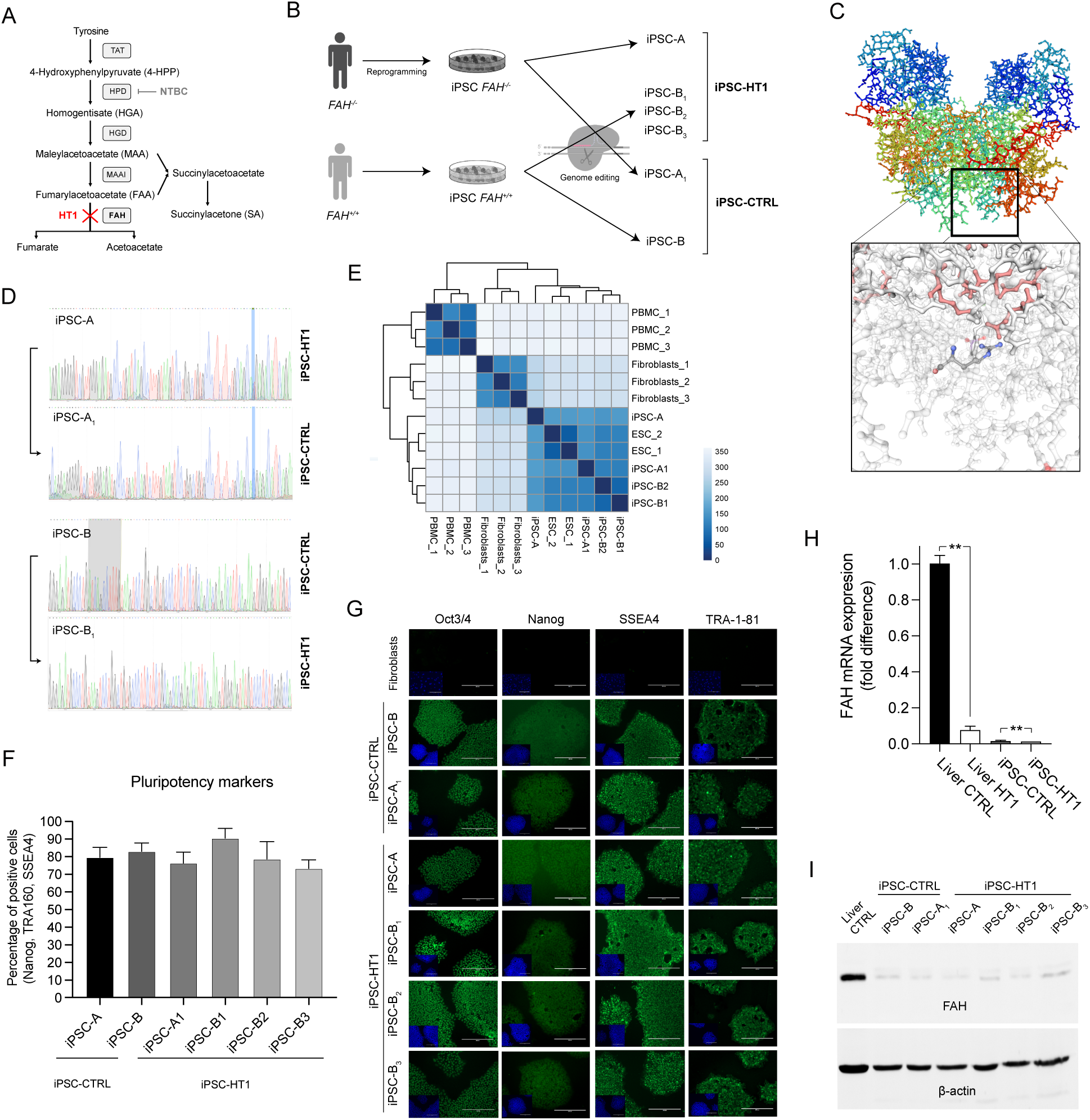
**A)** Pathway of tyrosine metabolism in HT1. **B)** iPSC groups including FAH deficient, healthy, and isogenic controls. **C)** 3D structure of FAH enzyme and location of p.W237R aminoacidic change resulting from exon 9 mutation in iPSC-A. **D**) Sequencing chromatogram of *FAH* gene in the iPSC lines showing the corrected mutation in iPSC-A_1_ and induced deletion in iPSC-B_1_. **E)** The transcriptomic profiles of iPSC lines are closely aligned and comparable to those of embryonic stem cells (ESC, clustering heatmap). **F-G**) No difference was observed in the expression of pluripotency markers between iPSC-HT1 and iPSC-CTRL at (F: flow cytometry, percentage of Nanog/TRA160/SSEA4-positive cells; G: fluorescence immunostaining, scale bar 200 µm). **H-I)** When compared to healthy liver tissue, expression of liver-specific *FAH* was negligible in iPSC and significantly lower in iPSC-HT1 than iPSC-CTRL (H: RT-qPCR, N=3 to 16, mean±SEM, **p<0.01; I: Western blot, N=3, representative image).

Treatment of HT1 evolved over the years. Tyrosine-free diet since birth, the unique therapeutic option for several years, still resulted in 90% mortality of affected children by 12 years of age.^6,10^ In the 80’s and early 90’s, liver transplantation (LT) in childhood significantly improved survival rates.^10^ HT1 treatment was then revolutionized when Lindstedt and colleagues first described the use of herbicide 2-(2-N-4-trifluoromethylbenzoyl)-1,3-cyclohexanedione (nitisinone or NTBC) to prevent the formation of tyrosine’s toxic metabolites in HT1.^11^ NTBC, a potent inhibitor of 4-hydroxyphenylpyruvate dioxygenase (HPD), proved to be mostly safe in rodents and primates when combined with low tyrosine-phenylalanine diet.^4,12–14^ In children with HT1, NTBC was shown to be effective in suppressing SA levels, preventing the progression of liver and kidney disease and eliminating the need for LT.^8,15^ Although tyrosinemic mice on the drug still develop HCC on the long term, and cancers have been observed in patients who started treatment later in life, no case of HCC has been described so far in patients properly receiving NTBC since birth.^16^ Approved without going through classical phases of clinical testing, the treatment has been routinely used to treat children with HT1 since birth for almost 30 years (together with dietary restrictions), and has shown a reassuring safety profile.^15^ The standard dose of NTBC used in daily clinical practice was established based on clinical observation.^17^ Nevertheless, the dose required to prevent liver cellular injury and potential long-term complications is still unknown, and targeted NTBC blood levels have been arbitrarily suggested because of the lack of solid data and suitable models.^18^ The *Fah^-/-^* mouse model greatly advanced our knowledge of HT1 pathophysiology and therapy;^8,16,19–22^ however, there are differences between FAH deficiency in mice and humans, especially for HCC development.^14,16^ Additionally, collection of human data on the pathophysiology of the disease has been hampered by the lack of *in vitro* human models.

Primary human hepatocytes (PHH), which are the “gold standard” for *in vitro* modeling and drug testing, suffer from challenging limitations (scarce availability, high variability among donors, rapid dedifferentiation upon *in vitro* culture and loss of functions upon cryopreservation).^23–25^ Transformed cell lines offer enhanced accessibility and reproducibility but are limited in their ability to fully represent the complexity of biological systems. In the past two decades, several groups including ours have harnessed stem cells from various sources to produce hepatocyte-like cells (HLC), aiming to model a range of liver diseases.^26^ HLC derived from induced pluripotent stem cells (iPSC), which can be easily reprogrammed from somatic cells of patients and controls, proved especially promising to model genetic liver diseases,^27,28^ despite differentiation protocols often leading to a phenotype more similar to fetal than adult PHH, with low yield.^29^ We recently described a robust protocol to consistently generate more mature hepatocytes from human iPSC.^30^ Also, we can effectively generate isogeneic controls through CRISPR/Cas9-based genome editing of healthy and patient-derived iPSC.^31^ In this study, by using such tools, we developed a representative, human, *in vitro* model of HT1 to study the consequences of FAH deficiency on hepatocytes and assess the effects of NTBC. This allowed us to recreate HT1-induced liver injury in a controlled environment and identify the optimal dose of NTBC necessary to prevent such cellular injury. We have validated our *in vitro* results using clinical data from a large cohort of HT1 patients. Our transcriptomic data revealed upregulation of pro-fibrotic and pro-inflammatory genes in FAH-deficient hepatocytes, the majority of which were corrected by NTBC treatment. Nevertheless, we identified a small population of genes in which dysregulation persisted despite NTBC treatment.

## Results

### Generation and characterization of FAH-deficient iPSC lines and isogeneic controls

We generated 4 FAH-deficient iPSC lines (iPSC-HT1) and 2 isogeneic controls (iPSC-CTRL; Fig. 1B). A first FAH-deficient iPSC line (iPSC-A) was reprogrammed from CD34-enriched peripheral blood mononuclear cells of an adolescent male suffering from HT1 and carrying an heterozygous c.709C>T mutation (p.W237R) in exon 9 and a deletion in exon 13 and 14 of the *FAH* gene (Fig. 1C), and extensively characterized (Fig. 1F, G).

We then obtained 3 additional FAH-deficient iPSC populations (iPSC-B1, iPSC-B2, iPSC-B3) by inducing pathogenic homozygous mutations (all in exon 9) in a previously characterized iPSC line derived from a healthy female subject.^30,31^ In order to avoid effects of confounding factors potentially due to interindividual variability, we generated isogeneic controls (Fig. 1B and 1D): the c.709C>T *FAH* mutation was corrected in patient-derived iPSC by CRISPR-Cas9^D10A^ nickase-based, homology-directed repair, thus obtaining the first isogeneic control (iPSC-A_1_), whereas the original healthy iPSC line served as isogeneic control (iPSC-B) of the other 3 populations.

No significant differences were noted among the 6 iPSC populations in terms of self-renewal potential. Transcriptomic profiles were comparable to those of embryonic stem cells (Fig. 1E and Supp. Fig. 1A). The expression of pluripotency markers of the 6 populations was comparable and not affected by genome editing (Fig. 1F, G and Supp. Fig. 1B). *FAH* gene expression was very low in iPSC-CTRL compared to liver tissue (≤2%), and even lower in iPSC-HT1 (*p*=0.007; Fig. 1H). Accordingly, FAH protein expression was negligible in all undifferentiated iPSC clones (Fig. 1I).

### Generation of FAH-deficient hepatocytes and isogenic controls

We differentiated the iPSC populations into HLC using our previously published protocol that recapitulates the progressive interactions between signaling pathways observed during liver organogenesis (Fig. 2A).^30^ Homogenous populations of HLC were consistently generated from both iPSC-CTRL and iPSC-HT1, with no difference in morphology after 4 weeks (Fig. 2B and Supp. Fig. 2A). Nevertheless, significant cell death was observed in iPSC-HT1 starting from day 16 of the differentiation (coinciding with the progressive acquisition of liver-specific functions).^30^ Cell death was eliminated when differentiating cells were supplemented with NTBC starting from day 16. After 4 weeks of differentiation, HLC acquired phenotype and functions consistent with neonatal hepatocytes, with negligible difference between HLC derived from iPSC-HT1 (HLC-HT1) and those derived from iPSC-CTRL (HLC-CTRL; Fig. 2B-F, and Supp. Fig. 2A, B). Expression of the albumin (*ALB*) and hepatocyte nuclear factor 4 α (*HNF4A*) genes, but not of the alpha-fetoprotein’s (*AFP*) gene, was higher in HLC-HT1 than in HLC-CTRL (Fig. 2D). Nevertheless, albumin was produced and secreted at comparable levels regardless of the presence of *FAH* deficiency (Fig. 2D, F and Supp. Fig. 2C), and no difference was observed in cytochrome P450 3A4 activity (Fig. 2E and Supp. Fig. 2D). As expected, *FAH* expression was substantial in HLC-CTRL (although <50% of that in the healthy human liver, likely due to HLC’s incomplete maturity), while still negligible in HLC-HT1 (Fig. 2G, H). The genes coding for the other enzymes involved in tyrosine metabolism, including tyrosine aminotransferase (TAT), homogentisic acid oxidase (HGD), and maleylacetoacetate isomerase (MAAI; also known as glutathione S-transferase zeta 1 - GSTZ1), were equally expressed in HLC, except for HPD. The latter was more expressed in HLC-HT1, with higher mRNA levels possibly due to the protein’s inhibition by NTBC (Supp. Fig. 2E).

**Figure 2.**
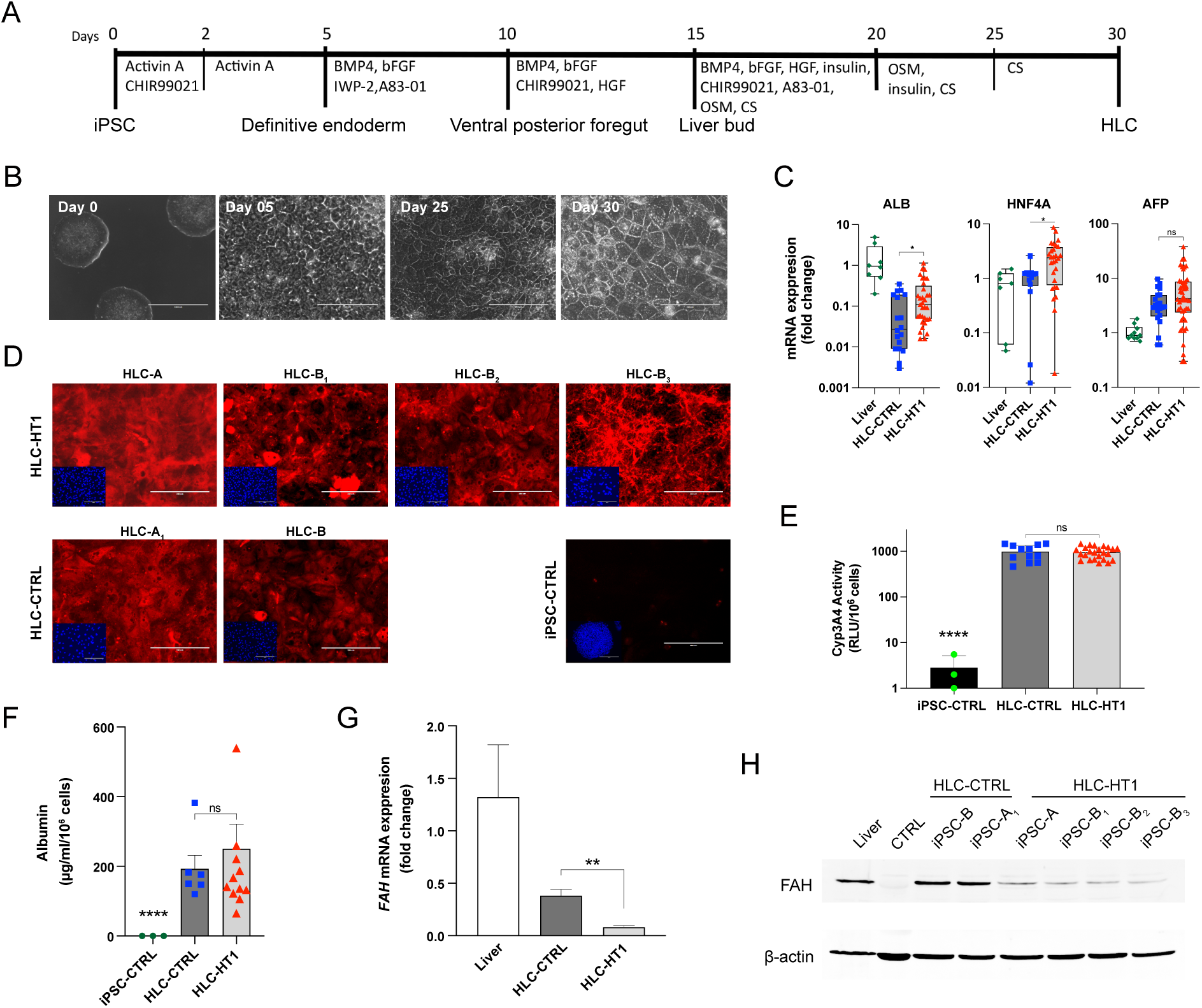
**A)** Differentiation protocol used to generate HLC from iPSC. **B)** Cell morphology through different differentiation stages from iPSC (day 0) to mature HLC (phase contrast, scale bar, from left: 1000 µm, 100 µm, 200 µm, 200 µm, representative images). **C)** mRNA expression of *ALB, HNF4A* and *AFP* was slightly higher in mature HLC-HT1 than in HLC-CTRL (RT-qPCR, N=10-to-46, mean±SEM, *p<0.05; healthy liver tissue served as positive control). **D)** Albumin was highly expressed in all HLC (fluorescence immunostaining, scale bar 200 µm, N=3, representative images). **E)** HLC-CTRL and HLC-HT1 showed comparable Cyp3A4 activity (normalized enzyme activity, N=3-to-24, mean±SEM). **F)** All HLC showed high albumin secretion levels, independently from *FAH* mutation (ELISA, N=3-to-12, mean±SEM, ****p<0.0001). **G)** *FAH* mRNA levels were significantly higher in HLC-CRTL than in HLC-HT1 (RT-qPCR, N=3, mean ± SEM, **p<0.01). **H)** FAH protein is well expressed in HLC-CTRL but barely detectable in HLC-HT1 (Western blot, N=3, representative image).

### HT1 disease phenotype in FAH-deficient hepatocytes

In order to assess whether HLC-HT1 could recapitulate the human HT1 disease phenotype *in vitro*, we measured SA in conditioned media upon withdrawal of NTBC (see online methods) and supplementation with L-tyrosine. Supplementation with 300 µmol/L tyrosine for 24 hours caused a significant accumulation of SA in HLC-HT1 culture supernatant, whereas SA remained undetectable in HLC-CTRL (Fig. 3A). Since the expression of TAT in HLC is lower than in the human liver (Supp. Fig. 2E), we also assessed the response of the cells to incubation with intermediary metabolites of tyrosine downstream of TAT. As expected, an increase in SA was observed when HLC-HT1 were supplemented with either 300 µmol/L 4-HPP (Fig. 3B) or 300 µmol/L HGA (Fig. 3C), whereas no increase was registered for HLC-CTRL. In FAH-deficient cells, tyrosine supplementation resulted in significant cell death, with HLC-HT1 (but not HLC-CTRL) showing >30% apoptotic cell death over the first 24 hours (Fig. 3D). Cell death rate was higher (>80%) when the cells were supplemented with 4-HPP and HGA, which can be explained by the previously mentioned relatively low TAT expression in HLC.

**Figure 3.**
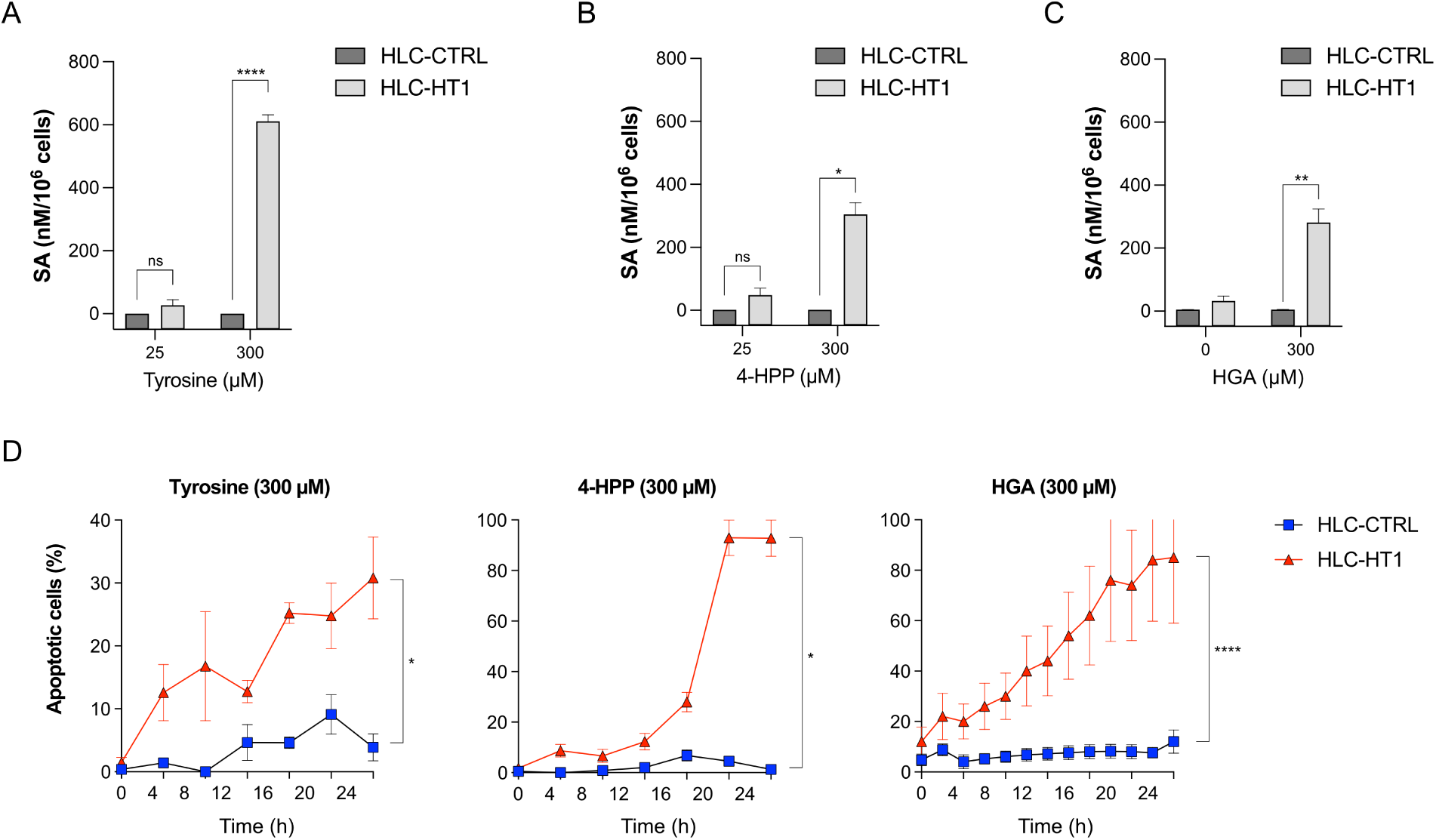
**A)** Supplementation with 300 µM tyrosine resulted in increased SA levels in the culture supernatant of HLC-HT1 but not of HLC-CTRL (N=3, mean±SEM, ****p<0.0001). **B-C)** SA levels increased in the supernatant of HLC-HT1, but not HLC-CTRL, upon supplementation with 300 µM 4-HPP and HGA (N=3, mean±SEM, *p<0.05, **p<0.01). **D)** HLC-HT1 showed significant apoptosis upon NTBC withdrawal and supplementation with 300 µM tyrosine, 4HPP or HGA, whereas no significant increase in cell death was observed in HLC-CTRL (N=3, *p<0.05, ****p<0.0001).

### NTBC prevents hepatocellular injury in HT1

As mentioned above, our *in vitro* model required daily NTBC supplementation starting from day 16 of the differentiation process to prevent cell death caused by the effect of defective tyrosine metabolism. NTBC supplementation to FAH-expressing hepatocytes (HLC-CTRL) caused no apoptosis for doses up to 50 µM, whereas addition of 100 µM NTBC resulted in increased cell death (Fig. 4A). Supplementation of 300 µM L-tyrosine was well tolerated by the liver cells, whereas a higher dose, with or without NTBC, increased apoptosis (Fig. 4A). In FAH-deficient HLC-HT1, 50 µM NTBC or more prevented accumulation of SA, even upon supplementation with 300 µM L-tyrosine, whereas a lower dose (20 µM) of NTBC was insufficient to inhibit SA buildup (Fig. 4B). In tyrosinemic hepatocytes, but not in HLC-CTRL, NTBC withdrawal and supplementation with 300 µM L-tyrosine resulted in significant apoptosis already within the first 24 hours (Fig. 4C). Treatment with 20 µM NTBC significantly reduced but did not eliminate cell death, whereas 50 µM NTBC were needed to bring apoptosis down to the negligible levels observed in controls. Conversely, treatment with 100 µM NTBC (still in the presence of 300 µM L-tyrosine supplementation) failed to suppress cell death in HLC-HT1 and increased apoptosis in HLC-CTRL (Fig. 4D).

**Figure 4.**
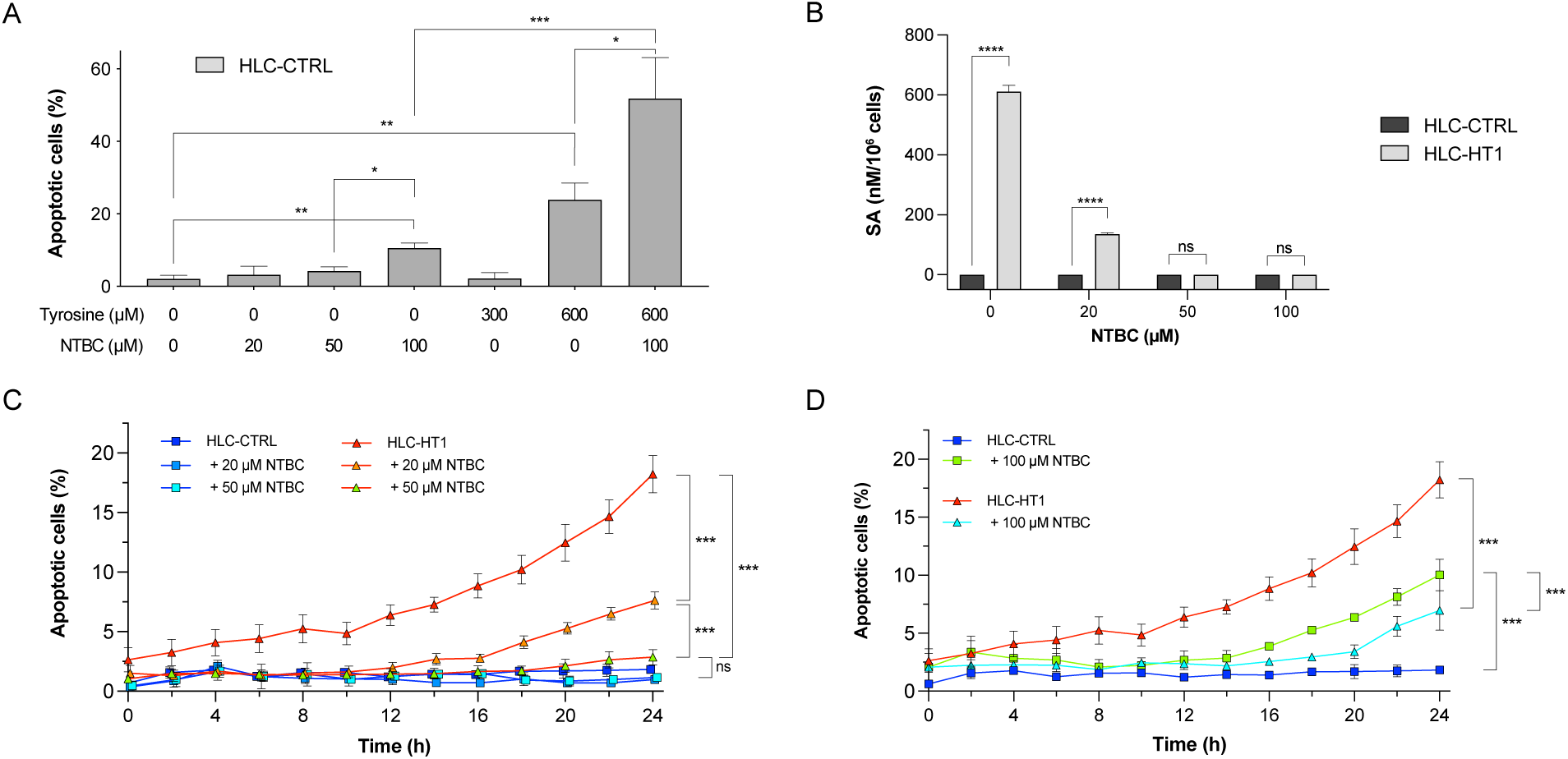
**A)** NTBC levels equal or lower than 50 µM did not result in any measurable cell death when supplemented to HLC-CTRL. Nevertheless, supplementation of 100 µM of NTBC and/or 600 µM tyrosine led to apoptosis (N=3, mean±SEM, *p<0.05). **B)** Supplementation of NTBC reduced SA levels in HLC-HT1, with ≥50 µM needed to reduce SA to undetectable levels (N=4 to 8, mean±SEM, *p<0.05). **C-D)** 50 µM NTBC effectively prevented apoptosis in HLC-HT1, whereas 20 µM NTBC was not sufficient to prevent it. No significant cell death was observed in HLC-CTRL with either NTBC concentration, while 100 µM NTBC led to apoptosis in both HLC-HT1 and HLC-CTRL (N=4, mean±SD, ***p<0.001).

### Identified NTBC threshold protects patients with HT1 from liver injury

We validated the identified effective dose of NTBC in a retrospective cohort of 99 HT1 patients (Table 1 and Fig. 5). Overall, we gathered 358 annual measurements over 48 months. For each measurement, subjects were divided into two groups based on NTBC plasma trough levels, using 50 µM as a threshold.

**Figure 5.**
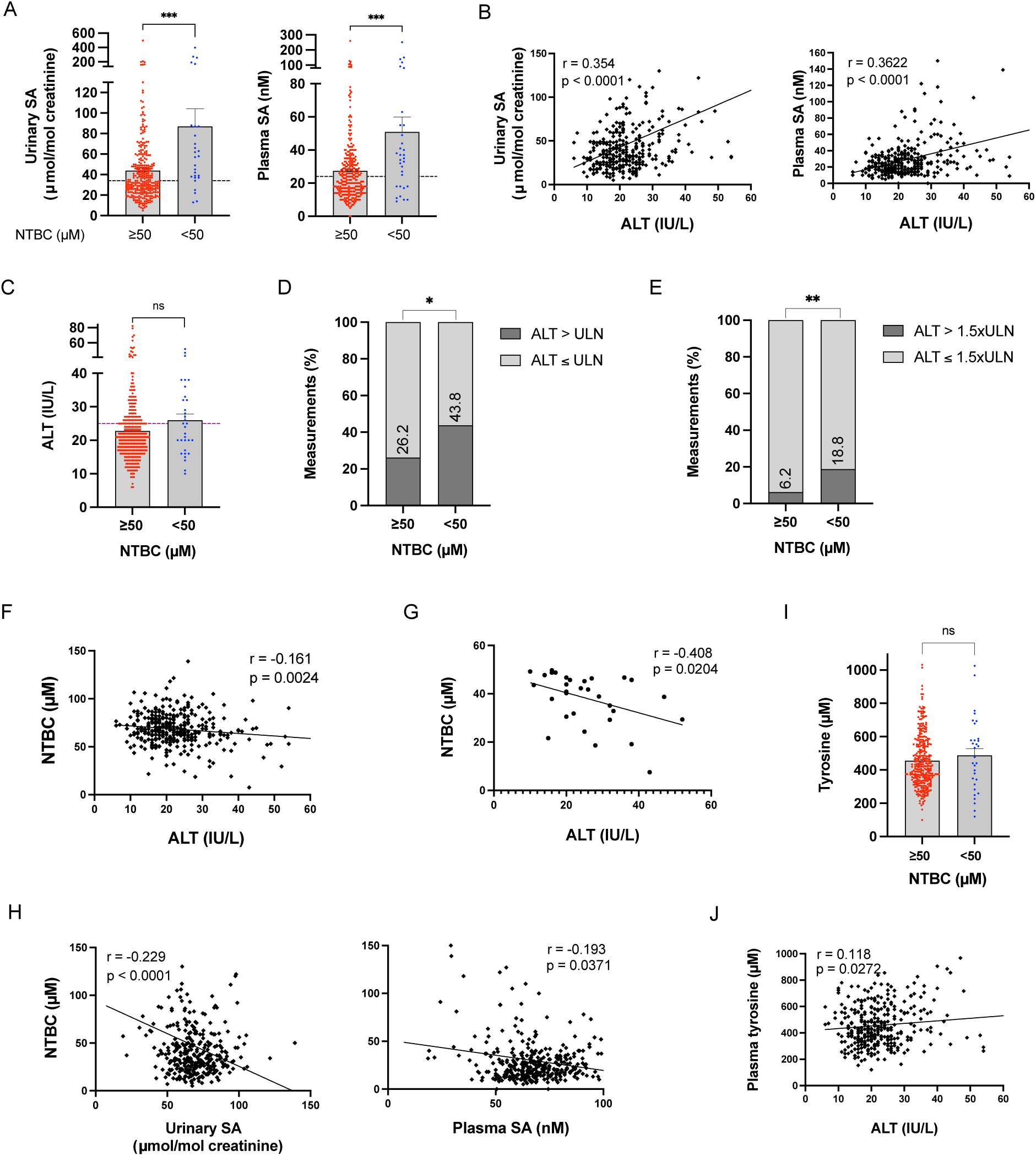
**A)** Urinary and plasma SA levels of 99 patients with HT1 were significantly lower when treated with ≥50 µM of NTBC (N=28/308 and N=32/317 urinary and plasma SA measurements, respectively; mean±SEM and scatter plot, ***p=0.0004 and p=0.0008, respectively; dashed lines showing the upper limits of normal). **B)** A significant positive correlation was detected between urinary and plasma SA and plasma ALT levels (N=334 and N=348, respectively, p<0.0001). **C)** Plasma ALT levels were slightly higher in patients with NTBC trough levels <50 µM, although the difference did not reach statistical significance (N=32/317, mean±SEM and scatter plot, p=0.069). **D)** Plasma ALT levels exceeded the ULN more often in patients with NTBC trough levels <50 µM (N=353, *p=0.0342). **E)** A greater proportion of abnormal ALT measurements (>1.5xULN) were observed in patients with NTBC <50 µM compared to those with NTBC ≥50 µM (N=353, **p=0.0097). **F-G)** NTBC plasma levels showed a weak but significant negative correlation with ALT, which strengthened when focusing on NTBC trough plasma levels <50 µM (N=351 and N=32, respectively; p=0.0024 and p=0.0204). **H)** A significant inverse correlation was observed between NTBC and SA levels (N=336 and N=350, respectively; p<0.0001 and p=0.0003, respectively). **I)** Tyrosine levels did not differ between patients with NTBC trough plasma levels <50 µM or ≥50 µM, despite 56.6% of measurements exceeding 400 µM (N=316/31, mean±SEM and scatter plot, ***p=0.465). **J)** The correlation between ALT and tyrosine plasma levels was weak across all measurements (N=348, p<0.0272).

**Table 1.**
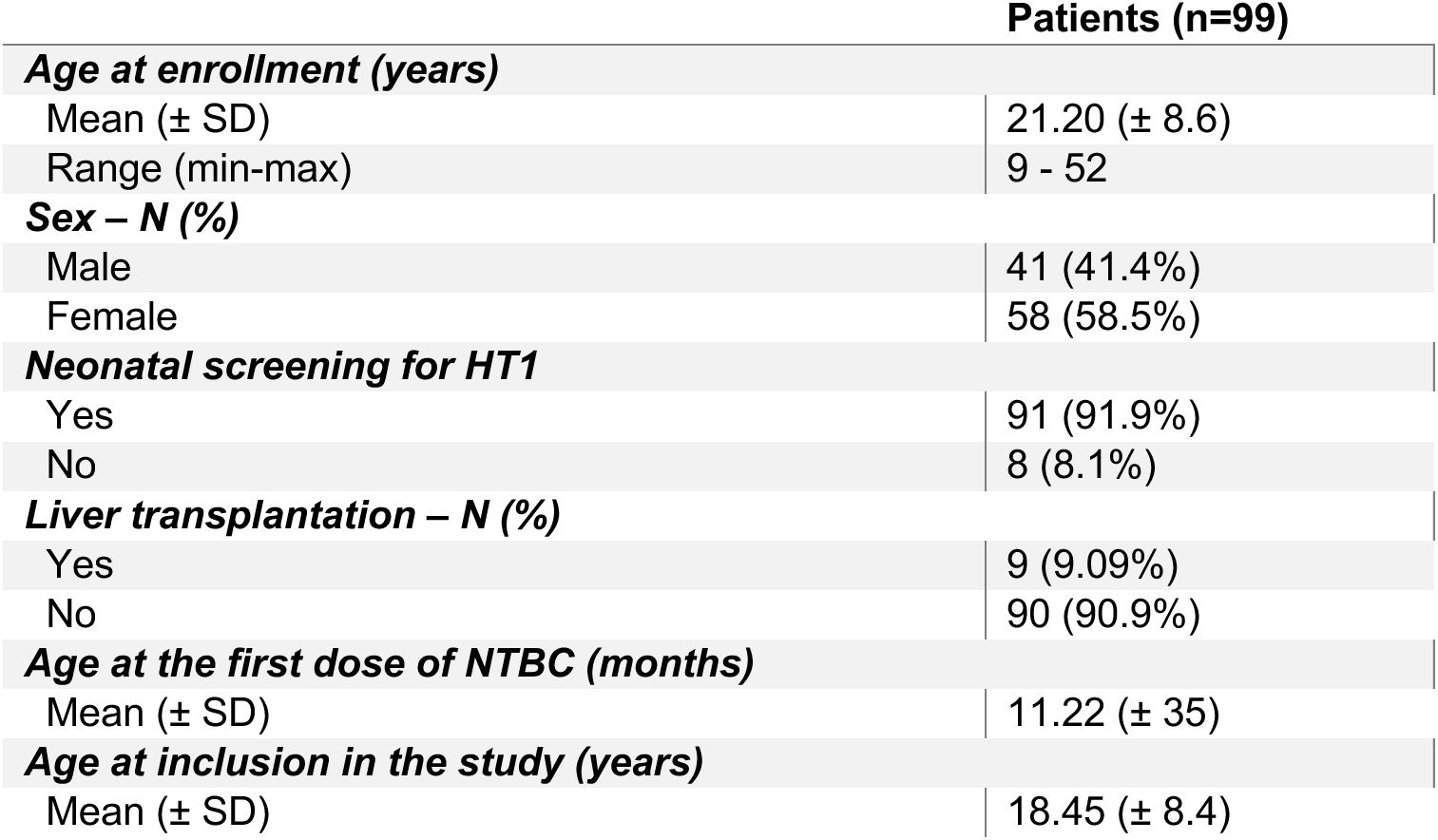
Patient demographics and clinical features.

Patients receiving less than 50 µM NTBC exhibited significantly higher urinary and plasma SA levels compared to those receiving 50 µM or more (Fig. 5A). ALT plasma levels showed a positive correlation with urinary and plasma SA levels (Fig. 5B) and were not statistically different between subjects with NTBC trough levels <50 µM or ≥50 µM. Nevertheless, for the former, they were above the upper limit of normal (ULN, >25 IU/L; Fig. 5C). With NTBC levels <50 µM, 43.8% of the annual measurements resulted in abnormal ALT levels (>1.5xULN in 18.8%), whereas 26.2% of the measurements were abnormal in patients with NTBC ≥50 µM (only 6.2% >1.5xULN; Fig. 5D and E). NTBC levels showed a weak but significant negative correlation with ALT plasma levels (Fig. 5F), correlation that became stronger when only measurements with NTBC <50 µM where considered (Fig. 5G). A significant correlation with NTBC levels was also seen for urinary and plasma SA levels (Fig. 5H). All patients being on a strict diet low in tyrosine and phenylalanine, average tyrosine plasma levels were within targeted range (458 ± 165 µM). Nevertheless, 56.6% of the measurements resulted in tyrosine levels >400 µM (median 548 µM, ma× 1032 µM). Interestingly, no difference in tyrosine levels was observed in patients based on whether NTBC plasma levels were ≥50 µM (Fig. 5I), and correlation with ALT levels was weak (Fig. 5J).

### *FAH* mutation affects the expression of genes responsible for HT1 pathophysiology

The analysis of the transcriptome showed a clear difference between pooled HLC-HT1 (after NTBC withdrawal) and HLC-CTRL, with 1058 differentially expressed genes (DEG), including 648 upregulated and 410 downregulated genes (Fig. 6A). Differences were noted between clones derived from different donors (Supp. Fig. 3B), with 1601 DEG between cells differentiated from patient-derived iPSC (HLC-A) and its isogenic control (HLC-A1), and 2319 DEG between cells differentiated from originally healthy iPSC (HLC-B) and one of its FAH-deficient counterparts (HLC-B1; Fig. 6B). A total of 342 DEG were common between the two genetic backgrounds, while only 216 genes showed a strong and significant differential expression in all groups analyzed, including 156 upregulated and 60 downregulated genes, which confirmed the importance of using an approach based on isogeneic controls (Fig. 6B, Supp Table 1).

**Figure 6.**
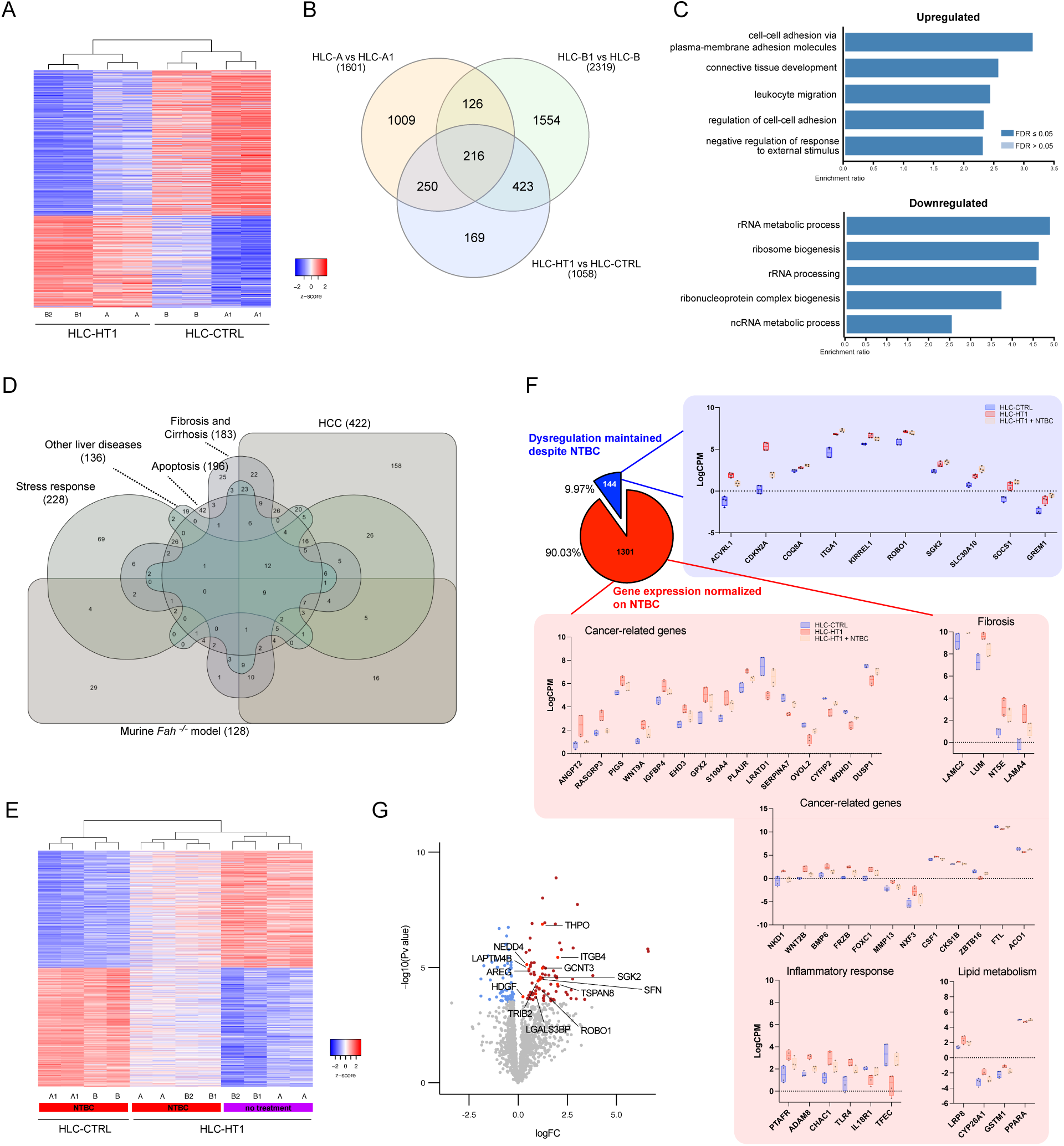
**A)** Heatmap showing the top 1,000 DEG between HLC-HT1 and HLC-CTRL (RNA-seq, unsupervised clustering). **B)** Importance of using isogeneic controls: when the transcriptome of HLC-HT1 was compared to their isogeneic HLC-CTRL’s, only 216 genes were commonly dysregulated between different genetic backgrounds. **C)** Enrichment analysis showing the top 5 significantly altered biological pathways in HLC-HT1 compared to HLC-CTRL (gene ontology with over representation analysis). **D)** Shared DEG among different liver-specific pathological pathways involved in HT1 pathophysiology (see Supp. Table 2 for references). **E)** Transcriptomic profile of HLC-HT1 moved closer to HLC-CTRL upon treatment with 50 µM NTBC (RNA-seq, top 1,000 DGE, unsupervised clustering). **F)** Most genes dysregulated in HLC-HT1 were normalized upon treatment with 50 µM NTBC, whereas 144 remained dysregulated (individual gene expression profiles in HLC-CTRL compared to HLC-HT1 with or without NTBC treatment). **G)** Some oncogenes associated with HCC remained upregulated in HLC-HT1 despite NTBC treatment.

We performed gene ontology analysis to determine the most significant biological processes behind the 1,058 DEG identified between HLC-HT1 and HLC-CTRL. Reactome pathway data were incorporated, and redundancy reduction techniques were applied to refine the results. Upregulated genes resulted in playing a role in cell-cell adhesion, cell-cell junction organization, and connective tissue development. On the contrary, significantly downregulated processes included ribosomal RNA metabolism, ribosome biogenesis, processing and assembly, RNA-protein complex biogenesis and non-coding RNA metabolic processes (Fig. 6C).

Among genes differentially expressed in HLC-HT1, 672 have been previously associated with HT1 and other related liver conditions, including 128 genes dysregulated in HT1 mouse models, 183 dysregulated in liver fibrosis and cirrhosis, 196 in apoptosis, 422 in hepatocellular carcinoma, 228 in stress response, and 136 genes in other liver diseases (Fig 6D, Supp. Table 2). A transcriptomic comparison between significantly altered genes in the HT1 *Fah^-/^*^-^ mouse model (464 genes)^33^ and our *in vitro* model (1,058 genes) identified 14 shared genes, all involved in processes that are crucial for HT1 pathophysiology (Table 2).

**Table 2.**
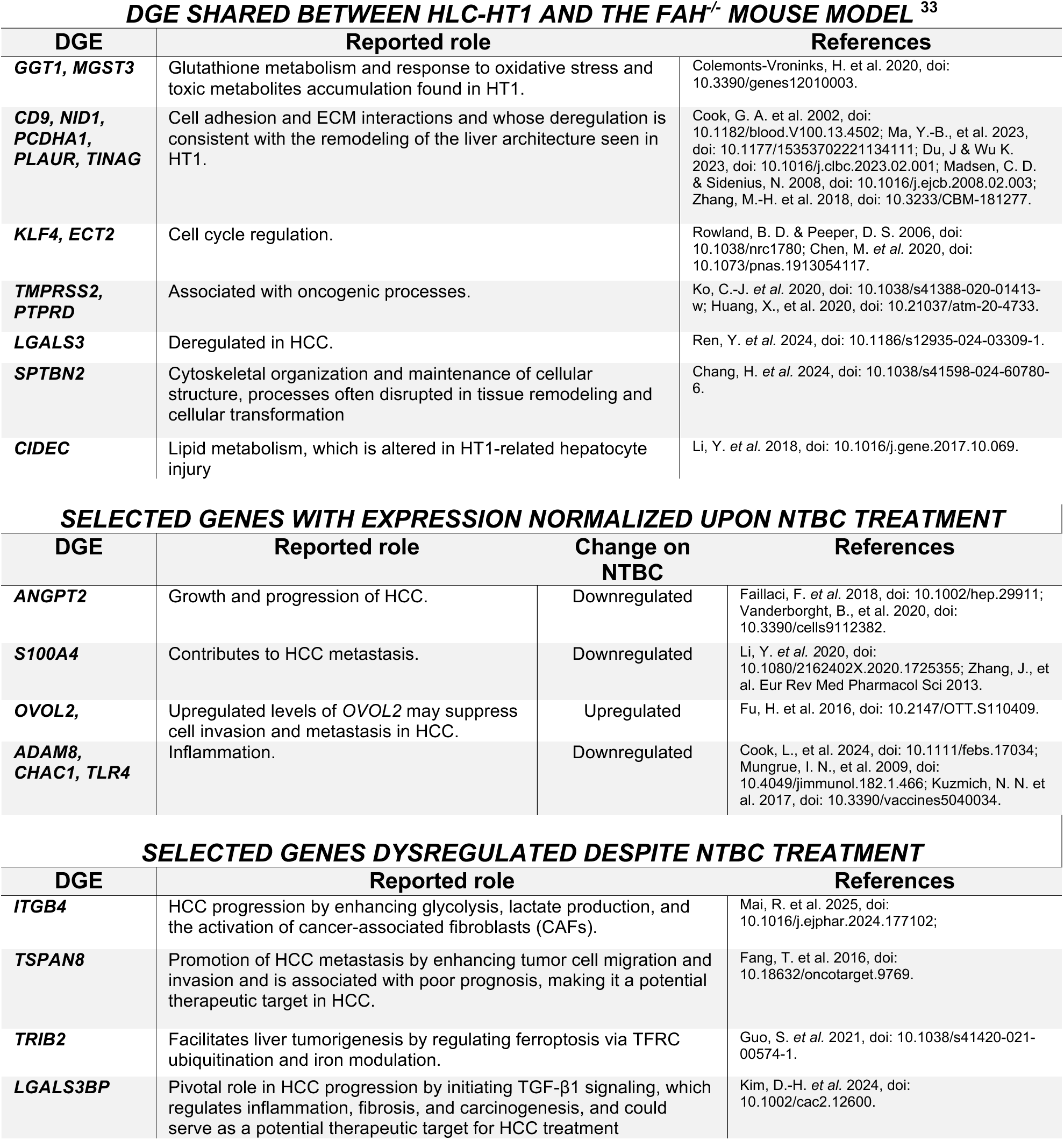
*FAH* deficiency in HLC-HT1 dysregulated the expression of genes involved in processes compatible with HT1 clinical presentation. NTBC reversed the expression of most of these genes, but not all.

### NTBC treatment corrects most but not all dysregulated pathways in HT1 hepatocytes

In order to assess the effect of NTBC on HT1 hepatocytes, we studied the transcriptomic profile of HLC-HT1 upon 50 µM NTBC treatment and compared it to both NTBC-treated healthy HLC-CTRL and untreated HLC-HT1. Overall, upon NTBC supplementation, the gene expression profile of HLC-HT1 became more similar to HLC-CTRL’s (Fig. 6E, supp Fig. 3A). Indeed, among the 1445 DEG in HLC-HT1, 1301 (90.03%) normalized their expression upon treatment with NTBC, going back to the levels observed in HLC-CTRL. Among dysregulated genes that normalized upon NTBC treatment were *ANGPT2, S100A4, OVOL2, ADAM8*, *CHAC1*, and *TLR4*, which are predominantly linked to cancer-related pathways, fibrosis, inflammatory responses, and lipid metabolism (Fig. 6F and Table 2). However, the expression of 144 genes (9.97%) remained significantly dysregulated (85 upregulated and 59 downregulated) despite NTBC treatment (Fig 6G and Supp. Table 3). We compared such 144 dysregulated genes with the 216 DEG mentioned above and identified 32 shared genes, among which four notable oncogenes (*ITGB4, TSPAN8, TRIB2*, and *LGALS3BP*), that could play a significant role in HT1-related HCC development and progression (Table 2).

## Discussion

We previously developed robust and reliable processes to differentiate human iPSC into HLC. Our protocol results in significantly improved hepatic functions with high homogeneity and yield, which makes it an effective tool for studying various liver diseases *in vitro.*^30^ By using this protocol, here we describe a first, *in vitro,* human model of HT1 that recapitulates the disease phenotype. We demonstrated that our novel model recreates the main features of HT1 liver disease under controlled experimental conditions, with acute and severe hepatocellular injury, SA accumulation and upregulation of genes involved in liver inflammation and fibrosis. The dysregulated genes and pathways identified in this human model align well with HT1 clinical phenotype and reflect what was previously shown in the *Fah*^-/-^ mouse model.^33^ The transcriptomic profile of HLC-HT1 shares significant similarities with several human pathophysiological processes involved in HT1, such as HCC, liver fibrosis and cirrhosis, apoptosis, and cellular responses to stress (Fig. 6D and Supp. Table 1). Interestingly, despite the limitation of a very short exposure to toxic tyrosine metabolites, our human HT1-HLC model showed results more representative of such processes than the murine *Fah^-/-^* model (Fig. 6D).

NTBC, an HPD inhibitor, is the mainstay of HT1 treatment. Nevertheless, NTBC was never assessed through randomized controlled clinical trials and neither the minimum effective nor the maximum tolerated doses are known despite NTBC being used in the clinic for more than 30 years. Based on only one *in vitro* study on purified human HPD, consensus recommendations suggested targeting NTBC trough plasma levels between 40 and 60 µM.^34,35^ Trough plasma concentrations within such a range were shown to be associated with normal SA levels.^36^ Based on extensive clinical experience, our center has suggested levels ≥50 µM to optimize control of the disease.^32^ With this study, we show that NTBC plasma levels of at least 50 µM are required to eliminate hepatocellular toxicity and effectively maintain SA within upper limits of normal. We validated such a threshold on retrospective clinical data collected over 4 years in a large cohort of HT1 patients followed in our center. As shown above, despite the limitation of random annual blood sampling and no control over therapeutic adherence, patients with NTBC trough levels ≥50 µM presented fewer episodes of elevated plasma ALT and showed lower plasma and urinary SA. Our *in vitro* model showed that a lower concentration of NTBC (20 µM), despite being a trough level that is often tolerated in HT1 patients, was ineffective to suppress SA formation and prevent hepatocellular death. This well aligns with previous observations in clinical practice by other groups.^37–39^ Interestingly, a dose of NTBC above the recommended therapeutic range (100 µM) appeared toxic on liver cells in culture, independently of whether the cells expressed a functional FAH enzyme (Fig. 4D).

In this study we only assessed two genetic backgrounds, which represents a theoretical limitation. Nevertheless, thanks to the use of isogeneic controls, we were able to evaluate the effect of *FAH* mutation on the transcriptome of human liver cells free of the confounding effects due to interindividual genetic variability. This controls better for genetic background than do comparisons between affected and unaffected individuals, providing a clearer picture of the involved signaling pathways and pathophysiological processes and reducing the need for large samples or cohorts when studying the mechanisms of rare diseases such as HT1.

Our study indicates that most genes dysregulated because of *FAH* mutation showed normalized expression upon treatment with an appropriate dose of NTBC (Fig. 6F). These normalized genes are involved in several pathophysiological processes, including fibrosis, inflammatory response, lipid metabolism, and tumorigenesis, specifically HCC (Table 2). These findings highlight the therapeutic impact of NTBC on the critical pathological processes of HT1 and underscore the potential of our *in vitro* model to assess the effect of treatments on the mechanisms of human disease.

Although most genes dysregulated by HT1 normalized on NTBC, the expression of almost 10% of them remained significantly altered. We explored genes whose expression remained significantly dysregulated despite NTBC treatment and identified 32 DEG of particular interest. Several of such genes are known to contribute to HCC by different mechanisms, are associated with poor prognosis, tumor progression and resistance to apoptosis, and could represent potential targets for further investigation.

Overall, our findings suggest that while NTBC treatment effectively normalizes the expression of most genes involved in the processes leading to the phenotype and complications of HT1, there are still significant dysregulations in some biological pathways that may pose a long-term risk for HT1 patients despite optimal therapy. Long-term liver cancer development is a known concern in the *Fah^-/-^* mouse model,^16,40^ but to date, no HCC has ever been documented in patients having started NTBC early in life. Our model does not allow for assessing whether such gene dysregulation uncorrected by NTBC persists over time. Nevertheless, these observations in a human model confirm previous findings in rodents and suggest that the possibility of long-term HCC development despite NTBC treatment should not be dismissed, warranting continued patient follow-up and further research to clarify the role of identified pathways in the tyrosinemic liver.

In conclusion, our study demonstrates that our iPSC-derived hepatocytes and isogeneic controls faithfully replicate HT1 phenotype, providing a valuable platform to investigate the disease pathophysiology and the effects of treatment in a representative human model under controlled conditions. This allowed us to confirm that NTBC is effective in preventing hepatocellular damage, and to robustly determine the minimal dose required to avert cellular injury, which was further validated in the world largest clinical cohort of HT1 patients. Moreover, this study described the genes and processes involved in HT1 pathophysiology and the effects of treatment on such genes and provided interesting clues to suggest a potential long-term oncogenic risk for patients on NTBC. Overall, our findings highlight the growing promise of such a representative *in vitro* disease modeling approach for studying metabolic liver diseases and assessing the effect of already available and new treatments. More importantly, they offer compelling evidence to establish minimal therapeutic levels of NTBC in HT1 patients.

## Online Methods

### Ethical approval

This project was approved by the CHU Sainte-Justine internal review board (2016-1134) and conducted according to the ethical guidelines of the 1975 Declaration of Helsinki.

### Human iPSC lines

We generated human iPSCs from a healthy female individual (iPSC-B) and a male HT1 patient with proven FAH-deficiency (iPSC-A). Sequencing of the *FAH* gene confirmed that iPSC-B had wild-type genotype whereas iPSC-A had a compound mutation with one allele carrying the p.W273R mutation and the second allele carrying an intronic deletion of 8,200 bp between exons 13 and 14. iPSC were reprogrammed from CD34-enriched peripheral blood mononuclear cells (iPSC-A) or skin fibroblasts (iPSC-B) CytoTune-iPS 2.0 Sendai reprogramming kit (ThermoFisher Scientific; A16517) according to the manufacturer’s instructions. The colonies with suggestive morphology were manually picked as we previously described, based on live staining for the pluripotency marker TRA-1-60 and then cultured as iPSC thereafter, with single colony sub-cloning for the first 5 passages.^30^ After passage 10, a temperature shift (incubation at 39°C for 72h) was performed to remove the cMyc gene, and the absence of Sendai virus was confirmed by RT-qPCR. PSC cultures were maintained on human recombinant vitronectin-coated plates (ThermoFisher Scientific), in Essential 8 Flex medium (ThermoFisher Scientific), at 37°C in humidified atmosphere containing 5% CO2 and 4% O2. Obtained iPSC populations were extensively characterized by qPCR, FACS, immunostaining and RNA-seq as previously described,^30^ information regarding material used for qPCR, flow cytometry and immunostaining can be found in supplementary tables (Supp. Table 2-5).

### Genome editing

FAH-deficient iPSC clones were generated from iPSC-B by CRISPR/eSpCas9-based *FAH* gene knockout, while the disease-causing mutation in patient-derived iPSC-A was corrected by CRISPR-Cas9^D10A^ nickase-based, homology-directed repair to create an isogeneic control, as we previously described.^31^

### DNA Sequencing

DNA was isolated as previously described.^31^ Sanger sequencing was performed by the Université de Montréal’s Institute for Research in Immunology and Cancer (IRIC) with the 48-capillary 3730 DNA Analyzer (Applied Biosystem; ABI 3730).

### iPSC differentiation into hepatocyte-like cells (HLC)

iPSC differentiation into HLC was conducted as described previously by our group.^30^ HLCs were extensively characterized by qPCR, flow cytometry, immunostaining, and functional assessment (see below), as we previously described.^30^ Information regarding material used for differentiation, qPCR, flow cytometry and immunostaining can be found in Supplementary Tables 4-7,9. HLC-HT1 received 50 µM of Nitisinone/NTBC (LGC; TRC-N490135) from day 16 until the end of differentiation. Gene expression of tyrosine metabolism pathway enzymes was quantified by qPCR using TaqMan assay (assay IDs listed in Supp. Table 4) as previously described.^30^ Albumin secretion: Albumin secretion by HLCs (day 30 of differentiation) were measured by Human Albumin ELISA Kit (Abcam, ab179887) according to manufacturers’ instruction.

### Cy3A4 Activity

Cyp3A4 activity in HLCs (day 30 of differentiation) was measured using P450-Glo Assay (Promega, V9002), according to manufacturers’ instructions.

### Western blot

HLCs (day 30 of differentiation) were prepared in RIPA buffer supplemented with 1 mM PMSF and protease and phosphatase inhibitor cocktails (Sigma; P8340). The protein concentration was measured using UV spectrophotometry (Thermo Scientific™ NanoDrop™ Eight; 13400533) and 10 µg of protein were loaded on 10% sodium dodecyl sulphate-polyacrylamide gel, and electrophoresed, then transferred on polyvinylidene difluoride membranes. Following the transfer, the membranes were blocked with 5% fat-free milk and PBS-T for one hour at room temperature (22°C). Membranes were then incubated in (1:1000) dilution of primary antibodies of FAH and beta actin (Supp. Table 8) overnight at 4°C. After, membranes were washed 5 times with PBS-T buffer and incubated with their corresponding secondary antibody coupled to horseradish peroxidase (1:10,000) for one hour at room temperature (22°C). After a few washes, membranes were exposed to chemiluminescence reagent (Clarity Max Western ECL Substrate, Bio-Rad Laboratories, USA, 1705062) and probed on the Chemidoc^TM^.

### Apoptosis assay

For all experiments, HLC-HT1 were supplemented with 50 µM NTBC starting from day 16 of the differentiation protocol to avoid apoptosis. To assess the effect of FAH deficiency on apoptosis, on day 30 of differentiation, NTBC was withdrawn, the cells were washed, and NTBC-free, stage-appropriate differentiation media was added. Both HLC-HT1 and HLC-CTRL cells were then supplemented with either 300 µM L-Tyrosine (Sigma; 93829), 4-Hydroxyphenylpyruvic acid (Sigma 114286; a kind gift of Dr. Grant Mitchell), or Homogentisic acid (Sigma H0751; a kind gift of Dr. Grant Mitchell). The Incucyte® Caspase-3/7 Dye for Apoptosis (Sartorius; 4440) was added to the media according to the manufacturer’s instructions, and apoptosis was monitored for 24 hours using the Incucyte® S3 Live Cell Analysis Instrument (Sartorius) and the data were analyzed with Incucyte® 2022B Rev2 GUI software. In a separate experiment, HLC-CTRL cells on day 30 of differentiation were treated with L-Tyrosine (300 and 600 µM) or NTBC (20, 50, and 100 µM). The percentage of apoptotic cells was identified using the same technique. In another experiment on day 30 of differentiation, HLC-HT1 and HLC-CTRL cells were treated with 300 µM L-Tyrosine and varying doses of NTBC (0, 20, 50, and 100 µM). NucBlue™ Live ReadyProbes™ Reagent (ThermoFisher Scientific; R37605) and CellEvent™ Caspase-3/7 Detection Reagents (ThermoFisher Scientific; C10423) were added, and images were captured every two hours using the EVOS FL Colour Imaging System (Life Technologies; AMF 4300) for 24 hours. For each experiment, three replicates were performed, and five images per replicate were analyzed using ImageJ (Fiji).

### Plasma biochemistry

Plasma ALT levels of 99 HT1 patients were measured by the CHU Sainte-Justine clinical laboratory with Health Canada-certified methods.

### Succinylacetone (SA) measurement

All succinylacetone assays were performed in the Biochemical Genetics Laboratory, within the Medical Genetics Service, at the Sherbrooke University Hospital Centre (CHUS). SA levels in cell culture supernatants and in patients’ plasma and urine specimens were each quantified by a sensitive method using gas chromatography–mass spectrometry (GC-MS), essentially as previously described.^41^ A stable isotope dilution process was used, with sample treatment consisting of an oximation step followed by a liquid-liquid extraction process then trimethylsilyl derivatization. GC-MS analysis was then performed as follows. An Agilent GC-MS system was used consisting of a 6890N model gas chromatograph (GC) equipped with a fused-silica capillary column coated with 5% phenyl-95% dimethylpolysiloxane (ZB-5, Phenomenex, 30 m x 0.25 mm i.d., 0.25 mm film thickness) and a 5973 model inert mass selective detector. The GC/MS was employed with helium as the carrier gas at a constant flow of 1 mL/min. The oven temperature started at 80 °C and remained at this temperature for 1 min increasing to 280°C at 17°C/min ramp rate and then holding at 280°C for 2.5 min. A 3 µL specimen volume was injected in the injection port adjusted to 250°C, using a splitless injection mode. The MS transfer line was at 280°C and the ion source temperature at 230°C. The mass spectrometer was operated at 70 eV in the electron impact mode with selected ion monitoring (SIM). The selected ion groups used for identification of succinylacetone (SA) and the internal standard ^13^C5-SA in SIM mode were *m*/*z* 620 and *m*/*z* 625, respectively. Dwell time for each ion was set at 100 ms. The assay limit of detection was 1 nmol/L and the lower limit of quantitation was 3 nmol/L. The reference limits (upper limits of normal) were 24 nmol/L for plasma and 34 µmol/mol creatinine for urine.

### Nitisinone (NTBC) measurent

Nitisinone (NTBC) (purity ≥ 95%) and clonazepam (purity ≥ 99.9%, 1 mg/ml in methanol), used as internal standards, were supplied by Sigma–Aldrich (North York, Ontario, Canada) and Cerilliant (Round Rock, Texas, USA), respectively. General reagents (sodium phosphate monobasic and sodium chloride) and solvents (methanol and acetonitrile) of the purest available quality were supplied by Fisher Scientific (Montreal, Quebec, Canada). Ultrapure water (18 MΩ) was produced by a Millipore Synthesis A-10 water purification system (Etobicoke, Ontario, Canada). NTBC was dissolved in acetonitrile to create two stock solutions of 1.00 mg/mL (3.04 mM). One was used to form calibration curve standards, and the other was used to prepare Quality Control (QC) samples. Physiological saline was subsequently used to dilute the stock solutions 2-fold, which were then further diluted 5-fold with human serum (MSG4000, Golden West Biologicals Inc., Temecula, CA 92590, USA) to create an NTBC working solution of 100 ng/ml (304 µM). NTBC calibration standards of 2.37, 4.75, 9.5, 19, 38, 76, and 152 µM in human plasma were prepared via serial dilutions. Pooled plasma without added NTBC was also included in the set of calibrators. QC samples were made to yield final concentrations of 30.4 and 60.8 µM. All stock solutions and pools of QC samples were stored at −80°C. The analysis was performed using the 1260 Infinity liquid chromatography system (Agilent Technologies Inc., Montreal, Canada), with OpenLab CDS version C.01.04 software. The system operates at room temperature and consists of a high-performance degasser (G4225A), a binary pump (G1312B), an autosampler (G1329B), a thermostatted column compartment (G1316A), and a diode array detector (G1315D) set at 278 nm. Chromatographic separation was performed on a Zorbax Eclipse XDB-C8 column (4.6 x 150 mm) with particle and pore sizes of 5 µm and 80 Å, respectively (Agilent Technologies Inc., Montreal, Canada). The column was protected by a 0.5 µm stainless steel disk and seal (Supelco, St. Louis, Missouri, USA). The mobile phase was prepared using a phosphate buffer (0.1M, pH 2.0) and acetonitrile in a 50:50 (v/v) ratio. The mobile phase was pumped at a flow rate of 1.7 mL/min in an isocratic mode and at a pressure of around 130 bars. Prior to processing, plasma samples were thawed at 20–25°C, and a 100 μL aliquot was transferred to a 1.5 mL conical container. A volume of 200 μL of acetonitrile containing the internal standard (clonazepam at a concentration of 4 µg/ml) was added to the samples to precipitate proteins and extract analytes from the biomatrix. Smaller volumes of plasma could be analyzed, but it was necessary to maintain the matrix–protein precipitation solution ratio. The samples were then vortex-mixed and centrifuged (4 min at 20°C and 16,000 g). The supernatant was transferred to 1.5 mL conical tubes and evaporated to dryness under a nitrogen stream at 37°C (Biotage Zymark TurboVap LV, Boston/Salem, New Hampshire, USA). Then, 100 µL of mobile phase was added to the tube, vortex-mixed for 10 seconds, and centrifuged for 2 minutes at 20°C and 16,000 g. The clear supernatant was transferred to a 200 µL glass tube, which was then loaded onto the autosampler before injection of 25 µL onto the column. Retention times of clonazepam and NTBC were around 1.8 and 5.0 min, respectively. Stop time was set at 7 min. Measured data were subjected to linear regression analysis of the NTBC to clonazepam peak area ratio against the concentration (µM). The linearity of the method was found to be in the range of 2.37 to 152 µM, which is suitable for NTBC measurement in the plasma of treated tyrosinemic patients (see a representative calibration curve in Figure below). The lower limit of quantification (2.37 µM) was adapted to expected NTBC plasma concentrations and was at least 10 times the signal of a blank sample. The method has been validated according to the latest US Food and Drug Administration and European Medicines Agency guidelines for its precision (concentration ranging from 2.7 to 152 µM: 101.7 ± 2.4%), selectivity (absence of spectral interference in blank plasma as well as in samples of patients at the retention time of NTBC and clonazepam), carry-over (0.7 ± 3.6%), matrix effect (0.8 ± 0.7%), stability 7 hours post-extraction (ratio difference: −1.8 ± 1.3%), and recovery (NTBC: 82.2 ± 2.7% and clonazepam: 82.5 ± 2.3%). An alternative proficiency testing procedure that fulfills CLIA and Standard Council of Canada requirements for the alternative assessment of non-commercially available analytes is used.

### RNA sequencing

Differentiation of HLCs from iPSCs was carried out until day 30 as described above, with 50 µM NTBC supplementation from day 16 onwards. HLCs were then harvested after a 24-hour NTBC washout to capture transcriptomic changes representative of the disease state. Total RNA was extracted as previously described,^30^ and then quantified (NanoDrop 8000, ThermoFisher Scientific), and quality-controlled (BioAnalyzer 2100, Agilent). Five hundred ng of total RNA were used as input for the TruSeq RNA v2 mRNA Sample Prep Kit (Illumina) and the KAPA mRNA Hyper Prep kit (Roche), and sequencing libraries were generated according to the manufacturers’ protocol. The libraries were then pooled and sequenced on the HiSeq 2500 (Illumina) and NextSeq 500 (Illumina) instruments as per the manufacturer’s instructions, generating single-end reads with a length of 100 bp and 75 bp, respectively. Following sequencing, HISAT2 was used for alignment and featureCounts to obtain the raw counts.^42,43^ Differential gene expression analysis was conducted using edgeR-Limma and the DESeq2 package.^44–46^ Significance thresholds were set at p ≤ 0.05 and log2 fold change ≥1. Enrichment analyses of the top 500 differentially expressed genes were performed using Enrichr and WebGestalt 2019, focusing on Gene Ontology biological processes and Reactome pathways.^47,48^ Results were visualized through volcano plots, z-score heatmaps, and logCPM expression plots. The heatmap shown in Supplementary Figure 1 was performed using the Biojupies platform.^49^ Sequencing data of connected pathophysiological mechanisms shown in Fig. 6D were obtained from the Gene Expression Omnibus public repository (NIH). RNAseq data are available on the Gene Expression Omnibus (GEO) repository (GSE301101).

### Statistical analysis

Unpaired t test, Mann-Whitney test, Welch’s test, Wilcoxon test, Pearson r, Chi-square were used as appropriate. Values of *p*<0.05 were considered statistically significant. Statistical analysis was done using GraphPad Prism 8 (GraphPad Software, La Jolla, CA, USA).

## Impact and implications

This study established the first representative *in vitro* human model of HT1 using patient-derived iPSC and isogeneic controls, providing a crucial platform for understanding the disease and evaluating treatments. We demonstrated that an appropriate therapeutic dose of NTBC, which we validated in the largest clinical cohort available, effectively reduces metabolic dysfunction and normalizes most gene dysregulation in HT1. Nevertheless, some oncogenic pathways remain unaffected by NTBC treatment, highlighting the need for continued follow-up and additional studies to fully address the disease’s long-term risks and complexities.

## Highlights

- We describe an *in vitro* human model of HT1 that faithfully recapitulates the disease and enables research on its pathophysiology and treatment.
- An adequate dose of NTBC (50 µmol/L) effectively prevents the accumulation of succinylacetone and hepatocellular injury in FAH-deficient hepatocytes, normalizing over 90% of dysregulated genes.
- We validate the identified dose in the world largest cohort of HT1 patients, suggesting therapeutic thresholds for a treatment never tested in clinical trials.
- Some oncogenic pathways remain unaffected by NTBC treatment in the short term, which supports a need for long-term follow-up and surveillance.

## Author contribution

QT.P.: study concept and design, acquisition, analysis, and interpretation of data, statistical analysis, drafting of the manuscript. F.T.: analysis, and interpretation of data, statistical analysis, drafting of the manuscript. MA.M.: acquisition of data. E.B.: acquisition, analysis, and interpretation of data. R.M.: acquisition of data. Y.T.: acquisition, analysis, and interpretation of data, critical revision of the manuscript. G.M. and NTBC S.G.: clinical care of the studied cohort of patients. D.C.: acquisition and analysis of data. PJ.W.: analysis, and interpretation of data, critical revision of the manuscript. Y.D.: study concept and design, analysis, and interpretation of data, critical revision of the manuscript. U.H.: study concept and design, acquisition, analysis, and interpretation of data, critical revision of the manuscript. C.R.: study concept and design, analysis, and interpretation of data, critical revision of the manuscript. M.P.: study concept and design, analysis, and interpretation of data, drafting of the manuscript, critical revision of the manuscript, study supervision, obtained funding.

## Data availability

The data that supports the findings of this study are available from the corresponding author, M.P., upon request. RNAseq data are available on the Gene Expression Omnibus repository (GSE301101).

## Abbreviations

4-HPP: 4-hydroxyphenylpyruvate
ALT: alanine aminotransferase
DEG: differentially expressed genes
ECM: extracellular matrix
FAA: fumarylacetoacetate
FAH: fumarylacetoacetate hydrolase
HCC: hepatocellular carcinoma
HGA: homogentisic acid
HLC: hepatocyte-like cells
HPD: 4-hydroxyphenylpyruvate dioxygenase
HT1: hereditary type 1 tyrosinemia
iPSC: induced pluripotent cells
LT: liver transplant
MAA: maleylacetoacetate
NTBC: nitisinone
PHH: primary human hepatocytes
SA: succinylacetone
TAT: tyrosine aminotransferase
ULN: upper limit of normal

## Supplementary Figures

**Supplementary Figure 1.**
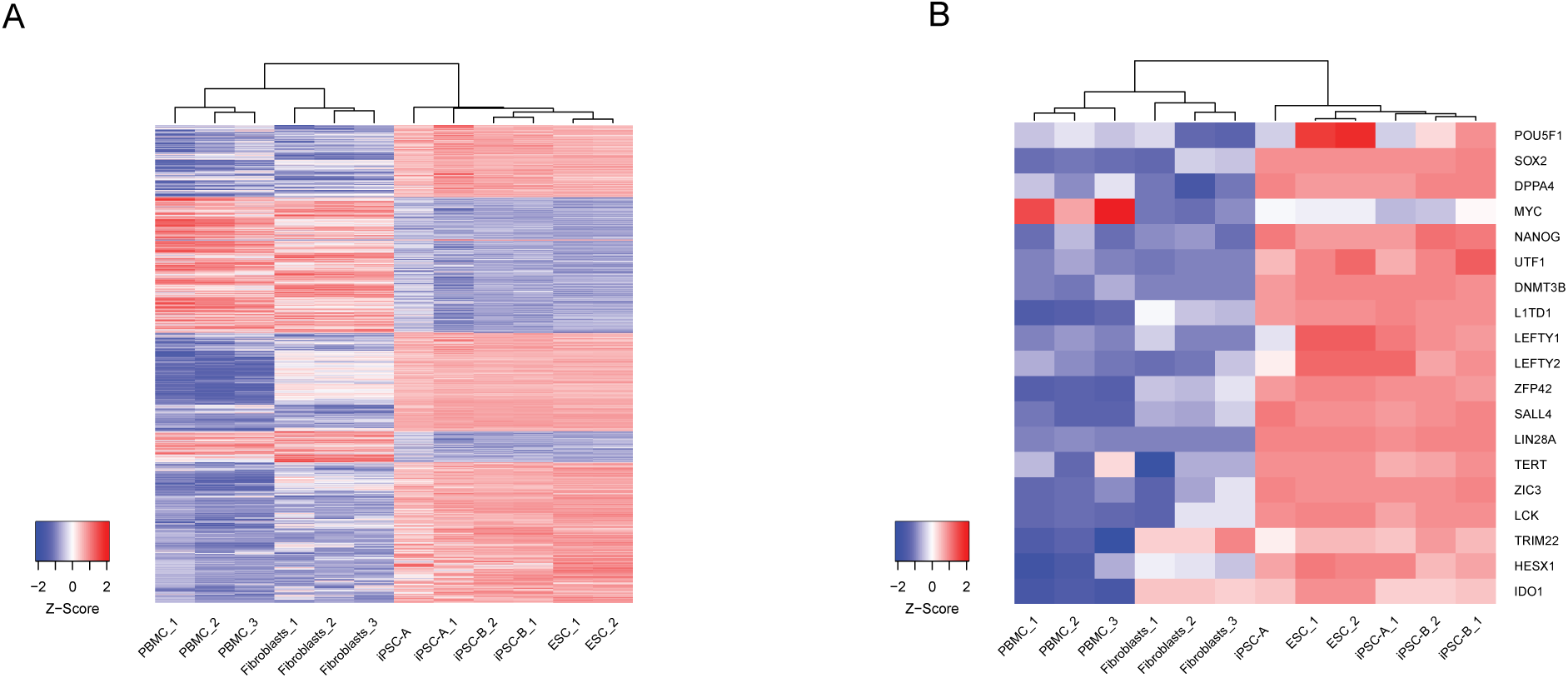
**A)** Heatmap of the top 1,000 differentially expressed genes shows that the transcriptomic profiles of iPSC lines closely match those of embryonic stem cells (ESC; RNA-seq, unsupervised clustering). **B)** Heatmap of pluripotency markers showing that the expression of pluripotency markers across the 6 populations was comparable and unaffected by genome editing (RNA-seq).

**Supplementary Figure 2.**
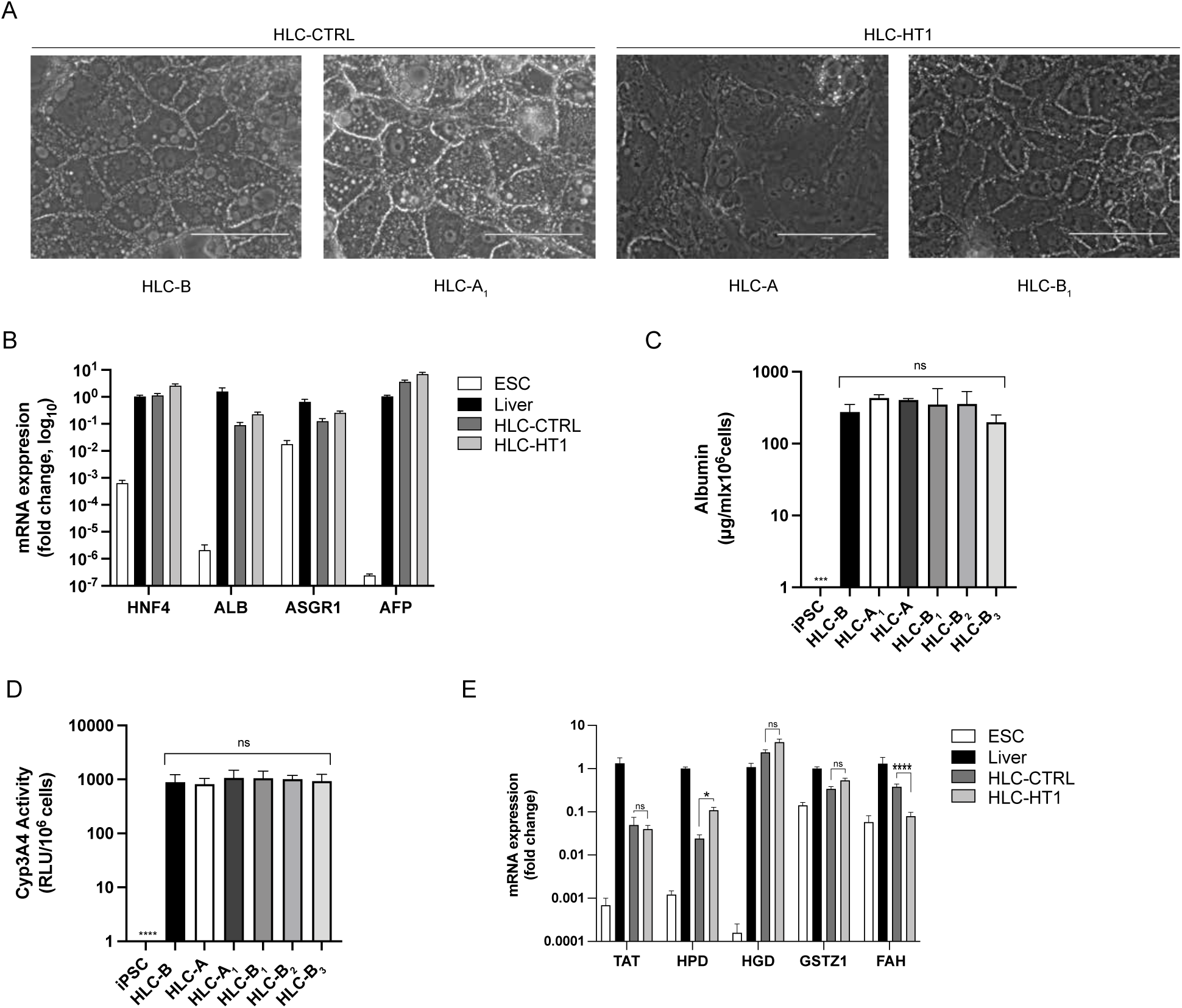
**A)** The morphology of mature HLC-HT1 and HLC-CTRL was identical, with no discernible differences related to FAH deficiency (phase contrast, scale bar = 100 µm, representative images). **B)** No difference was observed in the expression of *HNF4*, *ALB*, *ASGR1*, and *AFP* between HLC-HT1- and HLC-CTRL (RT-qPCR, N=3, mean ± SEM, unpaired t-test). **C)** All HLC lines (HT1 and CTRL) secreted albumin, with no significant differences observed between them (ELISA, N=3, mean ± SEM, unpaired t-test). **D)** The Cyp3A4 activity was detectable in all HLC lines (HT1 and CTRL), with no significant differences observed between them. **E)** All enzymes involved in tyrosine pathway were expressed in both HLC-HT1 and HLC-CTRL, except for FAH which was significantly less expressed in HLC-HT1 (RT-qPCR, N=3, mean ± SEM, fold change to expression in the healthy liver tissue, *p<0.05 and ****p<0.0001, multiple unpaired t-tests).

**Supplementary Figure 3.**
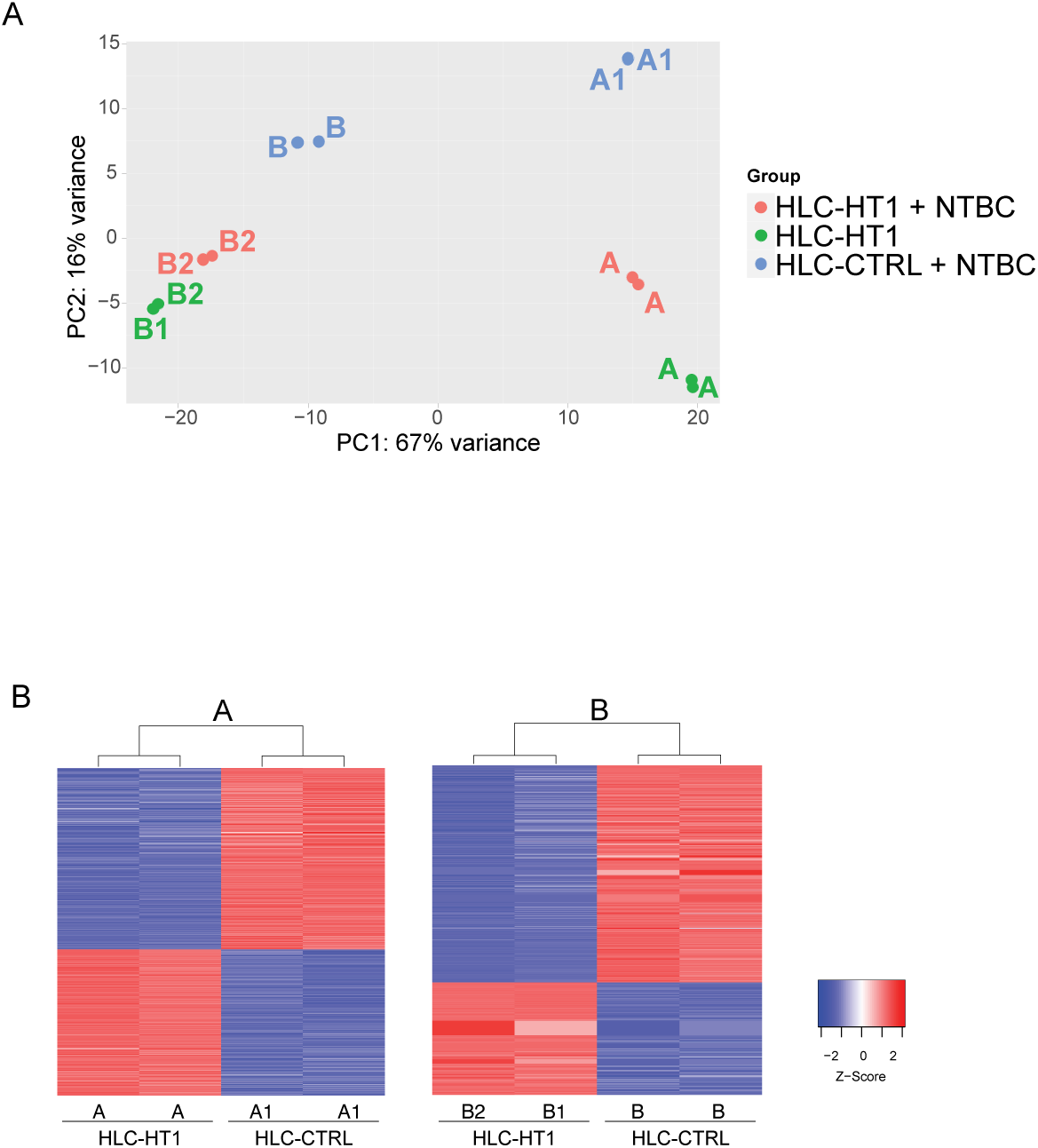
**A)** Principal component analysis (PCA) shows not only the differences in gene expression related to FAH deficiency, but also the impact of inter-donor variability (RNA-seq). **B)** Heatmaps showing the top 1,000 differentially expressed genes between clones derived from different donors and their isogenic controls (RNA-seq, unsupervised clustering).

## Supplementary Tables

See Supplementary Material.

## References

1. Larochelle, J. et al. [Hereditary tyrosinemia. I. Clinical and biological study of 62 cases]. Pediatrie 28, 5–18 (1973).

2. Larochelle, J. et al. Experience with 37 infants with tyrosinemia. Can. Med. Assoc. J. 97, 1051–1054 (1967).

3. Grompe, M. et al. A single mutation of the fumarylacetoacetate hydrolase gene in French Canadians with hereditary tyrosinemia type I. N. Engl. J. Med. 331, 353–357 (1994).

4. Lindblad, B., Lindstedt, S. & Steen, G. On the enzymic defects in hereditary tyrosinemia. Proc. Natl. Acad. Sci. U. S. A. 74, 4641–4645 (1977).

5. Russo, P. A., Mitchell, G. A. & Tanguay, R. M. Tyrosinemia: a review. Pediatr. Dev. Pathol. Off. J. Soc. Pediatr. Pathol. Paediatr. Pathol. Soc. 4, 212–221 (2001).

6. van Spronsen, F. J. et al. Hereditary tyrosinemia type I: a new clinical classification with difference in prognosis on dietary treatment. Hepatol. Baltim. Md 20, 1187–1191 (1994).

7. Bliksrud, Y. T., Ellingsen, A. & Bjørås, M. Fumarylacetoacetate inhibits the initial step of the base excision repair pathway: implication for the pathogenesis of tyrosinemia type I. J. Inherit. Metab. Dis. 36, 773–778 (2013).

8. Grompe, M. et al. Pharmacological correction of neonatal lethal hepatic dysfunction in a murine model of hereditary tyrosinaemia type I. Nat. Genet. 10, 453–460 (1995).

9. Tanguay, R. M., Angileri, F. & Vogel, A. Molecular Pathogenesis of Liver Injury in Hereditary Tyrosinemia 1. Adv. Exp. Med. Biol. 959, 49–64 (2017).

10. Paradis, K. et al. Liver transplantation for hereditary tyrosinemia: the Quebec experience. Am. J. Hum. Genet. 47, 338–342 (1990).

11. Lindstedt, S., Holme, E., Lock, E. A., Hjalmarson, O. & Strandvik, B. Treatment of hereditary tyrosinaemia type I by inhibition of 4-hydroxyphenylpyruvate dioxygenase. Lancet Lond. Engl. 340, 813–817 (1992).

12. Ellis, M. K. et al. Inhibition of 4-hydroxyphenylpyruvate dioxygenase by 2-(2-nitro-4-trifluoromethylbenzoyl)-cyclohexane-1,3-dione and 2-(2-chloro-4-methanesulfonylbenzoyl)-cyclohexane-1,3-dione. Toxicol. Appl. Pharmacol. 133, 12–19 (1995).

13. Lock, E. A. et al. Tissue distribution of 2-(2-nitro-4-trifluoromethylbenzoyl)cyclohexane-1-3-dione (NTBC): effect on enzymes involved in tyrosine catabolism and relevance to ocular toxicity in the rat. Toxicol. Appl. Pharmacol. 141, 439–447 (1996).

14. Lock, E. A., Gaskin, P., Ellis, M., Provan, W. M. & Smith, L. L. Tyrosinemia produced by 2-(2-nitro-4-trifluoromethylbenzoyl)-cyclohexane-1,3-dione (NTBC) in experimental animals and its relationship to corneal injury. Toxicol. Appl. Pharmacol. 215, 9–16 (2006).

15. Larochelle, J. et al. Effect of nitisinone (NTBC) treatment on the clinical course of hepatorenal tyrosinemia in Québec. Mol. Genet. Metab. 107, 49–54 (2012).

16. Al-Dhalimy, M., Overturf, K., Finegold, M. & Grompe, M. Long-term therapy with NTBC and tyrosine-restricted diet in a murine model of hereditary tyrosinemia type I. Mol. Genet. Metab. 75, 38–45 (2002).

17. Hall, M. G., Wilks, M. F., Provan, W. M., Eksborg, S. & Lumholtz, B. Pharmacokinetics and pharmacodynamics of NTBC (2-(2-nitro-4-fluoromethylbenzoyl)-1,3-cyclohexanedione) and mesotrione, inhibitors of 4-hydroxyphenyl pyruvate dioxygenase (HPPD) following a single dose to healthy male volunteers. Br. J. Clin. Pharmacol. 52, 169–177 (2001).

18. Mitchell, G. A. & Yang, H. Remaining Challenges in the Treatment of Tyrosinemia from the Clinician’s Viewpoint. Adv. Exp. Med. Biol. 959, 205–213 (2017).

19. Marhenke, S. et al. Activation of nuclear factor E2-related factor 2 in hereditary tyrosinemia type 1 and its role in survival and tumor development. Hepatol. Baltim. Md 48, 487–496 (2008).

20. Vogel, A., et al. Sustained Phosphorylation of Bid Is a Marker for Resistance to Fas-Induced Apoptosis During Chronic Liver Diseases. Gastroenterology 130, 104–119 (2006).

21. Paulk, N. K. et al. Adeno-associated virus gene repair corrects a mouse model of hereditary tyrosinemia in vivo. Hepatol. Baltim. Md 51, 1200–1208 (2010).

22. Yin, H. et al. Genome editing with Cas9 in adult mice corrects a disease mutation and phenotype. Nat. Biotechnol. 32, 551–553 (2014).

23. Stéphenne, X. et al. Cryopreservation of human hepatocytes alters the mitochondrial respiratory chain complex 1. Cell Transplant. 16, 409–419 (2007).

24. Zeilinger, K., Freyer, N., Damm, G., Seehofer, D. & Knöspel, F. Cell sources for in vitro human liver cell culture models. Exp. Biol. Med. 241, 1684–1698 (2016).

25. den Braver-Sewradj, S. P., et al. Inter-donor variability of phase I/phase II metabolism of three reference drugs in cryopreserved primary human hepatocytes in suspension and monolayer. Toxicol. Vitro Int. J. Publ. Assoc. BIBRA 33, 71–79 (2016).

26. Paganelli, M. et al. Differentiated umbilical cord matrix stem cells as a new in vitro model to study early events during hepatitis B virus infection. Hepatol. Baltim. Md 57, 59–69 (2013).

27. Rashid, S. T. et al. Modeling inherited metabolic disorders of the liver using human induced pluripotent stem cells. J. Clin. Invest. 120, 3127–3136 (2010).

28. Segeritz, C.-P. et al. hiPSC hepatocyte model demonstrates the role of unfolded protein response and inflammatory networks in α1-antitrypsin deficiency. J. Hepatol. 69, 851–860 (2018).

29. Baxter, M. et al. Phenotypic and functional analyses show stem cell-derived hepatocyte-like cells better mimic fetal rather than adult hepatocytes. J. Hepatol. 62, 581–589 (2015).

30. Raggi, C. et al. Leveraging interacting signaling pathways to robustly improve the quality and yield of human pluripotent stem cell-derived hepatoblasts and hepatocytes. Stem Cell Rep. 17, 584–598 (2022).

31. Pham, Q. T. et al. High-throughput assessment of mutations generated by genome editing in induced pluripotent stem cells by high-resolution melting analysis. Cytotherapy 22, 536–542 (2020).

32. Alvarez, F. et al. The Québec NTBC Study. in Hereditary Tyrosinemia: Pathogenesis, Screening and Management (ed. Tanguay, R. M.) 187–195 (Springer International Publishing, Cham, 2017). doi:10.1007/978-3-319-55780-9_17.

33. Colemonts-Vroninks, H. et al. Oxidative Stress, Glutathione Metabolism, and Liver Regeneration Pathways Are Activated in Hereditary Tyrosinemia Type 1 Mice upon Short-Term Nitisinone Discontinuation. Genes 12, 3 (2020).

34. Lock, E. A. et al. From toxicological problem to therapeutic use: the discovery of the mode of action of 2-(2-nitro-4-trifluoromethylbenzoyl)-1,3-cyclohexanedione (NTBC), its toxicology and development as a drug. J. Inherit. Metab. Dis. 21, 498–506 (1998).

35. Chinsky, J. M. et al. Diagnosis and treatment of tyrosinemia type I: a US and Canadian consensus group review and recommendations. Genet. Med. Off. J. Am. Coll. Med. Genet. 19, (2017).

36. Fuenzalida, K. et al. NTBC Treatment Monitoring in Chilean Patients with Tyrosinemia Type 1 and Its Association with Biochemical Parameters and Liver Biomarkers. J. Clin. Med. 10, 5832 (2021).

37. Kienstra, N. S. et al. Daily variation of NTBC and its relation to succinylacetone in tyrosinemia type 1 patients comparing a single dose to two doses a day. J. Inherit. Metab. Dis. 41, 181–186 (2018).

38. El-Karaksy, H. et al. Clinical practice. NTBC therapy for tyrosinemia type 1: how much is enough? Eur. J. Pediatr. 169, 689–693 (2010).

39. Jack, R. M. & Scott, C. R. Validation of a therapeutic range for nitisinone in patients treated for tyrosinemia type 1 based on reduction of succinylacetone excretion. JIMD Rep. 46, 75–78 (2019).

40. Neuckermans, J. et al. Hereditary Tyrosinemia Type 1 Mice under Continuous Nitisinone Treatment Display Remnants of an Uncorrected Liver Disease Phenotype. Genes 14, 693 (2023).

41. Cyr D, et al. A GC/MS validated method for the nanomolar range determination of succinylacetone in amniotic fluid and plasma: An analytical tool for tyrosinemia type I. J Chromatogr B 2006;832:24–9.

42. Kim D, et al. Graph-based genome alignment and genotyping with HISAT2 and HISAT-genotype. Nat Biotechnol 2019;37:907–15.

43. Liao Y, et al. featureCounts: an efficient general purpose program for assigning sequence reads to genomic features. Bioinformatics 2014;30:923–30.

44. Love MI, et al. Moderated estimation of fold change and dispersion for RNA-seq data with DESeq2. Genome Biol 2014;15:550.

45. Robinson MD, et al. edgeR: a Bioconductor package for differential expression analysis of digital gene expression data. Bioinformatics 2009;26:139–40.

46. Ritchie ME, et al. limma powers differential expression analyses for RNA-sequencing and microarray studies. Nucleic Acids Res 2015;43:e47–e47.

47. Liao Y, et al. WebGestalt 2019: gene set analysis toolkit with revamped UIs and APIs. Nucleic Acids Res 2019;47:W199–205.

48. Kuleshov MV, et al. Enrichr: a comprehensive gene set enrichment analysis web server 2016 update. Nucleic Acids Res 2016;44:W90–7.

49. Torre D, et al. BioJupies: Automated Generation of Interactive Notebooks for RNA-Seq Data Analysis in the Cloud. Cell Syst 2018;7:556–561.e3.

