## Supplementary Tables for "NTBC dosing and outcomes in hereditary tyrosinemia type 1: insights from a representative human model and 99 patients"

**Supplementary Table 1.** List of differentially expressed genes in HLC-HT1 compared to HLC-CTRL shared between all comparison groups (Fig. 6B).

| **216 dysregulated genes shared between all comparison groups (Fig. 6B)** | | | | | |
| --- | --- | --- | --- | --- | --- |
| **ENTREZID** | **SYMBOL** | **logFC** | **AveExpr** | **P.Value** | **adj.P.Val** |
| 728927 | ZNF736 | 7.78325711 | 1.07584314 | 1.77E-13 | 2.81E-09 |
| 140453 | MUC17 | 5.8793601 | -1.2686421 | 1.05E-12 | 9.98E-09 |
| 7409 | VAV1 | 5.08229633 | -0.4953203 | 1.02E-08 | 1.88E-05 |
| 166863 | RBM46 | 4.11034827 | -1.5531224 | 0.00029713 | 0.01668661 |
| 4316 | MMP7 | 4.09507636 | 1.38796344 | 0.00010077 | 0.00829353 |
| 5268 | SERPINB5 | 4.0146409 | -2.8756437 | 1.08E-05 | 0.00197932 |
| 251 | ALPG | 3.96267795 | -2.8352304 | 4.78E-06 | 0.00115223 |
| 100505817 | LINC02582 | 3.92185611 | -3.6910408 | 1.71E-07 | 0.00011569 |
| 10562 | OLFM4 | 3.9179037 | -2.4976577 | 8.77E-07 | 0.0003507 |
| 105369201 | LOC105369201 | 3.91544015 | -2.8246968 | 1.40E-06 | 0.00050468 |
| 2742 | GLRA2 | 3.72336101 | -3.2499759 | 0.0004756 | 0.02221859 |
| 3105 | HLA-A | 3.72285602 | 2.83390012 | 0.00037158 | 0.01907964 |
| 105372412 | LOC105372412 | 3.71461588 | -4.7018611 | 1.17E-06 | 0.00044289 |
| 55510 | DDX43 | 3.67679817 | -0.3194564 | 6.38E-06 | 0.00141611 |
| 90226 | UCN2 | 3.58517721 | -3.5972933 | 0.00026551 | 0.01541766 |
| 64600 | PLA2G2F | 3.55885583 | -4.7154776 | 3.18E-08 | 4.30E-05 |
| 9982 | FGFBP1 | 3.51329936 | -0.8599252 | 8.40E-08 | 9.16E-05 |
| 1048 | CEACAM5 | 3.49539278 | 0.36936832 | 4.16E-09 | 9.84E-06 |
| 242 | ALOX12B | 3.34960309 | -2.2180073 | 6.79E-06 | 0.00147101 |
| 6261 | RYR1 | 3.32212186 | 0.5427569 | 1.81E-08 | 2.77E-05 |
| 4066 | LYL1 | 3.28043589 | -2.3183199 | 4.61E-05 | 0.00496192 |
| 100861550 | PLUT | 3.17026304 | -4.1132126 | 3.16E-07 | 0.00016922 |
| 94 | ACVRL1 | 3.13918217 | 0.37584816 | 2.14E-07 | 0.00013508 |
| 114036 | LINC00310 | 3.13415145 | -3.0557979 | 1.57E-06 | 0.00055077 |
| 250 | ALPP | 3.13144022 | -3.0686352 | 1.12E-05 | 0.00198334 |
| 441161 | OOEP | 3.08963042 | -3.3204331 | 3.57E-05 | 0.00420185 |
| 10331 | B3GNT3 | 3.01255585 | 2.13072569 | 1.47E-06 | 0.00052171 |
| 101927769 | LOC101927769 | 2.99067421 | -4.5307203 | 4.72E-06 | 0.00115223 |
| 119385 | AGAP11 | 2.97380229 | -1.8600933 | 1.14E-06 | 0.00043909 |
| 56978 | PRDM8 | 2.96846565 | -3.6645701 | 3.36E-06 | 0.0009476 |
| 84969 | TOX2 | 2.95699443 | -1.6924997 | 2.51E-05 | 0.00340691 |
| 9048 | ARTN | 2.95128924 | -1.7025722 | 4.65E-05 | 0.00496982 |
| 25825 | BACE2 | 2.88051966 | 2.79318744 | 1.06E-08 | 1.88E-05 |
| 56000 | NXF3 | 2.8577932 | -3.6877621 | 1.09E-05 | 0.00197932 |
| 1510 | CTSE | 2.85341951 | 2.40325879 | 2.66E-06 | 0.00081342 |
| 54857 | GDPD2 | 2.84386852 | -0.4454127 | 0.00014953 | 0.01077627 |
| 3691 | ITGB4 | 2.75500145 | 4.4362814 | 3.40E-12 | 2.41E-08 |
| 10103 | TSPAN1 | 2.74687472 | 3.16585001 | 6.46E-09 | 1.41E-05 |
| 56154 | TEX15 | 2.74162023 | 0.02484527 | 1.41E-05 | 0.00232036 |
| 100130418 | CECR7 | 2.72322247 | -4.0513887 | 6.04E-06 | 0.00136168 |
| 9098 | USP6 | 2.71880018 | -0.8477675 | 2.09E-05 | 0.00302884 |
| 3009 | HIST1H1B | 2.69781522 | -4.365511 | 3.81E-05 | 0.0044123 |
| 56667 | MUC13 | 2.68565787 | 3.87812152 | 8.37E-06 | 0.00169483 |
| 440955 | TMEM89 | 2.62110662 | -4.2811556 | 0.00015336 | 0.01091375 |
| 2859 | GPR35 | 2.61947975 | -0.4996172 | 4.75E-06 | 0.00115223 |
| 101926962 | LOC101926962 | 2.6100496 | -5.094632 | 1.85E-08 | 2.77E-05 |
| 151790 | WDR49 | 2.58314134 | -4.1280767 | 2.60E-05 | 0.0034588 |
| 441307 | HRAT92 | 2.56812623 | -0.9295371 | 2.50E-07 | 0.00014222 |
| 489 | ATP2A3 | 2.52150066 | 1.89758458 | 1.69E-07 | 0.00011569 |
| 3434 | IFIT1 | 2.48307928 | 2.76698698 | 1.63E-07 | 0.00011569 |
| 219855 | SLC37A2 | 2.42836531 | 0.76231482 | 0.00031524 | 0.01734761 |
| 388364 | TMIGD1 | 2.40293874 | -1.9059132 | 0.00015697 | 0.01100503 |
| 7103 | TSPAN8 | 2.38747555 | 3.58414723 | 4.09E-05 | 0.00459074 |
| 5794 | PTPRH | 2.34849833 | 3.46939887 | 5.53E-05 | 0.00551594 |
| 4922 | NTS | 2.31276562 | 2.00681512 | 4.79E-08 | 5.44E-05 |
| 25878 | MXRA5 | 2.31199735 | 3.78250106 | 1.30E-06 | 0.00048626 |
| 154860 | FEZF1-AS1 | 2.30739659 | -3.7322216 | 0.00097657 | 0.03466215 |
| 79983 | POF1B | 2.27488242 | 1.43640266 | 2.49E-09 | 6.43E-06 |
| 51744 | CD244 | 2.26059832 | -1.1026921 | 0.00018205 | 0.01206426 |
| 100873982 | ABCC5-AS1 | 2.24046927 | -5.0454714 | 2.34E-05 | 0.00324221 |
| 84996 | URB1-AS1 | 2.23189879 | -0.0305008 | 6.36E-06 | 0.00141611 |
| 6398 | SECTM1 | 2.20803972 | 0.77895041 | 8.75E-06 | 0.0017251 |
| 23120 | ATP10B | 2.18487719 | -0.2757734 | 0.00049865 | 0.02265482 |
| 1381 | CRABP1 | 2.18402661 | -1.0321357 | 1.55E-05 | 0.00246659 |
| 7476 | WNT7A | 2.17676029 | -0.6046694 | 3.08E-06 | 0.00089269 |
| 374897 | SBSN | 2.15440884 | -4.2388704 | 0.00153382 | 0.04502988 |
| 58985 | IL22RA1 | 2.14702067 | 1.62044576 | 3.66E-07 | 0.00018889 |
| 1893 | ECM1 | 2.13218583 | 4.34816834 | 4.53E-06 | 0.00112739 |
| 4680 | CEACAM6 | 2.12532315 | 4.32214185 | 2.95E-07 | 0.00016125 |
| 126353 | MISP | 2.08466926 | 2.12199898 | 1.55E-07 | 0.00011569 |
| 8313 | AXIN2 | 2.08193009 | 2.4175532 | 1.35E-08 | 2.26E-05 |
| 136 | ADORA2B | 2.07284869 | -0.1916093 | 7.69E-06 | 0.00157187 |
| 8938 | BAIAP3 | 2.07214035 | 1.49188017 | 2.89E-06 | 0.00085353 |
| 248 | ALPI | 2.04911158 | -1.178216 | 0.00016439 | 0.0113061 |
| 9754 | STARD8 | 2.03809621 | 2.55733769 | 4.21E-07 | 0.00019362 |
| 388595 | TMEM82 | 2.02744706 | 1.4528315 | 0.00046656 | 0.022043 |
| 93010 | B3GNT7 | 2.02420896 | 2.89012179 | 1.71E-05 | 0.00262302 |
| 146547 | PRSS36 | 1.98460265 | -0.5297733 | 1.61E-07 | 0.00011569 |
| 2523 | FUT1 | 1.92801793 | 0.42826323 | 9.22E-05 | 0.00786445 |
| 2921 | CXCL3 | 1.8848438 | 0.75262743 | 9.28E-05 | 0.0078869 |
| 26577 | PCOLCE2 | 1.87892951 | 2.8638278 | 4.09E-06 | 0.00104701 |
| 143689 | PIWIL4 | 1.86675743 | 0.36050482 | 8.15E-11 | 4.22E-07 |
| 56106 | PCDHGA10 | 1.84277911 | 1.89933092 | 2.20E-06 | 0.00073529 |
| 84290 | CAPNS2 | 1.84032566 | -4.7479214 | 0.00038536 | 0.01953965 |
| 100820829 | MYZAP | 1.83398732 | -0.4414344 | 1.33E-05 | 0.00224231 |
| 57822 | GRHL3 | 1.76531796 | 0.72356941 | 2.51E-06 | 0.000792 |
| 4359 | MPZ | 1.73030133 | 0.99178967 | 7.97E-05 | 0.00711942 |
| 6035 | RNASE1 | 1.72751019 | 2.64235362 | 4.15E-07 | 0.00019362 |
| 8792 | TNFRSF11A | 1.72510267 | 2.16048489 | 3.63E-07 | 0.00018889 |
| 57188 | ADAMTSL3 | 1.71936498 | 3.19169563 | 2.52E-05 | 0.00340837 |
| 113763 | ZBED6CL | 1.71892827 | 3.22119708 | 6.53E-05 | 0.00613182 |
| 3651 | PDX1 | 1.71735869 | -0.5529028 | 7.11E-05 | 0.00648937 |
| 92610 | TIFA | 1.68836396 | 3.56710345 | 1.07E-07 | 9.49E-05 |
| 2564 | GABRE | 1.68371975 | 1.98649673 | 7.33E-06 | 0.00153125 |
| 51478 | HSD17B7 | 1.68005292 | 3.66015706 | 0.00037101 | 0.01907964 |
| 29881 | NPC1L1 | 1.65486066 | 4.15770772 | 0.00133547 | 0.04153424 |
| 6564 | SLC15A1 | 1.64751662 | 4.80486178 | 1.07E-05 | 0.00197932 |
| 54880 | BCOR | 1.62897965 | 4.72460746 | 4.24E-06 | 0.00106426 |
| 11187 | PKP3 | 1.61805255 | 2.05272847 | 3.09E-10 | 1.24E-06 |
| 360 | AQP3 | 1.61045817 | 4.50554035 | 6.77E-05 | 0.00631991 |
| 26240 | FAM50B | 1.58968528 | 2.13746029 | 6.84E-06 | 0.0014711 |
| 928 | CD9 | 1.58892187 | 6.70269196 | 4.86E-05 | 0.00510628 |
| 101 | ADAM8 | 1.58070327 | 2.02120178 | 0.00010658 | 0.00850885 |
| 53405 | CLIC5 | 1.55995364 | 3.00346297 | 3.90E-07 | 0.00019362 |
| 85315 | PAQR8 | 1.55183729 | 3.85213591 | 1.73E-05 | 0.00262302 |
| 5328 | PLAU | 1.54353526 | 3.20671672 | 1.07E-05 | 0.00197932 |
| 100271927 | RASA4B | 1.53898859 | -1.4432506 | 0.00101826 | 0.03526614 |
| 2224 | FDPS | 1.53015126 | 7.29306822 | 3.50E-10 | 1.24E-06 |
| 79789 | CLMN | 1.51424107 | 4.09610154 | 8.88E-08 | 9.16E-05 |
| 634 | CEACAM1 | 1.50998211 | 5.88237749 | 8.08E-05 | 0.00717701 |
| 4811 | NID1 | 1.48258424 | 6.85992605 | 0.00028087 | 0.01611195 |
| 10718 | NRG3 | 1.4507204 | 0.92036244 | 3.99E-06 | 0.0010291 |
| 120224 | TMEM45B | 1.43755208 | 3.67284175 | 6.99E-07 | 0.00029189 |
| 1595 | CYP51A1 | 1.42884372 | 7.83637169 | 0.00100946 | 0.03526614 |
| 2170 | FABP3 | 1.40417732 | 1.77587438 | 0.00035377 | 0.01849987 |
| 10252 | SPRY1 | 1.37349305 | 3.22975164 | 2.28E-07 | 0.00013975 |
| 79605 | PGBD5 | 1.3617225 | 3.20103816 | 2.60E-06 | 0.00080696 |
| 130271 | PLEKHH2 | 1.35352218 | 2.35192469 | 1.71E-05 | 0.00262302 |
| 4599 | MX1 | 1.34601708 | 4.56648053 | 5.57E-07 | 0.00024335 |
| 1382 | CRABP2 | 1.33783176 | 2.93768141 | 1.82E-05 | 0.00271876 |
| 30845 | EHD3 | 1.32323631 | 3.08723491 | 7.32E-06 | 0.00153125 |
| 2086 | ERV3-1 | 1.31824819 | 3.14499436 | 0.00021815 | 0.01340779 |
| 50853 | VILL | 1.28085942 | 2.6596658 | 0.00022059 | 0.01349941 |
| 5595 | MAPK3 | 1.23823338 | 4.97852722 | 2.61E-06 | 0.00080696 |
| 57458 | TMCC3 | 1.2226149 | 3.17853564 | 1.91E-07 | 0.00012323 |
| 5097 | PCDH1 | 1.21060849 | 6.71566247 | 0.00120621 | 0.03895 |
| 129642 | MBOAT2 | 1.16185006 | 2.52783164 | 2.47E-07 | 0.00014222 |
| 3430 | IFI35 | 1.1376923 | 2.77049213 | 8.77E-06 | 0.0017251 |
| 23474 | ETHE1 | 1.12690427 | 3.73944544 | 3.41E-06 | 0.00094857 |
| 28951 | TRIB2 | 1.09655194 | 3.87979601 | 0.00022214 | 0.01350266 |
| 3959 | LGALS3BP | 1.09580898 | 5.98795977 | 4.27E-07 | 0.00019362 |
| 3428 | IFI16 | 1.0561382 | 4.28369534 | 1.30E-07 | 0.00011078 |
| 5025 | P2RX4 | 1.02896303 | 5.00493648 | 1.09E-05 | 0.00197932 |
| 5125 | PCSK5 | 1.02662445 | 5.04730608 | 4.30E-05 | 0.00477044 |
| 155465 | AGR3 | 1.02377897 | 0.08876047 | 0.00091708 | 0.03323031 |
| 94005 | PIGS | 1.02084833 | 4.98247524 | 3.93E-08 | 4.85E-05 |
| 29937 | NENF | 1.02023361 | 4.23921282 | 7.21E-09 | 1.46E-05 |
| 1291 | COL6A1 | 0.97853895 | 6.82197903 | 0.00015651 | 0.01100503 |
| 200634 | KRTCAP3 | 0.96177498 | 3.16969602 | 1.12E-06 | 0.00043597 |
| 961 | CD47 | 0.96167103 | 5.07404263 | 4.30E-07 | 0.00019362 |
| 23266 | ADGRL2 | 0.95987062 | 5.6540529 | 2.10E-06 | 0.00070974 |
| 7172 | TPMT | 0.92875158 | 5.67073344 | 0.00132422 | 0.04145663 |
| 8612 | PLPP2 | 0.92872184 | 2.75123647 | 3.56E-05 | 0.00420185 |
| 64114 | TMBIM1 | 0.92660202 | 7.53367178 | 2.25E-06 | 0.00073529 |
| 112770 | GLMP | 0.89919014 | 5.71012954 | 7.65E-07 | 0.00031497 |
| 1181 | CLCN2 | 0.88595647 | 2.50150761 | 1.46E-05 | 0.00236089 |
| 29116 | MYLIP | 0.87086257 | 4.06412186 | 0.00022135 | 0.01350266 |
| 3177 | SLC29A2 | 0.83509697 | 4.20278141 | 0.00014821 | 0.01076033 |
| 91860 | CALML4 | 0.78065746 | 4.10401142 | 5.28E-05 | 0.00541183 |
| 79573 | TTC13 | 0.77748784 | 3.90426836 | 1.71E-06 | 0.00059383 |
| 5274 | SERPINI1 | 0.75341831 | 2.84868018 | 0.00013077 | 0.0098497 |
| 23580 | CDC42EP4 | 0.71946441 | 4.87758268 | 4.30E-05 | 0.00477044 |
| 5641 | LGMN | 0.68371855 | 6.70467582 | 0.00151152 | 0.04488417 |
| 57104 | PNPLA2 | 0.61986578 | 5.51176731 | 2.62E-05 | 0.00346331 |
| 1366 | CLDN7 | 0.59615516 | 6.39485696 | 6.15E-07 | 0.00026048 |
| 90780 | PYGO2 | 0.46367665 | 4.6870536 | 7.33E-06 | 0.00153125 |
| 51026 | GOLT1B | -0.4424389 | 5.80575788 | 5.95E-05 | 0.00572556 |
| 51520 | LARS | -0.4868557 | 6.51269771 | 2.50E-05 | 0.0034068 |
| 7342 | UBP1 | -0.5234459 | 6.2490166 | 0.00025526 | 0.01500625 |
| 113 | ADCY7 | -0.5724411 | 3.87105304 | 2.25E-05 | 0.00315872 |
| 55527 | FEM1A | -0.5755952 | 4.83334557 | 2.55E-05 | 0.00342123 |
| 9871 | SEC24D | -0.610231 | 6.50286673 | 0.00035669 | 0.0185159 |
| 2971 | GTF3A | -0.6124858 | 5.89914084 | 5.44E-06 | 0.00124675 |
| 25852 | ARMC8 | -0.7178767 | 5.24372343 | 1.61E-07 | 0.00011569 |
| 10973 | ASCC3 | -0.7627342 | 6.79956474 | 4.02E-05 | 0.00454364 |
| 6118 | RPA2 | -0.7801603 | 4.63381242 | 3.89E-06 | 0.00102364 |
| 2908 | NR3C1 | -0.7920883 | 5.6282937 | 9.68E-05 | 0.00808665 |
| 55226 | NAT10 | -0.7972266 | 5.14823405 | 0.00010316 | 0.00839337 |
| 81609 | SNX27 | -0.8224297 | 5.6541271 | 3.86E-06 | 0.00102364 |
| 91749 | MFSD4B | -0.8814961 | 3.01004168 | 4.99E-07 | 0.0002212 |
| 2002 | ELK1 | -0.9057794 | 5.27907343 | 4.20E-08 | 4.97E-05 |
| 115572 | TENT5B | -0.9099456 | 2.72083971 | 5.94E-06 | 0.00134882 |
| 64762 | GAREM1 | -0.9193477 | 4.72186905 | 3.01E-05 | 0.00379873 |
| 84898 | PLXDC2 | -0.9808355 | 3.72722038 | 3.07E-05 | 0.00384614 |
| 26751 | SH3YL1 | -1.0134293 | 4.25512133 | 2.59E-07 | 0.00014408 |
| 390916 | NUDT19 | -1.1643237 | 2.89277358 | 3.82E-06 | 0.00102354 |
| 3708 | ITPR1 | -1.1928449 | 2.85780149 | 0.00015058 | 0.01079144 |
| 26999 | CYFIP2 | -1.2135037 | 5.06906057 | 4.17E-06 | 0.001058 |
| 5255 | PHKA1 | -1.2781311 | 3.48960161 | 1.29E-05 | 0.00220256 |
| 55840 | EAF2 | -1.3221386 | 1.21277938 | 1.89E-07 | 0.00012323 |
| 5190 | PEX6 | -1.4039241 | 4.91117985 | 7.50E-06 | 0.00154337 |
| 104548973 | SALRNA2 | -1.4058281 | -2.9477197 | 8.34E-05 | 0.00731869 |
| 23240 | TMEM131L | -1.4098181 | 4.89707541 | 2.62E-08 | 3.73E-05 |
| 59345 | GNB4 | -1.4931637 | 4.38601883 | 1.04E-07 | 9.49E-05 |
| 2201 | FBN2 | -1.5231305 | 5.50675784 | 1.71E-05 | 0.00262302 |
| 728588 | MS4A18 | -1.5280497 | -4.9612113 | 0.00061622 | 0.02631205 |
| 205860 | TRIML2 | -1.6140821 | 1.67077812 | 2.36E-07 | 0.00013975 |
| 223117 | SEMA3D | -1.6639622 | 2.40325486 | 8.81E-06 | 0.0017251 |
| 57325 | KAT14 | -1.7275419 | 4.65319695 | 1.63E-07 | 0.00011569 |
| 100996702 | LINC01356 | -1.8418083 | 1.84760909 | 0.00064188 | 0.02685009 |
| 9423 | NTN1 | -1.8630424 | 2.29958172 | 3.70E-06 | 0.00100866 |
| 53904 | MYO3A | -1.8782749 | -0.7623865 | 1.37E-06 | 0.00050347 |
| 1285 | COL4A3 | -1.8927929 | 1.26553081 | 1.40E-06 | 0.00050468 |
| 100133091 | LOC100133091 | -1.9001609 | -5.1249711 | 1.02E-05 | 0.00191805 |
| 1286 | COL4A4 | -1.9511209 | 3.17313463 | 3.71E-08 | 4.79E-05 |
| 386617 | KCTD8 | -2.0026816 | -0.7086124 | 9.60E-06 | 0.00184123 |
| 651746 | ANKRD33B | -2.0582806 | 3.87280285 | 2.23E-06 | 0.00073529 |
| 10720 | UGT2B11 | -2.0666898 | 2.77417785 | 0.00036572 | 0.01884677 |
| 115207 | KCTD12 | -2.162618 | 4.23068237 | 1.12E-06 | 0.00043597 |
| 728215 | FAM155A | -2.1785848 | 0.20179494 | 0.0008994 | 0.0327835 |
| 139065 | SLITRK4 | -2.191255 | 2.31484319 | 8.91E-11 | 4.22E-07 |
| 83478 | ARHGAP24 | -2.3040421 | 1.6833765 | 2.36E-07 | 0.00013975 |
| 23359 | FAM189A1 | -2.5058558 | -2.7431907 | 5.27E-06 | 0.00121675 |
| 100288428 | LMCD1-AS1 | -2.532167 | -1.5217778 | 8.06E-07 | 0.00032692 |
| 139793 | PAGE3 | -2.7553626 | -3.9559308 | 3.52E-05 | 0.00419735 |
| 6565 | SLC15A2 | -2.7573402 | 3.16311415 | 1.98E-13 | 2.81E-09 |
| 83851 | SYT16 | -2.8249054 | -0.6960514 | 1.72E-05 | 0.00262302 |
| 827 | CAPN6 | -2.8668929 | 1.10172076 | 2.18E-05 | 0.00309128 |
| 100506013 | APELA | -2.8826317 | 0.23197274 | 4.08E-07 | 0.00019362 |
| 100507524 | ARHGEF26-AS1 | -2.9485961 | -2.4887774 | 3.04E-05 | 0.00382105 |
| 100861510 | LRRC3-DT | -3.2281414 | -2.788914 | 9.30E-05 | 0.0078869 |
| 152816 | ODAPH | -3.3526511 | -0.2658878 | 1.68E-09 | 5.30E-06 |
| 165257 | C1QL2 | -3.4273245 | -1.380828 | 7.45E-06 | 0.00154337 |
| 4846 | NOS3 | -3.4912055 | 1.41536675 | 0.00036469 | 0.01883525 |
| 353345 | GPR141 | -5.2770257 | -2.1147412 | 2.27E-09 | 6.43E-06 |

**Supplementary Table 2.** Datasets used to generate the Venn diagram comparing differentially expressed genes (DEGs) in HLC- HT1 vs HLC-CTRL and DEGs previously identified in transcriptomic analysis carried out in HT1 and in several liver conditions.

|  | References | Genes |
| --- | --- | --- |
| HYPERTYROSINEMIA TYPE 1 (in mice, experimental) | | TOTAL: 1198 |
| Luijerink, M.C. et al. Journal of Hepatology (2003) | | 86 |
| Angileri, F et al., Cancers (2014) | | 552 |
| Pletscher-Frankild, S et al., Methods (2015) | | 91 |
| Colemonts-Vroninks, H et al., Genes (2021) | | 523 |
| HEPATOCELLULAR CARCINOMA | | TOTAL: 5840 |
| GSE Liver Carcinoma | C2239176 | 5725 |
| GSE Liver neoplasm | C0023903 | 1424 |
| LIVER FIBROSIS AND CIRRHOSIS | | TOTAL: 2409 |
| GSE Cirrhosis of the liver | C0023890 | 1182 |
| GSE Liver Cirrhosis, Experimental | C0023893 | 870 |
| GSE Liver fibrosis | C0239946 | 1179 |
| CELLULAR RESPONSE TO STRESS | | TOTAL: 2742 |
| GENE ONTOLOGY Cellular Response to stress | GO:0033554 | 1831 |
| GENE ONTOLOGY Inflammatory Response to stress | GO:0006954 | 735 |
| GENE ONTOLOGY Response to Oxidative Stress | GO:0006979 | 444 |
| GENE ONTOLOGY Response to ER Stress | GO:0034976 | 295 |
| REACTOME Oxidative Stress induced Senescence | R-HSA-2559580 | 126 |
| WIKIPATHWAYS Oxidative Stress | WP408 | 34 |
| WEIGEL Oxidative Stress Response | M14591 | 33 |
| BIOCARTA Stress Pathway | M9670 | 24 |
| APOPTOSIS | | TOTAL: 2194 |
| GENE ONTOLOGY Apoptotic Process | GO:0006915 | 1932 |
| REACTOME Apoptosis | R-HSA-109581 | 179 |
| HALLMARK Apoptosis | M5902 | 161 |
| WIKIPATHWAYS Apoptosis Modulation & Signaling | WP1772 | 93 |
| ALCALA Apoptosis | M16169 | 87 |
| KEGG Apoptosis | HSA04210 | 87 |
| WIKIPATHWAYS Apoptosis | WP254 | 86 |
| OTHER LIVER PHYSIOPATHOLOGIES | | TOTAL: 1836 |
| GSE Liver Diseases | C0023895 | 1019 |
| GSE Hepatic Steatosis | C2711227 | 1143 |
| GSE Hepatitis | C0019158 | 656 |
| GSE Hepatomegaly | C0019209 | 523 |
| GSE Hepatic failure | C0085605 | 293 |
| GSE Abnormality of the liver | C4021780 | 75 |
| GSE Liver dysfunction | C0086565 | 73 |
| GSE Decreased liver function | C0232744 | 59 |

**Supplementary Table 3.** List of genes upregulated in HLC-HT1 despite treatment with 50 µM NTBC (Fig. 6F).

| **Upregulated genes despite NTBC treatment (FDR ≤ 0.05)** | | | | | |
| --- | --- | --- | --- | --- | --- |
| **ENTREZID** | **SYMBOL** | **logFC** | **p-value** | **FDR** | **(-) log10 for volcano plot** |
| 728927 | ZNF736 | 6.629619784 | 0.00000196 | 0.002654953 | 5.707743929 |
| 140453 | MUC17 | 6.600029006 | 0.00000155 | 0.002595449 | 5.809668302 |
| 5646 | PRSS3 | 3.806450671 | 0.0000225 | 0.010543405 | 4.647817482 |
| 144486 | CEP83-DT | 3.387100743 | 0.000239951 | 0.044439898 | 3.619877436 |
| 56978 | PRDM8 | 3.021333255 | 1.86E-08 | 0.000176182 | 7.730487056 |
| 8638 | OASL | 2.962649853 | 0.000227182 | 0.043590094 | 3.643626081 |
| 219855 | SLC37A2 | 2.797583056 | 0.00000141 | 0.002595449 | 5.850780887 |
| 2859 | GPR35 | 2.776656114 | 0.0000313 | 0.012883871 | 4.504455662 |
| 79861 | TUBAL3 | 2.705837782 | 0.0000942 | 0.029568464 | 4.025949097 |
| 114036 | LINC00310 | 2.690554085 | 0.000216948 | 0.04307869 | 3.663644349 |
| 4066 | LYL1 | 2.506811206 | 0.000147643 | 0.034936113 | 3.830787139 |
| 2710 | GK | 2.430174754 | 0.000117275 | 0.0322211 | 3.930794558 |
| 10331 | B3GNT3 | 2.202653084 | 0.00000169 | 0.002595449 | 5.772113295 |
| 94 | ACVRL1 | 2.123202482 | 0.000000862 | 0.001997085 | 6.064492734 |
| 8786 | RGS11 | 2.026950889 | 0.0000648 | 0.023001014 | 4.188424994 |
| **3691** | **ITGB4** | **2.017122961** | **0.00000364** | **0.004138921** | **5.438898616** |
| **7103** | **TSPAN8** | **2.00191847** | **0.0000533** | **0.019165124** | **4.273272791** |
| 3434 | IFIT1 | 1.974345802 | 0.0000283 | 0.012089533 | 4.548213564 |
| 8651 | SOCS1 | 1.960263543 | 0.0000248 | 0.011184756 | 4.605548319 |
| 26585 | GREM1 | 1.931876397 | 0.000109999 | 0.031234182 | 3.958611263 |
| 84815 | MGC12916 | 1.929777638 | 1.3E-09 | 0.000037 | 8.886056648 |
| 55532 | SLC30A10 | 1.891235278 | 0.000000128 | 0.000536058 | 6.89279003 |
| 81615 | TMEM163 | 1.89079597 | 0.0000227 | 0.010543405 | 4.643974143 |
| 1029 | CDKN2A | 1.841782892 | 0.000260508 | 0.046522762 | 3.584178935 |
| 5243 | ABCB1 | 1.804454998 | 0.0000137 | 0.007717755 | 4.863279433 |
| 79983 | POF1B | 1.803571917 | 0.000242584 | 0.044439898 | 3.615137847 |
| 51090 | PLLP | 1.670600488 | 0.000275702 | 0.04773513 | 3.559560083 |
| 23120 | ATP10B | 1.66162165 | 0.0000529 | 0.019165124 | 4.276544328 |
| 6286 | S100P | 1.661532362 | 0.000230775 | 0.043590094 | 3.63681124 |
| 1294 | COL7A1 | 1.569986263 | 0.0000212 | 0.010192394 | 4.673664139 |
| 27284 | SULT1B1 | 1.529978086 | 0.000138695 | 0.034245716 | 3.857939195 |
| 7476 | WNT7A | 1.395127041 | 0.0000105 | 0.00661906 | 4.978810701 |
| 5787 | PTPRB | 1.38257491 | 0.0000206 | 0.010192394 | 4.68613278 |
| 2646 | GCKR | 1.356619994 | 0.000000114 | 0.000536058 | 6.943095149 |
| 4359 | MPZ | 1.325693027 | 0.000103314 | 0.030439666 | 3.985840824 |
| 342979 | PALM3 | 1.325123198 | 0.000167965 | 0.037015478 | 3.774781206 |
| 7098 | TLR3 | 1.309970176 | 0.000272697 | 0.047504473 | 3.56431964 |
| 4940 | OAS3 | 1.301532425 | 0.000264276 | 0.046900649 | 3.577942275 |
| 143689 | PIWIL4 | 1.28599154 | 0.000215288 | 0.043049947 | 3.666980177 |
| 53345 | TM6SF2 | 1.285931619 | 0.00000943 | 0.006461467 | 5.025488307 |
| 26027 | ACOT11 | 1.285595189 | 0.000103985 | 0.030439666 | 3.983029304 |
| 220323 | OAF | 1.250442 | 9.65E-09 | 0.000137063 | 8.015472687 |
| 8824 | CES2 | 1.237286109 | 0.0000188 | 0.009717961 | 4.725842151 |
| **7066** | **THPO** | **1.236468995** | **0.000000132** | **0.000536058** | **6.879426069** |
| **9245** | **GCNT3** | **1.23463443** | **0.0000107** | **0.00661906** | **4.970616222** |
| 8313 | AXIN2 | 1.234014839 | 0.0000285 | 0.012089533 | 4.54515514 |
| **10110** | **SGK2** | **1.13320688** | **0.0000323** | **0.013090577** | **4.490797478** |
| **2810** | **SFN** | **1.124374905** | **0.0000257** | **0.011418911** | **4.590066877** |
| 2224 | FDPS | 1.07719361 | 0.000085 | 0.027638254 | 4.070581074 |
| **6091** | **ROBO1** | **1.076913675** | **0.0000787** | **0.026306097** | **4.104025268** |
| 7104 | TM4SF4 | 1.070739455 | 0.0000332 | 0.01329092 | 4.478861916 |
| 282969 | FUOM | 1.065842025 | 0.0000862 | 0.027638254 | 4.064492734 |
| **28951** | **TRIB2** | **1.015804272** | **0.0000377** | **0.01425585** | **4.42365865** |
| 79639 | TMEM53 | 0.952959439 | 0.0000692 | 0.024193332 | 4.159893906 |
| 183 | AGT | 0.938959593 | 0.0000711 | 0.024314429 | 4.148130399 |
| 9123 | SLC16A3 | 0.895762669 | 0.000102352 | 0.030439666 | 3.989903667 |
| 648987 | LOC648987 | 0.890882279 | 0.000197607 | 0.040659757 | 3.704197675 |
| **3959** | **LGALS3BP** | **0.890583655** | **0.000137211** | **0.034176495** | **3.86261107** |
| 113791 | PIK3IP1 | 0.839394257 | 0.0000204 | 0.010192394 | 4.690369833 |
| 27163 | NAAA | 0.82543215 | 0.00000613 | 0.005669956 | 5.212539525 |
| 79789 | CLMN | 0.811332999 | 0.000135578 | 0.034068467 | 3.867810777 |
| 6640 | SNTA1 | 0.80479083 | 0.000016 | 0.008713654 | 4.795880017 |
| 101928099 | PCAT7 | 0.764203455 | 0.00024619 | 0.04452581 | 3.608729592 |
| 25981 | DNAH1 | 0.733872878 | 0.000129252 | 0.03379236 | 3.888562728 |
| 79762 | C1orf115 | 0.719499364 | 0.000151462 | 0.035305344 | 3.819696313 |
| 1384 | CRAT | 0.712990684 | 0.0000474 | 0.017470762 | 4.324221658 |
| 140885 | SIRPA | 0.708950995 | 0.000000124 | 0.000536058 | 6.906578315 |
| 2626 | GATA4 | 0.672199832 | 0.00000619 | 0.005669956 | 5.208309351 |
| 7464 | CORO2A | 0.649098435 | 0.0000187 | 0.009717961 | 4.728158393 |
| 5833 | PCYT2 | 0.635575383 | 0.00011663 | 0.0322211 | 3.933189724 |
| 55243 | KIRREL1 | 0.628821688 | 0.000230883 | 0.043590094 | 3.636608043 |
| **374** | **AREG** | **0.62838846** | **0.00000841** | **0.006280776** | **5.075204004** |
| 64856 | VWA1 | 0.625381089 | 0.00000908 | 0.006461467 | 5.041914151 |
| 56997 | COQ8A | 0.622185788 | 0.00000795 | 0.006280776 | 5.099632871 |
| 105370828 | NA (ENTREZID: 105370828) | 0.618874608 | 0.00000053 | 0.001368282 | 6.27572413 |
| 57104 | PNPLA2 | 0.579203831 | 0.00000174 | 0.002595449 | 5.759450752 |
| 326625 | MMAB | 0.576927868 | 0.000152934 | 0.035305344 | 3.815495952 |
| 4835 | NQO2 | 0.573612848 | 0.000205802 | 0.041445099 | 3.686550409 |
| **55353** | **LAPTM4B** | **0.557242308** | **0.0000118** | **0.006955334** | **4.928117993** |
| 55001 | TTC22 | 0.548299422 | 0.0000139 | 0.007717755 | 4.8569852 |
| 11346 | SYNPO | 0.537475108 | 0.000163136 | 0.037015478 | 3.787450191 |
| 3672 | ITGA1 | 0.473768492 | 0.000219682 | 0.043318587 | 3.658205526 |
| **4734** | **NEDD4** | **0.447467428** | **0.00000742** | **0.006280776** | **5.129596095** |
| 65124 | SOWAHC | 0.405399285 | 0.00000219 | 0.002822808 | 5.659555885 |
| **3068** | **HDGF** | **0.244425388** | **0.000183864** | **0.038961422** | **3.735503296** |
| **Downregulated genes despite NTBC treatment (FDR ≤ 0.05)** | | | | | |
| **ENTREZID** | **SYMBOL** | **logFC** | **p-value** | **FDR** | **(-) log10 for volcano plot** |
| 10899 | JTB | -0.219401956 | 0.000278935 | 0.047961106 | 3.554496988 |
| 4170 | MCL1 | -0.232642794 | 0.00014104 | 0.034524521 | 3.850657701 |
| 51569 | UFM1 | -0.266387911 | 0.000133152 | 0.034061776 | 3.875652306 |
| 4799 | NFX1 | -0.280941004 | 0.000190616 | 0.039507628 | 3.719840648 |
| 54801 | HAUS6 | -0.301771085 | 0.000114375 | 0.032155242 | 3.941668893 |
| 116224 | FAM122A | -0.304899848 | 0.000146442 | 0.034936113 | 3.834334348 |
| 8027 | STAM | -0.314232928 | 0.000225446 | 0.043590094 | 3.646957466 |
| 6446 | SGK1 | -0.336969179 | 0.0000233 | 0.01064995 | 4.632644079 |
| 23002 | DAAM1 | -0.352468864 | 0.0000102 | 0.00661906 | 4.991399828 |
| 122809 | SOCS4 | -0.35291604 | 0.0000451 | 0.01686536 | 4.345823458 |
| 51026 | GOLT1B | -0.357021915 | 0.00000587 | 0.005669956 | 5.231361899 |
| 148423 | C1orf52 | -0.357822777 | 0.000202458 | 0.041062795 | 3.693665058 |
| 54915 | YTHDF1 | -0.361208782 | 0.0000209 | 0.010192394 | 4.679853714 |
| 729830 | FAM160A1 | -0.361769517 | 0.000290384 | 0.049373949 | 3.537027317 |
| 7286 | TUFT1 | -0.364481846 | 0.00000776 | 0.006280776 | 5.110138279 |
| 55286 | C4orf19 | -0.369985584 | 0.0000111 | 0.006703146 | 4.954677021 |
| 11216 | AKAP10 | -0.38816536 | 0.000238957 | 0.044439898 | 3.621680243 |
| 81617 | CAB39L | -0.400572329 | 0.000168163 | 0.037015478 | 3.774269554 |
| 57182 | ANKRD50 | -0.411448764 | 0.0000106 | 0.00661906 | 4.974694135 |
| 440515 | ZNF506 | -0.423676679 | 0.0000842 | 0.027638254 | 4.074687909 |
| 5074 | PAWR | -0.458914905 | 0.0000135 | 0.007717755 | 4.869666232 |
| 10424 | PGRMC2 | -0.472135689 | 0.00000167 | 0.002595449 | 5.777283529 |
| 23767 | FLRT3 | -0.49661312 | 0.00000383 | 0.004186547 | 5.416801226 |
| 6509 | SLC1A4 | -0.497398695 | 0.000294746 | 0.049817261 | 3.53055208 |
| 254251 | LCORL | -0.498655857 | 0.000228112 | 0.043590094 | 3.641851868 |
| 58480 | RHOU | -0.500530547 | 0.000000181 | 0.000633466 | 6.742321425 |
| 11221 | DUSP10 | -0.517826676 | 0.000185953 | 0.039023826 | 3.730596811 |
| 2908 | NR3C1 | -0.534860258 | 0.0000065 | 0.005766717 | 5.187086643 |
| 84898 | PLXDC2 | -0.566266343 | 0.000245342 | 0.04452581 | 3.610228099 |
| 83604 | TMEM47 | -0.568921074 | 0.000000961 | 0.001997085 | 6.017276612 |
| 956 | ENTPD3 | -0.569230073 | 0.0000958 | 0.029568464 | 4.018634491 |
| 2239 | GPC4 | -0.580567324 | 0.000000429 | 0.001218004 | 6.367542708 |
| 3491 | CCN1 | -0.592122282 | 0.0000178 | 0.009514639 | 4.749579998 |
| 158586 | ZXDB | -0.624003192 | 0.000231805 | 0.043590094 | 3.634877201 |
| 23584 | VSIG2 | -0.636747594 | 0.0000949 | 0.029568464 | 4.022733788 |
| 11167 | FSTL1 | -0.637545962 | 0.000000985 | 0.001997085 | 6.00656377 |
| 148206 | ZNF714 | -0.645657217 | 0.000179237 | 0.038556421 | 3.746572334 |
| 6638 | SNRPN | -0.685510587 | 0.00000825 | 0.006280776 | 5.083546051 |
| 6579 | SLCO1A2 | -0.772104018 | 0.0000285 | 0.012089533 | 4.54515514 |
| 114880 | OSBPL6 | -0.779469854 | 0.000107569 | 0.030916002 | 3.968312869 |
| 10544 | PROCR | -0.789770932 | 0.000170302 | 0.037119346 | 3.768780252 |
| 23284 | ADGRL3 | -0.804657674 | 0.000280385 | 0.047961106 | 3.552245224 |
| 100506428 | CBR3-AS1 | -0.83438971 | 0.000101915 | 0.030439666 | 3.991761891 |
| 283455 | KSR2 | -0.866540161 | 0.00012067 | 0.032324882 | 3.918400687 |
| 100132111 | C2CD4D-AS1 | -0.879735908 | 0.000270921 | 0.047504473 | 3.56715733 |
| 23768 | FLRT2 | -0.917598528 | 0.00012973 | 0.03379236 | 3.886959582 |
| 143686 | SESN3 | -0.961796454 | 0.000000201 | 0.000633466 | 6.696803943 |
| 6565 | SLC15A2 | -1.034995605 | 0.00000956 | 0.006461467 | 5.019542108 |
| 139065 | SLITRK4 | -1.063532869 | 0.00000954 | 0.006461467 | 5.020451625 |
| 101927873 | LINC01508 | -1.100882274 | 0.000186908 | 0.039023826 | 3.72837211 |
| 54873 | PALMD | -1.105983628 | 0.00000486 | 0.005114582 | 5.313363731 |
| 9423 | NTN1 | -1.33155843 | 0.00000583 | 0.005669956 | 5.234331445 |
| 53904 | MYO3A | -1.413704192 | 0.0000346 | 0.01335913 | 4.460923901 |
| 1740 | DLG2 | -1.453955621 | 0.000241078 | 0.044439898 | 3.61784242 |
| 940 | CD28 | -1.524057927 | 0.00000313 | 0.003709014 | 5.504455662 |
| 29947 | DNMT3L | -1.524057927 | 0.00000313 | 0.003709014 | 5.504455662 |
| 54769 | DIRAS2 | -1.750474971 | 0.00000828 | 0.006280776 | 5.081969663 |
| 100133091 | LOC100133091 | -1.893677599 | 0.00017125 | 0.037119346 | 3.76636942 |
| 389898 | UBE2NL | -1.895103077 | 0.0000312 | 0.012883871 | 4.505845406 |
| **Not significant and normalized with NTBC treatment (FDR ≥ 0.05)** | | | | | |
| **ENTREZID** | **SYMBOL** | **logFC** | **p-value** | **FDR** | **(-) log10 for volcano plot** |
| 8939 | FUBP3 | -0.141786815 | 0.004984196 | 0.192552718 | 2.302404888 |
| 9318 | COPS2 | -0.165507173 | 0.008433972 | 0.245120413 | 2.073967845 |
| 4150 | MAZ | -0.172750682 | 0.005073581 | 0.193895468 | 2.294685402 |
| 5932 | RBBP8 | -0.173302376 | 0.003487217 | 0.163668657 | 2.457521027 |
| 1459 | CSNK2A2 | -0.176746458 | 0.007137215 | 0.228479379 | 2.14647122 |
| 5805 | PTS | -0.178477848 | 0.007306624 | 0.229467755 | 2.136283241 |
| 56681 | SAR1A | -0.1785402 | 0.001075176 | 0.090047638 | 2.968520438 |
| 10328 | EMC8 | -0.178558285 | 0.005555084 | 0.203003874 | 2.25530937 |
| 93624 | TADA2B | -0.186919652 | 0.006532784 | 0.216525907 | 2.184901701 |
| 23087 | TRIM35 | -0.189680073 | 0.002953207 | 0.154939588 | 2.529706111 |
| 55813 | UTP6 | -0.191793524 | 0.007896134 | 0.237511356 | 2.10258549 |
| 5048 | PAFAH1B1 | -0.193720711 | 0.003367345 | 0.16124074 | 2.472712386 |
| 57511 | COG6 | -0.194113226 | 0.004999079 | 0.192795108 | 2.30111 |
| 342371 | ATXN1L | -0.195452261 | 0.009639392 | 0.259687407 | 2.015950358 |
| 3981 | LIG4 | -0.199612752 | 0.00374305 | 0.17170258 | 2.426774372 |
| 55623 | THUMPD1 | -0.201580142 | 0.008960296 | 0.252408336 | 2.047677643 |
| 80209 | PROSER1 | -0.204164576 | 0.009657809 | 0.259936967 | 2.015121388 |
| 10413 | YAP1 | -0.206333528 | 0.007794044 | 0.235688905 | 2.108237147 |
| 22930 | RAB3GAP1 | -0.210338141 | 0.001745515 | 0.116074737 | 2.758076415 |
| 92609 | TIMM50 | -0.211789609 | 0.002209144 | 0.131890902 | 2.655775974 |
| 8886 | DDX18 | -0.212004822 | 0.00333306 | 0.160682928 | 2.477156868 |
| 4238 | MFAP3 | -0.217136773 | 0.002903553 | 0.153571991 | 2.537070242 |
| 5229 | PGGT1B | -0.217222685 | 0.006112985 | 0.210482178 | 2.21374667 |
| 148867 | SLC30A7 | -0.218296164 | 0.00973001 | 0.261090495 | 2.011886713 |
| 6461 | SHB | -0.218641517 | 0.000667392 | 0.069238034 | 3.175619004 |
| 1974 | EIF4A2 | -0.218803293 | 0.001915317 | 0.120588544 | 2.717759337 |
| 170506 | DHX36 | -0.21948693 | 0.00915407 | 0.254356697 | 2.038385771 |
| 9868 | TOMM70 | -0.220120835 | 0.001693031 | 0.114461016 | 2.77133509 |
| 65005 | MRPL9 | -0.220787593 | 0.007187171 | 0.228991676 | 2.143442022 |
| 9255 | AIMP1 | -0.221504716 | 0.007211181 | 0.229039691 | 2.141993604 |
| 10818 | FRS2 | -0.221541707 | 0.001328535 | 0.09849545 | 2.876627 |
| 22931 | RAB18 | -0.221988779 | 0.007746422 | 0.234999627 | 2.110898848 |
| 11177 | BAZ1A | -0.222383933 | 0.002214436 | 0.131890902 | 2.654736867 |
| 178 | AGL | -0.22425185 | 0.008837032 | 0.2511787 | 2.053693572 |
| 55218 | EXD2 | -0.226227715 | 0.002245192 | 0.132413438 | 2.648746514 |
| 27297 | CRCP | -0.227400272 | 0.000437088 | 0.059383377 | 3.359431117 |
| 51122 | COMMD2 | -0.229993852 | 0.000562179 | 0.065155448 | 3.250125381 |
| 150275 | CCDC117 | -0.230354168 | 0.000642909 | 0.068888332 | 3.191850495 |
| 55671 | PPP4R3A | -0.231119468 | 0.001230491 | 0.096238624 | 2.909921558 |
| 113174 | SAAL1 | -0.231928829 | 0.004892201 | 0.191341678 | 2.310495708 |
| 4088 | SMAD3 | -0.232145239 | 0.00059932 | 0.066475312 | 3.222341229 |
| 84747 | UNC119B | -0.23281748 | 0.004298542 | 0.183268934 | 2.366678826 |
| 9871 | SEC24D | -0.23348172 | 0.000361317 | 0.054591567 | 3.442111604 |
| 8065 | CUL5 | -0.234603223 | 0.004941736 | 0.191975237 | 2.306120459 |
| 55031 | USP47 | -0.235287083 | 0.004629343 | 0.187812491 | 2.33448064 |
| 90196 | SYS1 | -0.237224553 | 0.001943153 | 0.121195766 | 2.711493003 |
| 84612 | PARD6B | -0.237372938 | 0.00034959 | 0.053890511 | 3.456440999 |
| 8446 | DUSP11 | -0.239361295 | 0.007684371 | 0.234612665 | 2.114391676 |
| 160760 | PPTC7 | -0.239454664 | 0.005904546 | 0.208014385 | 2.22881349 |
| 4659 | PPP1R12A | -0.239757102 | 0.000695651 | 0.069552812 | 3.157608586 |
| 64844 | MARCHF7 | -0.240469348 | 0.002794874 | 0.151162781 | 2.553637766 |
| 5937 | RBMS1 | -0.240779738 | 0.007889906 | 0.237511356 | 2.102928171 |
| 8019 | BRD3 | -0.245115828 | 0.008145906 | 0.240645298 | 2.089060606 |
| 50999 | TMED5 | -0.247514725 | 0.00031032 | 0.051189683 | 3.508190233 |
| 663 | BNIP2 | -0.24837381 | 0.008803699 | 0.250935818 | 2.055334814 |
| 57337 | SENP7 | -0.248634633 | 0.004647812 | 0.187812491 | 2.332751447 |
| 79872 | CBLL1 | -0.251157871 | 0.00564166 | 0.203810348 | 2.248593091 |
| 79738 | BBS10 | -0.253406446 | 0.00500405 | 0.192795108 | 2.30067836 |
| 57606 | SLAIN2 | -0.25508911 | 0.008807805 | 0.250935818 | 2.055132309 |
| 56888 | KCMF1 | -0.255232383 | 0.004074476 | 0.178266186 | 2.389928236 |
| 9044 | BTAF1 | -0.257095813 | 0.004497302 | 0.185633035 | 2.347047948 |
| 3364 | HUS1 | -0.25956686 | 0.001293032 | 0.097647996 | 2.888390727 |
| 4293 | MAP3K9 | -0.259920754 | 0.000311879 | 0.051189683 | 3.506013867 |
| 8504 | PEX3 | -0.261021462 | 0.002281824 | 0.133317689 | 2.641717856 |
| 55686 | MREG | -0.263298467 | 0.002123482 | 0.128289957 | 2.672951416 |
| 7325 | UBE2E2 | -0.263985217 | 0.004682915 | 0.18861186 | 2.329483725 |
| 221092 | HNRNPUL2 | -0.269281839 | 0.003928336 | 0.176310508 | 2.405791373 |
| 153129 | SLC38A9 | -0.269482613 | 0.004447363 | 0.18559875 | 2.351897421 |
| 55331 | ACER3 | -0.27360071 | 0.002846157 | 0.152197023 | 2.545741147 |
| 2002 | ELK1 | -0.274125846 | 0.000562002 | 0.065155448 | 3.250262139 |
| 51571 | CYRIB | -0.274914257 | 0.00136771 | 0.099890634 | 2.864005978 |
| 51701 | NLK | -0.275199464 | 0.004356677 | 0.1835428 | 2.360844637 |
| 5876 | RABGGTB | -0.275358465 | 0.007777979 | 0.235453863 | 2.109133234 |
| 149986 | LSM14B | -0.276135496 | 0.004022592 | 0.177087614 | 2.395494014 |
| 51163 | DBR1 | -0.277443253 | 0.000319875 | 0.051402596 | 3.495019701 |
| 55527 | FEM1A | -0.277495947 | 0.005307188 | 0.198019459 | 2.275135528 |
| 8897 | MTMR3 | -0.278924038 | 0.004216121 | 0.181664271 | 2.375086934 |
| 26985 | AP3M1 | -0.278971978 | 0.000674302 | 0.069238034 | 3.171145552 |
| 23099 | ZBTB43 | -0.279144608 | 0.004551753 | 0.186593911 | 2.341821313 |
| 23111 | SPART | -0.282192558 | 0.000802435 | 0.075622593 | 3.095590137 |
| 79009 | DDX50 | -0.282893863 | 0.00435406 | 0.1835428 | 2.361105591 |
| 51174 | TUBD1 | -0.283542029 | 0.006385856 | 0.214753 | 2.194780879 |
| 23390 | ZDHHC17 | -0.284035545 | 0.004309439 | 0.183458058 | 2.365579262 |
| 80781 | COL18A1 | -0.287966607 | 0.000421609 | 0.058398024 | 3.375090127 |
| 57109 | REXO4 | -0.288651949 | 0.007269541 | 0.22933191 | 2.13849301 |
| 3516 | RBPJ | -0.288967627 | 0.007101483 | 0.228303396 | 2.148650948 |
| 51566 | ARMCX3 | -0.289775004 | 0.000444546 | 0.060108942 | 3.352083293 |
| 54465 | ETAA1 | -0.290308906 | 0.00175627 | 0.116474313 | 2.755408717 |
| 2152 | F3 | -0.291315377 | 0.002909728 | 0.153571991 | 2.536147607 |
| 55142 | HAUS2 | -0.293552762 | 0.001528902 | 0.106928972 | 2.815620351 |
| 9693 | RAPGEF2 | -0.295290955 | 0.000580805 | 0.066232742 | 3.235969654 |
| 7456 | WIPF1 | -0.297769159 | 0.006139456 | 0.210482178 | 2.211870109 |
| 55833 | UBAP2 | -0.298724 | 0.000686228 | 0.069238034 | 3.163531566 |
| 64764 | CREB3L2 | -0.299693982 | 0.000406266 | 0.057079117 | 3.391189522 |
| 9113 | LATS1 | -0.300339714 | 0.000564613 | 0.065171503 | 3.248249127 |
| 22834 | ZNF652 | -0.302788926 | 0.008919623 | 0.251716736 | 2.049653501 |
| 91298 | C12orf29 | -0.304011545 | 0.000398255 | 0.056532267 | 3.399838763 |
| 153443 | SRFBP1 | -0.304924009 | 0.006142872 | 0.210482178 | 2.211628534 |
| 9183 | ZW10 | -0.30503508 | 0.000903314 | 0.08116964 | 3.044161259 |
| 121512 | FGD4 | -0.305656176 | 0.000423715 | 0.058404846 | 3.372926161 |
| 9645 | MICAL2 | -0.308777259 | 0.008369921 | 0.24425889 | 2.077278641 |
| 23468 | CBX5 | -0.309795522 | 0.000544981 | 0.064791918 | 3.263618639 |
| 55276 | PGM2 | -0.312989502 | 0.007633444 | 0.234109795 | 2.117279476 |
| 154043 | CNKSR3 | -0.317075385 | 0.004330288 | 0.1835428 | 2.363483219 |
| 49854 | ZBTB21 | -0.318690933 | 0.006497026 | 0.216275551 | 2.187285395 |
| 55795 | PCID2 | -0.318727599 | 0.002182276 | 0.131005794 | 2.661090324 |
| 55074 | OXR1 | -0.321099513 | 0.000323737 | 0.051402596 | 3.489807662 |
| 133 | ADM | -0.325167964 | 0.001462958 | 0.103887127 | 2.834768142 |
| 133619 | PRRC1 | -0.327920307 | 0.001123547 | 0.091939854 | 2.949408756 |
| 116068 | LYSMD3 | -0.328476042 | 0.000314118 | 0.051260748 | 3.502907176 |
| 54989 | ZNF770 | -0.329418042 | 0.003476318 | 0.163427233 | 2.458880503 |
| 7813 | EVI5 | -0.330642603 | 0.009038113 | 0.253844914 | 2.043922233 |
| 643650 | LINC00842 | -0.335198827 | 0.007201285 | 0.228991676 | 2.142590001 |
| 27443 | CECR2 | -0.335472712 | 0.00591755 | 0.208144184 | 2.227858064 |
| 7644 | ZNF91 | -0.336748187 | 0.000506309 | 0.062971482 | 3.295584353 |
| 161424 | NOP9 | -0.336917413 | 0.002240164 | 0.132413438 | 2.649720186 |
| 3570 | IL6R | -0.338697175 | 0.001112132 | 0.091533314 | 2.953843663 |
| 10477 | UBE2E3 | -0.340943678 | 0.001626997 | 0.111261127 | 2.788613248 |
| 89853 | MVB12B | -0.346463455 | 0.004947366 | 0.191975237 | 2.30562596 |
| 151246 | SGO2 | -0.348897937 | 0.005616334 | 0.203739022 | 2.250547073 |
| 9440 | MED17 | -0.354811693 | 0.001714759 | 0.115380501 | 2.765796909 |
| 9846 | GAB2 | -0.358756893 | 0.001238182 | 0.096238624 | 2.907215514 |
| 55175 | KLHL11 | -0.359410778 | 0.000466975 | 0.061633976 | 3.330706369 |
| 65998 | C11orf95 | -0.363970347 | 0.007861282 | 0.237216892 | 2.104506624 |
| 54407 | SLC38A2 | -0.36595897 | 0.003156108 | 0.15735649 | 2.500848144 |
| 23198 | PSME4 | -0.375574124 | 0.000363605 | 0.054627364 | 3.439370153 |
| 100861548 | PINK1-AS | -0.376988521 | 0.001240477 | 0.096238624 | 2.906411284 |
| 167227 | DCP2 | -0.377911635 | 0.000480505 | 0.061633976 | 3.318302089 |
| 7884 | SLBP | -0.380067181 | 0.003008486 | 0.155404722 | 2.521652005 |
| 340371 | NRBP2 | -0.383096803 | 0.008054226 | 0.240230816 | 2.093976188 |
| 25822 | DNAJB5 | -0.387194738 | 0.002150033 | 0.129343632 | 2.667554874 |
| 7517 | XRCC3 | -0.389796341 | 0.004733492 | 0.189573334 | 2.324818353 |
| 2184 | FAH | -0.39590193 | 0.000514705 | 0.062971482 | 3.288441613 |
| 163126 | EID2 | -0.40028444 | 0.007497586 | 0.231658272 | 2.125078544 |
| 51715 | RAB23 | -0.401256766 | 0.000591576 | 0.066407614 | 3.227989453 |
| 54819 | ZCCHC10 | -0.402859289 | 0.002704307 | 0.148438915 | 2.567944008 |
| 373 | TRIM23 | -0.405256735 | 0.00144452 | 0.103402233 | 2.840276441 |
| 100505549 | LOC100505549 | -0.414861682 | 0.005342873 | 0.198574459 | 2.272225149 |
| 375743 | PTAR1 | -0.425581891 | 0.003294999 | 0.159661255 | 2.482144713 |
| 113 | ADCY7 | -0.428542491 | 0.000394127 | 0.056320771 | 3.404363812 |
| 3682 | ITGAE | -0.431066261 | 0.001316545 | 0.098111059 | 2.880564292 |
| 10769 | PLK2 | -0.432020218 | 0.002441834 | 0.138119259 | 2.612283863 |
| 5214 | PFKP | -0.432975803 | 0.001793421 | 0.117336838 | 2.746317749 |
| 154743 | BMT2 | -0.437686931 | 0.000489465 | 0.062046189 | 3.310278358 |
| 29954 | POMT2 | -0.437751849 | 0.000303702 | 0.051027253 | 3.517552348 |
| 162993 | ZNF846 | -0.438993515 | 0.005941805 | 0.208550747 | 2.226081605 |
| 662 | BNIP1 | -0.439542472 | 0.001876035 | 0.119439462 | 2.726759064 |
| 27010 | TPK1 | -0.439892431 | 0.000450433 | 0.060616288 | 3.346369799 |
| 2971 | GTF3A | -0.444417821 | 0.00915487 | 0.254356697 | 2.038347818 |
| 26999 | CYFIP2 | -0.445608587 | 0.006464745 | 0.215898284 | 2.189448601 |
| 55214 | P3H2 | -0.44726029 | 0.005033089 | 0.193370476 | 2.29816539 |
| 8034 | SLC25A16 | -0.449779032 | 0.005884636 | 0.207828655 | 2.230280396 |
| 26751 | SH3YL1 | -0.466264884 | 0.005403609 | 0.199813058 | 2.267316084 |
| 65055 | REEP1 | -0.468913719 | 0.001861044 | 0.119252776 | 2.730243359 |
| 56180 | MOSPD1 | -0.483668341 | 0.001627249 | 0.111261127 | 2.788545987 |
| 115708 | TRMT61A | -0.483951468 | 0.00671392 | 0.220118925 | 2.173023838 |
| 54504 | CPVL | -0.503701965 | 0.004729241 | 0.189573334 | 2.325208554 |
| 91749 | MFSD4B | -0.503748042 | 0.007814927 | 0.236068996 | 2.107075074 |
| 81609 | SNX27 | -0.505173022 | 0.000714904 | 0.070978013 | 3.145752273 |
| 19 | ABCA1 | -0.508094031 | 0.002202469 | 0.131890902 | 2.657090196 |
| 84938 | ATG4C | -0.509368571 | 0.003662492 | 0.169375324 | 2.436223315 |
| 170959 | ZNF431 | -0.515331475 | 0.000693981 | 0.069552812 | 3.15865242 |
| 80176 | SPSB1 | -0.524661659 | 0.002367548 | 0.136086096 | 2.625701207 |
| 1902 | LPAR1 | -0.524829824 | 0.000455983 | 0.061073821 | 3.341051348 |
| 11169 | WDHD1 | -0.528893491 | 0.002839119 | 0.152107164 | 2.546816404 |
| 100507487 | LINC02615 | -0.559092934 | 0.00062512 | 0.06753908 | 3.204036606 |
| 51776 | MAP3K20 | -0.565898456 | 0.008282961 | 0.24322096 | 2.081814383 |
| 10810 | WASF3 | -0.567405146 | 0.000383399 | 0.056193108 | 3.416349024 |
| 10825 | NEU3 | -0.579775248 | 0.002674949 | 0.147485795 | 2.572684494 |
| 150967 | LINC01963 | -0.585433263 | 0.008251632 | 0.24255184 | 2.083460149 |
| 55839 | CENPN | -0.588209513 | 0.005574716 | 0.203003874 | 2.253777253 |
| 6671 | SP4 | -0.588241431 | 0.000607254 | 0.0668193 | 3.216629616 |
| 4649 | MYO9A | -0.594645237 | 0.002906839 | 0.153571991 | 2.536579022 |
| 2013 | EMP2 | -0.597049892 | 0.000385901 | 0.056193108 | 3.413524096 |
| 857 | CAV1 | -0.620546725 | 0.000553146 | 0.065155448 | 3.257160224 |
| 3070 | HELLS | -0.629876541 | 0.002855938 | 0.15243302 | 2.544251225 |
| 2697 | GJA1 | -0.640074014 | 0.001354771 | 0.0994024 | 2.868134108 |
| 8660 | IRS2 | -0.658916327 | 0.005468748 | 0.201146497 | 2.262112088 |
| 166336 | PRICKLE2 | -0.666745123 | 0.00148747 | 0.105066449 | 2.827551785 |
| 90952 | ESAM | -0.679084517 | 0.001445701 | 0.103402233 | 2.839921519 |
| 55283 | MCOLN3 | -0.722119225 | 0.002548148 | 0.141871895 | 2.593775351 |
| 94240 | EPSTI1 | -0.725123851 | 0.000553228 | 0.065155448 | 3.257095847 |
| 11010 | GLIPR1 | -0.754926664 | 0.000804297 | 0.075622593 | 3.094583551 |
| 116931 | MED12L | -0.827039852 | 0.005313993 | 0.198019459 | 2.274579022 |
| 101927230 | LINC01969 | -0.86920057 | 0.004457769 | 0.18559875 | 2.35088244 |
| 11279 | KLF8 | -0.899606483 | 0.008898777 | 0.251546418 | 2.050669676 |
| 3400 | ID4 | -0.912907498 | 0.008343191 | 0.243980335 | 2.078667814 |
| 114821 | ZBED9 | -0.95557866 | 0.002395721 | 0.136874237 | 2.62056376 |
| 83887 | TTLL2 | -0.967753666 | 0.000931965 | 0.082183696 | 3.030600397 |
| 59345 | GNB4 | -0.977438911 | 0.000545352 | 0.064791918 | 3.26332309 |
| 60468 | BACH2 | -1.073278955 | 0.002051227 | 0.124720721 | 2.687986276 |
| 139221 | PWWP3B | -1.091468652 | 0.001503846 | 0.105755919 | 2.822796635 |
| 654429 | LRTM2 | -1.116991802 | 0.001379015 | 0.099890634 | 2.86043101 |
| 100302112 | MIR1284 | -1.116991802 | 0.001379015 | 0.099890634 | 2.86043101 |
| 102465857 | MIR7976 | -1.116991802 | 0.001379015 | 0.099890634 | 2.86043101 |
| 692149 | SCARNA14 | -1.116991802 | 0.001379015 | 0.099890634 | 2.86043101 |
| 651746 | ANKRD33B | -1.122523977 | 0.000787122 | 0.075507828 | 3.103957949 |
| 50507 | NOX4 | -1.161779871 | 0.00391546 | 0.1761957 | 2.407217208 |
| 115207 | KCTD12 | -1.211680641 | 0.003839759 | 0.174201182 | 2.415696033 |
| 100128338 | NA (ENTREZID: 100128338) | -1.302296039 | 0.001301197 | 0.097969659 | 2.885656947 |
| 387700 | SLC16A12 | -1.440205848 | 0.000918713 | 0.082183696 | 3.036820138 |
| 100287072 | LOC100287072 | -1.510883176 | 0.001256507 | 0.096747942 | 2.900835088 |
| 26525 | IL36RN | -1.781611515 | 0.009383055 | 0.256385482 | 2.027655738 |
| 100128055 | SMARCA5-AS1 | -1.922561507 | 0.00092916 | 0.082183696 | 3.031909495 |
| 1215 | CMA1 | -2.002564526 | 0.002762656 | 0.150278934 | 2.558673189 |
| 100507524 | ARHGEF26-AS1 | -2.293108849 | 0.004470328 | 0.185633035 | 2.34966061 |
| 9350 | CER1 | -2.330790007 | 0.003678319 | 0.169830684 | 2.43435061 |
| 54798 | DCHS2 | -3.383110676 | 0.000339952 | 0.053038153 | 3.468582399 |
| 55051 | NRDE2 | -0.000882436 | 0.99027764 | 0.995591924 | 0.004243027 |
| 51201 | ZDHHC2 | -0.004113548 | 0.964365272 | 0.984643715 | 0.015758438 |
| 9797 | TATDN2 | -0.009663214 | 0.888675147 | 0.950703108 | 0.051256965 |
| 10491 | CRTAP | -0.011020931 | 0.888952682 | 0.950703108 | 0.051121355 |
| 28968 | SLC6A16 | -0.013256073 | 0.937107957 | 0.97210257 | 0.028210375 |
| 9603 | NFE2L3 | -0.01395963 | 0.872434315 | 0.944156276 | 0.059267261 |
| 2512 | FTL | -0.014947796 | 0.906197222 | 0.95983026 | 0.042777274 |
| 5465 | PPARA | -0.017878225 | 0.798994907 | 0.908145643 | 0.097455989 |
| 646300 | COL6A4P2 | -0.018061579 | 0.95801597 | 0.982088287 | 0.018627251 |
| 140606 | SELENOM | -0.020486325 | 0.88651163 | 0.949996954 | 0.052315563 |
| 646471 | LOC646471 | -0.021792067 | 0.838915634 | 0.928187711 | 0.076281712 |
| 51236 | HGH1 | -0.027031039 | 0.774780256 | 0.896132569 | 0.110821455 |
| 2033 | EP300 | -0.028974773 | 0.538882531 | 0.773064029 | 0.268505895 |
| 222255 | ATXN7L1 | -0.03013737 | 0.756718035 | 0.887782862 | 0.121065915 |
| 4953 | ODC1 | -0.03065946 | 0.784606127 | 0.901286095 | 0.105348305 |
| 64784 | CRTC3 | -0.035348673 | 0.603914189 | 0.812906901 | 0.219024766 |
| 2889 | RAPGEF1 | -0.036763827 | 0.453783947 | 0.723146442 | 0.343150872 |
| 26608 | TBL2 | -0.039815084 | 0.567576457 | 0.792186547 | 0.245975627 |
| 50485 | SMARCAL1 | -0.041499593 | 0.715052103 | 0.867651146 | 0.145662312 |
| 55751 | TMEM184C | -0.043954489 | 0.363691086 | 0.645720528 | 0.439267343 |
| 10016 | PDCD6 | -0.045117452 | 0.322737991 | 0.601519216 | 0.491149909 |
| 143 | PARP4 | -0.04523702 | 0.440559796 | 0.71471742 | 0.355995138 |
| 64802 | NMNAT1 | -0.045817628 | 0.691002874 | 0.856929008 | 0.160520146 |
| 29803 | REPIN1 | -0.046352235 | 0.406993762 | 0.686217437 | 0.390412247 |
| 9497 | SLC4A7 | -0.04822344 | 0.576537077 | 0.7961465 | 0.239172758 |
| 54537 | SHLD2 | -0.051180481 | 0.463476388 | 0.728471565 | 0.333972386 |
| 8932 | MBD2 | -0.053002091 | 0.309366258 | 0.586647181 | 0.509527056 |
| 91768 | CABLES1 | -0.053630561 | 0.481244825 | 0.740736022 | 0.317633928 |
| 902 | CCNH | -0.056439718 | 0.442522963 | 0.716101871 | 0.354064188 |
| 152302 | CIDECP1 | -0.058028058 | 0.425468174 | 0.702120733 | 0.371132921 |
| 3097 | HIVEP2 | -0.06060758 | 0.591030389 | 0.805842359 | 0.228390188 |
| 8480 | RAE1 | -0.061031931 | 0.306396296 | 0.583216064 | 0.513716489 |
| 63941 | NECAB3 | -0.06632929 | 0.598364917 | 0.810657561 | 0.223033878 |
| 79813 | EHMT1 | -0.06857105 | 0.136264096 | 0.498711659 | 0.865618561 |
| 29925 | GMPPB | -0.068992647 | 0.469398096 | 0.731614829 | 0.328458676 |
| 9024 | BRSK2 | -0.069383181 | 0.732227271 | 0.875569501 | 0.1353541 |
| 7132 | TNFRSF1A | -0.069611043 | 0.354554316 | 0.636396175 | 0.450317224 |
| 54974 | THG1L | -0.069740332 | 0.590684662 | 0.805637091 | 0.228644306 |
| 49855 | SCAPER | -0.072951096 | 0.372154081 | 0.649776495 | 0.429277214 |
| 60312 | AFAP1 | -0.073684837 | 0.355598444 | 0.63745062 | 0.449040148 |
| 114885 | OSBPL11 | -0.073896713 | 0.256227649 | 0.525009676 | 0.591374008 |
| 26015 | RPAP1 | -0.074568147 | 0.42593473 | 0.702469457 | 0.370656947 |
| 4306 | NR3C2 | -0.080280179 | 0.702087177 | 0.862000383 | 0.153608959 |
| 80131 | LRRC8E | -0.08093721 | 0.354868972 | 0.636727507 | 0.449931972 |
| 79026 | AHNAK | -0.080972459 | 0.337193247 | 0.618115058 | 0.472121132 |
| 5893 | RAD52 | -0.081711452 | 0.507856614 | 0.755319933 | 0.294258887 |
| 10985 | GCN1 | -0.081733887 | 0.320886125 | 0.59936002 | 0.493649061 |
| 26521 | TIMM8B | -0.083378258 | 0.117404147 | 0.498711659 | 0.930316562 |
| 146330 | FBXL16 | -0.083796592 | 0.7893648 | 0.903712891 | 0.102722244 |
| 23135 | KDM6B | -0.084268514 | 0.422398165 | 0.69960312 | 0.374277977 |
| 100505696 | SH3BP5-AS1 | -0.084558319 | 0.584748707 | 0.80155428 | 0.23303073 |
| 55257 | MRGBP | -0.08602033 | 0.30510452 | 0.581868685 | 0.515551358 |
| 253143 | PRR14L | -0.086612265 | 0.229703623 | 0.498711659 | 0.638832155 |
| 65117 | RSRC2 | -0.086823774 | 0.19895059 | 0.498711659 | 0.701254769 |
| 5139 | PDE3A | -0.086969231 | 0.732998833 | 0.875915405 | 0.134896717 |
| 63892 | THADA | -0.087098945 | 0.265108185 | 0.535643085 | 0.576576864 |
| 1387 | CREBBP | -0.088472491 | 0.265899492 | 0.53654179 | 0.575282492 |
| 9950 | GOLGA5 | -0.088700483 | 0.246600386 | 0.515134109 | 0.608006248 |
| 3658 | IREB2 | -0.089571403 | 0.057153096 | 0.498711659 | 1.242960239 |
| 26122 | EPC2 | -0.089960299 | 0.318203948 | 0.596318542 | 0.497294436 |
| 8729 | GBF1 | -0.090851669 | 0.086334072 | 0.498711659 | 1.063817775 |
| 11171 | STRAP | -0.091085542 | 0.054367898 | 0.496013172 | 1.264657458 |
| 30968 | STOML2 | -0.092019619 | 0.131063806 | 0.498711659 | 0.882517225 |
| 9801 | MRPL19 | -0.092316621 | 0.241373607 | 0.508178511 | 0.61731022 |
| 7469 | NELFA | -0.092732804 | 0.161843713 | 0.498711659 | 0.790904167 |
| 79882 | ZC3H14 | -0.093312057 | 0.190641654 | 0.498711659 | 0.719782203 |
| 84181 | CHD6 | -0.093484076 | 0.398785299 | 0.678223098 | 0.39926086 |
| 6341 | SCO1 | -0.093862162 | 0.099835189 | 0.498711659 | 1.000716356 |
| 9040 | UBE2M | -0.094225348 | 0.112329976 | 0.498711659 | 0.949504334 |
| 29982 | NRBF2 | -0.094498612 | 0.059557878 | 0.498711659 | 1.225060784 |
| 399818 | EEF1AKMT2 | -0.095058727 | 0.402900163 | 0.682354177 | 0.394802557 |
| 54849 | DEF8 | -0.095444497 | 0.117695233 | 0.498711659 | 0.929241127 |
| 22794 | CASC3 | -0.095554495 | 0.148898747 | 0.498711659 | 0.827108957 |
| 4836 | NMT1 | -0.095784465 | 0.10656671 | 0.498711659 | 0.972378442 |
| 1829 | DSG2 | -0.096691296 | 0.208947802 | 0.498711659 | 0.679962193 |
| 79637 | ARMC7 | -0.097449914 | 0.27515194 | 0.546818262 | 0.560427421 |
| 57496 | MRTFB | -0.098820282 | 0.210006479 | 0.498711659 | 0.677767306 |
| 55314 | TMEM144 | -0.099178082 | 0.306934569 | 0.583946874 | 0.512954196 |
| 3980 | LIG3 | -0.100616589 | 0.309478952 | 0.586741158 | 0.509368882 |
| 55854 | ZC3H15 | -0.101075794 | 0.110616416 | 0.498711659 | 0.956180417 |
| 51663 | ZFR | -0.101878764 | 0.073863334 | 0.498711659 | 1.131571093 |
| 81554 | RCC1L | -0.102743861 | 0.366594638 | 0.646156072 | 0.435813892 |
| 7337 | UBE3A | -0.102767324 | 0.055230055 | 0.49843982 | 1.257824524 |
| 11157 | LSM6 | -0.105358513 | 0.305860304 | 0.582722983 | 0.514476884 |
| 8539 | API5 | -0.10641797 | 0.08452375 | 0.498711659 | 1.073021243 |
| 9373 | PLAA | -0.107488281 | 0.020216696 | 0.356832556 | 1.694289819 |
| 52 | ACP1 | -0.107527348 | 0.050693755 | 0.493681625 | 1.295045538 |
| 51406 | NOL7 | -0.109176813 | 0.079802592 | 0.498711659 | 1.097983002 |
| 54778 | RNF111 | -0.1092955 | 0.275935653 | 0.547597267 | 0.559192182 |
| 1352 | COX10 | -0.109799864 | 0.177555424 | 0.498711659 | 0.750666056 |
| 3735 | KARS1 | -0.109815902 | 0.152695739 | 0.498711659 | 0.816173082 |
| 57466 | SCAF4 | -0.110943615 | 0.086666877 | 0.498711659 | 1.062146853 |
| 26984 | SEC22A | -0.111267802 | 0.30955101 | 0.586741158 | 0.509267775 |
| 6883 | TAF12 | -0.113895321 | 0.181453562 | 0.498711659 | 0.741234502 |
| 83605 | CCM2 | -0.114445559 | 0.232674867 | 0.498711659 | 0.633250526 |
| 53834 | FGFRL1 | -0.114568024 | 0.13319724 | 0.498711659 | 0.875504774 |
| 136647 | MPLKIP | -0.1145929 | 0.278842293 | 0.55106674 | 0.554641355 |
| 55573 | CDV3 | -0.115280369 | 0.130611041 | 0.498711659 | 0.884020109 |
| 101928141 | LINC01028 | -0.11562024 | 0.610604027 | 0.81600003 | 0.214240336 |
| 2130 | EWSR1 | -0.116153059 | 0.094619654 | 0.498711659 | 1.024018644 |
| 10209 | EIF1 | -0.117243667 | 0.036844846 | 0.453685651 | 1.433623254 |
| 81608 | FIP1L1 | -0.117360902 | 0.137134164 | 0.498711659 | 0.862854337 |
| 30849 | PIK3R4 | -0.117465132 | 0.085012535 | 0.498711659 | 1.070517033 |
| 4236 | MFAP1 | -0.118958087 | 0.057943778 | 0.498711659 | 1.236993192 |
| 8455 | ATRN | -0.119563292 | 0.144737342 | 0.498711659 | 0.839419407 |
| 114971 | PTPMT1 | -0.119723944 | 0.11468292 | 0.498711659 | 0.940501258 |
| 9166 | EBAG9 | -0.120670752 | 0.153916004 | 0.498711659 | 0.81271622 |
| 23339 | VPS39 | -0.122054368 | 0.135585872 | 0.498711659 | 0.867785561 |
| 64762 | GAREM1 | -0.122338435 | 0.606241007 | 0.812911475 | 0.217354691 |
| 23645 | PPP1R15A | -0.122574778 | 0.418007919 | 0.695822187 | 0.378815491 |
| 54148 | MRPL39 | -0.122592522 | 0.146351835 | 0.498711659 | 0.834601828 |
| 125206 | SLC5A10 | -0.123439643 | 0.177683325 | 0.498711659 | 0.750353327 |
| 387893 | KMT5A | -0.124203557 | 0.050987905 | 0.493681625 | 1.292532832 |
| 5356 | PLRG1 | -0.124575298 | 0.135207442 | 0.498711659 | 0.868999404 |
| 9666 | DZIP3 | -0.125657275 | 0.217266262 | 0.498711659 | 0.663007707 |
| 5754 | PTK7 | -0.125680254 | 0.062623107 | 0.498711659 | 1.203265389 |
| 64689 | GORASP1 | -0.128495583 | 0.101576156 | 0.498711659 | 0.993208226 |
| 85395 | FAM207A | -0.129396804 | 0.257869053 | 0.526944405 | 0.588600775 |
| 4152 | MBD1 | -0.1294055 | 0.017805407 | 0.337716862 | 1.749448095 |
| 203069 | R3HCC1 | -0.130058295 | 0.321323231 | 0.599787874 | 0.493057875 |
| 8664 | EIF3D | -0.130687396 | 0.034113948 | 0.439382009 | 1.467068017 |
| 6730 | SRP68 | -0.131833703 | 0.101669162 | 0.498711659 | 0.992810756 |
| 64122 | FN3K | -0.132389797 | 0.521163416 | 0.76364923 | 0.283026078 |
| 23001 | WDFY3 | -0.132719055 | 0.120404304 | 0.498711659 | 0.919357988 |
| 55197 | RPRD1A | -0.133077678 | 0.20418166 | 0.498711659 | 0.68998327 |
| 5007 | OSBP | -0.134382816 | 0.07982112 | 0.498711659 | 1.097882183 |
| 6921 | ELOC | -0.134412765 | 0.023422536 | 0.375753064 | 1.630366085 |
| 85364 | ZCCHC3 | -0.134956508 | 0.072982818 | 0.498711659 | 1.136779372 |
| 3608 | ILF2 | -0.135280447 | 0.02989459 | 0.417048939 | 1.524407398 |
| 7336 | UBE2V2 | -0.136371567 | 0.012095312 | 0.288126156 | 1.917382924 |
| 10427 | SEC24B | -0.140290111 | 0.080839672 | 0.498711659 | 1.092375457 |
| 51194 | IPO11 | -0.140803403 | 0.247203466 | 0.515900516 | 0.606945444 |
| 3836 | KPNA1 | -0.141052261 | 0.023852909 | 0.377550077 | 1.622458649 |
| 8621 | CDK13 | -0.141116166 | 0.026833093 | 0.398846075 | 1.571329264 |
| 221079 | ARL5B | -0.141153796 | 0.411065963 | 0.689350983 | 0.386088482 |
| 5718 | PSMD12 | -0.141213716 | 0.030512236 | 0.420580071 | 1.515525965 |
| 48 | ACO1 | -0.141229234 | 0.30080947 | 0.577133277 | 0.521708496 |
| 1399 | CRKL | -0.142510518 | 0.051146833 | 0.493681625 | 1.291181253 |
| 79109 | MAPKAP1 | -0.142661658 | 0.034736768 | 0.441715419 | 1.459210592 |
| 29914 | UBIAD1 | -0.142865334 | 0.320218484 | 0.598593624 | 0.494553603 |
| 54708 | MARCHF5 | -0.143004263 | 0.129103212 | 0.498711659 | 0.889062953 |
| 2140 | EYA3 | -0.143168308 | 0.10514082 | 0.498711659 | 0.97822864 |
| 22941 | SHANK2 | -0.144707608 | 0.182405734 | 0.498711659 | 0.738961514 |
| 123720 | WHAMM | -0.14488227 | 0.123380946 | 0.498711659 | 0.908751904 |
| 130074 | FAM168B | -0.146819786 | 0.071907399 | 0.498711659 | 1.14322642 |
| 59338 | PLEKHA1 | -0.14691631 | 0.053908375 | 0.494444092 | 1.268343759 |
| 8846 | ALKBH1 | -0.147741506 | 0.212604745 | 0.498711659 | 0.672427047 |
| 9330 | GTF3C3 | -0.148211067 | 0.196198863 | 0.498711659 | 0.707303514 |
| 25 | ABL1 | -0.148380972 | 0.067373135 | 0.498711659 | 1.171513244 |
| 79871 | RPAP2 | -0.150190909 | 0.239863197 | 0.50597396 | 0.620036382 |
| 8732 | RNGTT | -0.152310995 | 0.089195867 | 0.498711659 | 1.049655269 |
| 10273 | STUB1 | -0.153150532 | 0.084189441 | 0.498711659 | 1.074742374 |
| 9716 | AQR | -0.153432285 | 0.036408568 | 0.450466788 | 1.438796402 |
| 55181 | SMG8 | -0.154543823 | 0.112529714 | 0.498711659 | 0.948732785 |
| 1434 | CSE1L | -0.154975567 | 0.035350794 | 0.444228985 | 1.451600827 |
| 8803 | SUCLA2 | -0.155465485 | 0.049283668 | 0.493681625 | 1.307296977 |
| 28512 | NKIRAS1 | -0.15657496 | 0.088857183 | 0.498711659 | 1.051307459 |
| 3308 | HSPA4 | -0.156739596 | 0.08525314 | 0.498711659 | 1.069289616 |
| 4928 | NUP98 | -0.157264848 | 0.02248328 | 0.371584146 | 1.648140331 |
| 9732 | DOCK4 | -0.157764948 | 0.290762342 | 0.56495119 | 0.536461842 |
| 1810 | DR1 | -0.157920865 | 0.06921625 | 0.498711659 | 1.159791934 |
| 27229 | TUBGCP4 | -0.157945782 | 0.136571648 | 0.498711659 | 0.86463945 |
| 28652 | TRAV30 | -0.15815423 | 0.698345292 | 0.86078016 | 0.15592979 |
| 51164 | DCTN4 | -0.158360981 | 0.022400496 | 0.371098064 | 1.649742365 |
| 55005 | RMND1 | -0.160246534 | 0.131278324 | 0.498711659 | 0.881806976 |
| 283149 | BCL9L | -0.160632563 | 0.015882975 | 0.323454888 | 1.799068148 |
| 85403 | EAF1 | -0.161392551 | 0.040980972 | 0.474573692 | 1.387417745 |
| 57688 | ZSWIM6 | -0.161441127 | 0.020900166 | 0.359672853 | 1.679850264 |
| 115752 | DIS3L | -0.165309359 | 0.119001505 | 0.498711659 | 0.924447546 |
| 132660 | LIN54 | -0.165766087 | 0.019363894 | 0.348968345 | 1.713007303 |
| 51010 | EXOSC3 | -0.167758949 | 0.059724963 | 0.498711659 | 1.223844111 |
| 84321 | THOC3 | -0.168094088 | 0.255944855 | 0.524777563 | 0.591853596 |
| 904 | CCNT1 | -0.168707121 | 0.019061316 | 0.347348646 | 1.719847119 |
| 4686 | NCBP1 | -0.1688976 | 0.026083662 | 0.393103921 | 1.583631436 |
| 5187 | PER1 | -0.169208483 | 0.200013508 | 0.498711659 | 0.698940673 |
| 6400 | SEL1L | -0.169610123 | 0.045481853 | 0.493681625 | 1.34216185 |
| 9895 | TECPR2 | -0.170289362 | 0.202583186 | 0.498711659 | 0.693396603 |
| 22908 | SACM1L | -0.170652951 | 0.066724353 | 0.498711659 | 1.175715629 |
| 497258 | BDNF-AS | -0.1712297 | 0.518890852 | 0.762032881 | 0.284923986 |
| 169792 | GLIS3 | -0.171732154 | 0.037459359 | 0.457487527 | 1.426439658 |
| 79039 | DDX54 | -0.171734203 | 0.129655793 | 0.498711659 | 0.887208074 |
| 2054 | STX2 | -0.17217807 | 0.036075542 | 0.448103684 | 1.442787135 |
| 57474 | ZNF490 | -0.17249889 | 0.175473114 | 0.498711659 | 0.755789417 |
| 22862 | FNDC3A | -0.173116649 | 0.074369081 | 0.498711659 | 1.128607585 |
| 83737 | ITCH | -0.173438665 | 0.06801736 | 0.498711659 | 1.167380229 |
| 23165 | NUP205 | -0.174218177 | 0.086755975 | 0.498711659 | 1.061700605 |
| 51295 | ECSIT | -0.174288869 | 0.204566817 | 0.498711659 | 0.689164812 |
| 51622 | CCZ1 | -0.174408076 | 0.065009609 | 0.498711659 | 1.187022446 |
| 9169 | SCAF11 | -0.176375756 | 0.019575589 | 0.350914681 | 1.708285162 |
| 29889 | GNL2 | -0.176574737 | 0.033387899 | 0.436387242 | 1.476410909 |
| 6882 | TAF11 | -0.176980464 | 0.039725042 | 0.469324524 | 1.400935635 |
| 79658 | ARHGAP10 | -0.177538657 | 0.018672163 | 0.344643459 | 1.72880537 |
| 4616 | GADD45B | -0.178690963 | 0.04621157 | 0.493681625 | 1.335249276 |
| 130507 | UBR3 | -0.182707051 | 0.015647013 | 0.321255914 | 1.805568557 |
| 55022 | PID1 | -0.182935847 | 0.044604704 | 0.491737288 | 1.350619338 |
| 79697 | RIOX1 | -0.183107511 | 0.044627821 | 0.491737288 | 1.350394318 |
| 56947 | MFF | -0.183950833 | 0.085103775 | 0.498711659 | 1.070051175 |
| 84280 | BTBD10 | -0.183984962 | 0.018562628 | 0.34397431 | 1.731360539 |
| 9441 | MED26 | -0.184215513 | 0.067040946 | 0.498711659 | 1.173659866 |
| 152185 | SPICE1 | -0.185260436 | 0.116046373 | 0.498711659 | 0.935368429 |
| 91833 | WDR20 | -0.185389793 | 0.013795035 | 0.303181134 | 1.860277193 |
| 2060 | EPS15 | -0.185509467 | 0.011387689 | 0.28048456 | 1.943564402 |
| 5771 | PTPN2 | -0.185867652 | 0.013680256 | 0.302061328 | 1.863905776 |
| 285855 | RPL7L1 | -0.187563445 | 0.045687848 | 0.493681625 | 1.340199298 |
| 63894 | VIPAS39 | -0.188057846 | 0.108945365 | 0.498711659 | 0.962791242 |
| 23568 | ARL2BP | -0.188376582 | 0.036860461 | 0.453685651 | 1.433439238 |
| 27230 | SERP1 | -0.189047927 | 0.056639567 | 0.498711659 | 1.246880075 |
| 55076 | TMEM45A | -0.189591638 | 0.139144226 | 0.498711659 | 0.856534811 |
| 9056 | SLC7A7 | -0.18965206 | 0.046510341 | 0.493681625 | 1.332450476 |
| 28977 | MRPL42 | -0.189900668 | 0.029272198 | 0.413730247 | 1.533544666 |
| 60592 | SCOC | -0.190354177 | 0.101774139 | 0.498711659 | 0.992362563 |
| 64422 | ATG3 | -0.190600473 | 0.013086408 | 0.297030712 | 1.883179544 |
| 5533 | PPP3CC | -0.191859752 | 0.17734849 | 0.498711659 | 0.751172505 |
| 5062 | PAK2 | -0.192914326 | 0.014478275 | 0.308670759 | 1.839283179 |
| 114 | ADCY8 | -0.194385348 | 0.725710266 | 0.87249314 | 0.139236733 |
| 2176 | FANCC | -0.195015934 | 0.054589681 | 0.497179132 | 1.262889444 |
| 85460 | ZNF518B | -0.196194815 | 0.016188648 | 0.327172 | 1.79078942 |
| 54764 | ZRANB1 | -0.19874143 | 0.016738095 | 0.331256573 | 1.776293972 |
| 339230 | CCDC137 | -0.198861279 | 0.098458529 | 0.498711659 | 1.006746657 |
| 104548973 | SALRNA2 | -0.20094027 | 0.437047395 | 0.71152962 | 0.359471464 |
| 7247 | TSN | -0.201147882 | 0.044313917 | 0.491137265 | 1.35345986 |
| 55750 | AGK | -0.201368443 | 0.235832532 | 0.500932432 | 0.627396286 |
| 4528 | MTIF2 | -0.202023883 | 0.130922988 | 0.498711659 | 0.882984092 |
| 747 | DAGLA | -0.202886494 | 0.154079272 | 0.498711659 | 0.812255782 |
| 54455 | FBXO42 | -0.203174146 | 0.012925139 | 0.296214143 | 1.888564778 |
| 64651 | CSRNP1 | -0.205268032 | 0.051932819 | 0.493681625 | 1.284558103 |
| 51433 | ANAPC5 | -0.205307665 | 0.011730667 | 0.284451148 | 1.930677293 |
| 90 | ACVR1 | -0.206239853 | 0.010331706 | 0.268064651 | 1.985827961 |
| 83940 | TATDN1 | -0.206366187 | 0.019108993 | 0.347597594 | 1.718762199 |
| 79023 | NUP37 | -0.207006526 | 0.219555256 | 0.498711659 | 0.658456162 |
| 54834 | GDAP2 | -0.207888456 | 0.016343911 | 0.32809721 | 1.786644011 |
| 203427 | SLC25A43 | -0.208280674 | 0.010469852 | 0.269762522 | 1.980059457 |
| 134492 | NUDCD2 | -0.208763384 | 0.027541477 | 0.40289712 | 1.560012773 |
| 2958 | GTF2A2 | -0.209687498 | 0.016818763 | 0.331414825 | 1.774205949 |
| 10492 | SYNCRIP | -0.210808354 | 0.023984703 | 0.378779555 | 1.620065655 |
| 204851 | HIPK1 | -0.212079464 | 0.017711468 | 0.337528275 | 1.751745441 |
| 79753 | SNIP1 | -0.213300606 | 0.010063528 | 0.26477537 | 1.997249741 |
| 8458 | TTF2 | -0.213649128 | 0.043384935 | 0.484833829 | 1.362661049 |
| 55596 | ZCCHC8 | -0.214903063 | 0.01081975 | 0.273624835 | 1.965782774 |
| 8495 | PPFIBP2 | -0.218918428 | 0.016583944 | 0.329762676 | 1.780312177 |
| 57213 | SPRYD7 | -0.219647278 | 0.073666729 | 0.498711659 | 1.132728614 |
| 10450 | PPIE | -0.219829249 | 0.034840154 | 0.44223789 | 1.457919934 |
| 51397 | COMMD10 | -0.220061902 | 0.033875423 | 0.439042933 | 1.470115273 |
| 55605 | KIF21A | -0.221412908 | 0.013549204 | 0.300791879 | 1.868086218 |
| 9764 | KIAA0513 | -0.221428626 | 0.016179719 | 0.327172 | 1.791029025 |
| 9941 | EXOG | -0.221886276 | 0.022327503 | 0.370104757 | 1.651159844 |
| 2553 | GABPB1 | -0.223289325 | 0.01198181 | 0.28728883 | 1.921477571 |
| 10395 | DLC1 | -0.223755334 | 0.060652022 | 0.498711659 | 1.217154716 |
| 10771 | ZMYND11 | -0.22422279 | 0.060859158 | 0.498711659 | 1.21567406 |
| 10949 | HNRNPA0 | -0.227249221 | 0.01232769 | 0.29008361 | 1.909118295 |
| 5984 | RFC4 | -0.230141295 | 0.024554336 | 0.381055674 | 1.609871806 |
| 10927 | SPIN1 | -0.230260778 | 0.012446008 | 0.290898875 | 1.904969924 |
| 60487 | TRMT11 | -0.230987099 | 0.019635843 | 0.351772713 | 1.706950449 |
| 7343 | UBTF | -0.231798383 | 0.037800604 | 0.460269357 | 1.422501261 |
| 8898 | MTMR2 | -0.232672913 | 0.012372354 | 0.290268247 | 1.907547662 |
| 285352 | KIF9-AS1 | -0.234422122 | 0.154339176 | 0.498711659 | 0.811523823 |
| 10389 | SCML2 | -0.23525584 | 0.088734002 | 0.498711659 | 1.051909931 |
| 100288428 | LMCD1-AS1 | -0.236617971 | 0.705528301 | 0.863251437 | 0.151485561 |
| 11275 | KLHL2 | -0.237762999 | 0.044873063 | 0.492781862 | 1.348014285 |
| 147339 | C18orf25 | -0.239520803 | 0.03304916 | 0.435178498 | 1.480839574 |
| 6117 | RPA1 | -0.240074932 | 0.10701187 | 0.498711659 | 0.970568047 |
| 54892 | NCAPG2 | -0.240151716 | 0.020566298 | 0.35878954 | 1.686843876 |
| 64852 | TUT1 | -0.240286981 | 0.12289007 | 0.498711659 | 0.910483208 |
| 54529 | ASNSD1 | -0.241064618 | 0.011016729 | 0.276293147 | 1.957947334 |
| 9812 | DELE1 | -0.241990428 | 0.018473769 | 0.343365893 | 1.733444491 |
| 140710 | SOGA1 | -0.24205781 | 0.116132437 | 0.498711659 | 0.93504646 |
| 401613 | SERTM2 | -0.243248326 | 0.439513318 | 0.713915009 | 0.357027961 |
| 126129 | CPT1C | -0.243592567 | 0.325869603 | 0.605369145 | 0.486956148 |
| 7110 | TMF1 | -0.243955491 | 0.055734085 | 0.498711659 | 1.253879124 |
| 9097 | USP14 | -0.244143124 | 0.01034634 | 0.268064651 | 1.985213254 |
| 10116 | FEM1B | -0.246024827 | 0.010358194 | 0.268064651 | 1.984715959 |
| 9128 | PRPF4 | -0.246518995 | 0.020218048 | 0.356832556 | 1.694260777 |
| 340554 | ZC3H12B | -0.246768309 | 0.29026018 | 0.564400512 | 0.53721254 |
| 154467 | CCDC167 | -0.249461188 | 0.069943387 | 0.498711659 | 1.155253341 |
| 22881 | ANKRD6 | -0.250424886 | 0.018570615 | 0.34397431 | 1.731173714 |
| 27107 | ZBTB11 | -0.253679714 | 0.01879016 | 0.344994774 | 1.726069522 |
| 146059 | CDAN1 | -0.256668551 | 0.110784398 | 0.498711659 | 0.955521398 |
| 439921 | MXRA7 | -0.257679872 | 0.034189524 | 0.439589858 | 1.466106946 |
| 79663 | HSPBAP1 | -0.258049708 | 0.075237202 | 0.498711659 | 1.123567364 |
| 54956 | PARP16 | -0.258552489 | 0.051040162 | 0.493681625 | 1.292087956 |
| 284439 | SLC25A42 | -0.259151197 | 0.046594355 | 0.493681625 | 1.331666696 |
| 104 | ADARB1 | -0.259745804 | 0.089352909 | 0.498711659 | 1.048891304 |
| 22936 | ELL2 | -0.260194308 | 0.011601775 | 0.283202405 | 1.935475561 |
| 79621 | RNASEH2B | -0.260406113 | 0.044970449 | 0.493129872 | 1.347072776 |
| 79596 | OBI1 | -0.260483809 | 0.01020344 | 0.267175344 | 1.991253385 |
| 91283 | MSANTD3 | -0.260551077 | 0.011386871 | 0.28048456 | 1.943595599 |
| 389203 | SMIM20 | -0.260741344 | 0.025100197 | 0.385008994 | 1.60032287 |
| 131405 | TRIM71 | -0.262667704 | 0.059013427 | 0.498711659 | 1.229049164 |
| 9889 | ZBED4 | -0.262831493 | 0.014996778 | 0.31380509 | 1.824002037 |
| 57534 | MIB1 | -0.262931179 | 0.014088944 | 0.306540571 | 1.851121557 |
| 57001 | SDHAF3 | -0.267289104 | 0.010964423 | 0.275761554 | 1.960014218 |
| 4863 | NPAT | -0.267709967 | 0.029669581 | 0.416159794 | 1.527688587 |
| 10390 | CEPT1 | -0.268014546 | 0.018092916 | 0.339898128 | 1.742491433 |
| 80349 | WDR61 | -0.269057714 | 0.012625733 | 0.292441787 | 1.898743399 |
| 124411 | ZNF720 | -0.273022448 | 0.033230886 | 0.435838808 | 1.478458079 |
| 11196 | SEC23IP | -0.274908088 | 0.027018459 | 0.400411871 | 1.568339425 |
| 55132 | LARP1B | -0.276190718 | 0.016727574 | 0.331256573 | 1.77656704 |
| 8809 | IL18R1 | -0.277312338 | 0.200010602 | 0.498711659 | 0.698946983 |
| 56605 | ERO1B | -0.277358986 | 0.09445376 | 0.498711659 | 1.024780749 |
| 54545 | MTMR12 | -0.277819762 | 0.033175092 | 0.435710797 | 1.479187864 |
| 6118 | RPA2 | -0.277859263 | 0.167238315 | 0.498711659 | 0.776664217 |
| 54468 | MIOS | -0.278296432 | 0.017144585 | 0.332932369 | 1.765873023 |
| 55781 | RIOK2 | -0.278705274 | 0.01405079 | 0.306540571 | 1.852299257 |
| 6478 | SIAH2 | -0.279795974 | 0.018084056 | 0.339898128 | 1.742704157 |
| 4983 | OPHN1 | -0.280773607 | 0.065126465 | 0.498711659 | 1.186242494 |
| 6498 | SKIL | -0.282310498 | 0.015880911 | 0.323454888 | 1.799124588 |
| 64398 | MPP5 | -0.282584743 | 0.016769254 | 0.331256573 | 1.775486257 |
| 23098 | SARM1 | -0.28901685 | 0.141097591 | 0.498711659 | 0.850480401 |
| 23526 | ARHGAP45 | -0.29251893 | 0.036263083 | 0.449586194 | 1.440535276 |
| 26278 | SACS | -0.294976759 | 0.166588488 | 0.498711659 | 0.778355014 |
| 2972 | BRF1 | -0.29571464 | 0.162772414 | 0.498711659 | 0.788419196 |
| 55296 | TBC1D19 | -0.298440922 | 0.37267014 | 0.650387461 | 0.428675403 |
| 92345 | NAF1 | -0.299631101 | 0.026795309 | 0.398565301 | 1.57194123 |
| 85865 | GTPBP10 | -0.300146659 | 0.031721726 | 0.429536679 | 1.49864319 |
| 642 | BLMH | -0.302829728 | 0.020828696 | 0.359548613 | 1.681337919 |
| 8338 | H2AC20 | -0.303393948 | 0.363601897 | 0.645642915 | 0.43937386 |
| 79066 | METTL16 | -0.306019383 | 0.040711758 | 0.473084522 | 1.390280144 |
| 22800 | RRAS2 | -0.306188541 | 0.01337103 | 0.299188661 | 1.873835137 |
| 57801 | HES4 | -0.306728862 | 0.013944477 | 0.305284057 | 1.85559777 |
| 51095 | TRNT1 | -0.30706722 | 0.010666935 | 0.271648091 | 1.971960351 |
| 168451 | THAP5 | -0.308607021 | 0.013294066 | 0.298879665 | 1.876342169 |
| 29842 | TFCP2L1 | -0.309242751 | 0.051060196 | 0.493681625 | 1.291917522 |
| 1827 | RCAN1 | -0.311213693 | 0.081103482 | 0.498711659 | 1.0909605 |
| 6949 | TCOF1 | -0.312237641 | 0.027836877 | 0.404568902 | 1.55537949 |
| 200424 | TET3 | -0.316209234 | 0.075663576 | 0.498711659 | 1.121113137 |
| 84928 | TMEM209 | -0.316606474 | 0.022165588 | 0.368928404 | 1.654320743 |
| 6690 | SPINK1 | -0.316778313 | 0.048386287 | 0.493681625 | 1.315277703 |
| 84986 | ARHGAP19 | -0.317687963 | 0.0387038 | 0.465675598 | 1.412246393 |
| 84233 | TMEM126A | -0.317805592 | 0.016149397 | 0.326844006 | 1.791843689 |
| 9063 | PIAS2 | -0.320948538 | 0.121013977 | 0.498711659 | 0.917164466 |
| 55632 | G2E3 | -0.324370061 | 0.020528482 | 0.35878954 | 1.687643164 |
| 162073 | ITPRIPL2 | -0.324760877 | 0.01711842 | 0.332701942 | 1.766536322 |
| 84668 | FAM126A | -0.325046334 | 0.013721695 | 0.302271167 | 1.862592238 |
| 29886 | SNX8 | -0.32560791 | 0.06277753 | 0.498711659 | 1.202195776 |
| 81027 | TUBB1 | -0.327161466 | 0.149247673 | 0.498711659 | 0.826092431 |
| 9467 | SH3BP5 | -0.328572655 | 0.049382419 | 0.493681625 | 1.30642764 |
| 100507066 | CLIP1-AS1 | -0.329585496 | 0.277287573 | 0.549234809 | 0.557069593 |
| 55299 | BRIX1 | -0.332784583 | 0.030127422 | 0.41791598 | 1.521038029 |
| 23239 | PHLPP1 | -0.333675457 | 0.020563919 | 0.35878954 | 1.686894115 |
| 7342 | UBP1 | -0.334160339 | 0.024535154 | 0.381055674 | 1.610211212 |
| 55254 | TMEM39A | -0.33639845 | 0.011748237 | 0.284634119 | 1.930027301 |
| 22934 | RPIA | -0.337133007 | 0.043068528 | 0.483586376 | 1.365839971 |
| 115825 | WDFY2 | -0.337634577 | 0.021966585 | 0.367338732 | 1.658237455 |
| 388524 | RPSAP58 | -0.337890549 | 0.057785848 | 0.498711659 | 1.238178509 |
| 441951 | ZFAS1 | -0.339016636 | 0.017303146 | 0.334131729 | 1.761874928 |
| 6996 | TDG | -0.340466594 | 0.01291305 | 0.296176124 | 1.888971167 |
| 8487 | GEMIN2 | -0.342992757 | 0.077914942 | 0.498711659 | 1.108379248 |
| 85463 | ZC3H12C | -0.344672978 | 0.083135067 | 0.498711659 | 1.080215749 |
| 22982 | DIP2C | -0.34658449 | 0.054721834 | 0.497543972 | 1.261839356 |
| 25966 | C2CD2 | -0.34726307 | 0.013106799 | 0.297030712 | 1.882503361 |
| 11228 | RASSF8 | -0.349115386 | 0.040217775 | 0.470727014 | 1.39558196 |
| 65095 | KRI1 | -0.349683036 | 0.119127529 | 0.498711659 | 0.923987866 |
| 58512 | DLGAP3 | -0.349782829 | 0.114625689 | 0.498711659 | 0.940718041 |
| 1629 | DBT | -0.353838112 | 0.054055821 | 0.494974215 | 1.267157532 |
| 84678 | KDM2B | -0.355755115 | 0.012019499 | 0.287526255 | 1.920113634 |
| 10813 | UTP14A | -0.357178026 | 0.020080955 | 0.356542282 | 1.697215637 |
| 25852 | ARMC8 | -0.359812993 | 0.024717029 | 0.38247413 | 1.607003733 |
| 4286 | MITF | -0.366491013 | 0.033899674 | 0.439042933 | 1.469804478 |
| 55164 | SHQ1 | -0.366672104 | 0.044760866 | 0.492778626 | 1.34910152 |
| 55315 | SLC29A3 | -0.372154596 | 0.050937623 | 0.493681625 | 1.292961325 |
| 102724159 | LOC102724159 | -0.373364842 | 0.032752055 | 0.435178498 | 1.484761445 |
| 79649 | MAP7D3 | -0.375099658 | 0.212991161 | 0.498711659 | 0.671638419 |
| 54954 | FAM120C | -0.375511719 | 0.016256121 | 0.32760294 | 1.788983077 |
| 11281 | POU6F2 | -0.37740765 | 0.048088006 | 0.493681625 | 1.317963231 |
| 8360 | H4C4 | -0.383624072 | 0.216730305 | 0.498711659 | 0.664080358 |
| 3552 | IL1A | -0.385135172 | 0.013185031 | 0.297462033 | 1.879918845 |
| 54552 | GNL3L | -0.387271417 | 0.024280065 | 0.379643412 | 1.614750155 |
| 8808 | IL1RL2 | -0.390119185 | 0.234662332 | 0.499323556 | 0.629556618 |
| 205 | AK4 | -0.395073131 | 0.046912859 | 0.493681625 | 1.328708099 |
| 220963 | SLC16A9 | -0.403331116 | 0.284654829 | 0.5577404 | 0.545681444 |
| 22876 | INPP5F | -0.406309117 | 0.059502911 | 0.498711659 | 1.225461787 |
| 6502 | SKP2 | -0.408704855 | 0.046818157 | 0.493681625 | 1.329585686 |
| 7704 | ZBTB16 | -0.4208849 | 0.048014913 | 0.493681625 | 1.318623854 |
| 84460 | ZMAT1 | -0.421060277 | 0.025367949 | 0.387270387 | 1.595714644 |
| 85389 | SNORD14C | -0.421501919 | 0.123286615 | 0.498711659 | 0.909084071 |
| 241 | ALOX5AP | -0.435397939 | 0.035131709 | 0.444228985 | 1.454300723 |
| 23705 | CADM1 | -0.438192692 | 0.021442044 | 0.362624675 | 1.668733817 |
| 159090 | FAM122B | -0.443207145 | 0.010659698 | 0.271648091 | 1.972255099 |
| 83742 | MARVELD1 | -0.447682502 | 0.027657075 | 0.403609671 | 1.558193753 |
| 83539 | CHST9 | -0.448276261 | 0.017529332 | 0.335951946 | 1.756234633 |
| 9886 | RHOBTB1 | -0.459567091 | 0.014368263 | 0.308670759 | 1.842595731 |
| 148641 | SLC35F3 | -0.460488084 | 0.094802678 | 0.498711659 | 1.023179394 |
| 1843 | DUSP1 | -0.461761272 | 0.023026327 | 0.374689142 | 1.637775332 |
| 26747 | NUFIP1 | -0.471630195 | 0.012853599 | 0.295622799 | 1.890975253 |
| 9783 | RIMS3 | -0.472817282 | 0.017314452 | 0.334131729 | 1.761591249 |
| 84532 | ACSS1 | -0.474914636 | 0.019289948 | 0.348703239 | 1.714668943 |
| 6906 | SERPINA7 | -0.476071318 | 0.06815994 | 0.498711659 | 1.166470801 |
| 58495 | OVOL2 | -0.477003873 | 0.073735963 | 0.498711659 | 1.132320643 |
| 8404 | SPARCL1 | -0.489151682 | 0.02019514 | 0.356832556 | 1.694753132 |
| 284486 | THEM5 | -0.498064288 | 0.010349366 | 0.268064651 | 1.985086254 |
| 2977 | GUCY1A2 | -0.507247286 | 0.014419548 | 0.308670759 | 1.841048353 |
| 9657 | IQCB1 | -0.511408804 | 0.014180696 | 0.306912278 | 1.848302453 |
| 90649 | ZNF486 | -0.513601993 | 0.048538786 | 0.493681625 | 1.31391109 |
| 9529 | BAG5 | -0.51406625 | 0.010759568 | 0.273303001 | 1.968205165 |
| 79977 | GRHL2 | -0.518643761 | 0.026306564 | 0.395015801 | 1.579935873 |
| 24147 | FJX1 | -0.525148621 | 0.030268732 | 0.418657591 | 1.519005772 |
| 22797 | TFEC | -0.527784204 | 0.341963421 | 0.622279629 | 0.466020347 |
| 23281 | MTUS2 | -0.54153629 | 0.013057369 | 0.297030712 | 1.884144323 |
| 100033420 | SNORD116-8 | -0.544853611 | 0.196293118 | 0.498711659 | 0.707094926 |
| 389658 | ALKAL1 | -0.548873191 | 0.31182119 | 0.589334578 | 0.506094375 |
| 283643 | TEDC1 | -0.549601423 | 0.114002848 | 0.498711659 | 0.943084299 |
| 28965 | SLC27A6 | -0.552115201 | 0.147806718 | 0.498711659 | 0.830305826 |
| 10602 | CDC42EP3 | -0.558369831 | 0.011211621 | 0.278528703 | 1.950331592 |
| 205860 | TRIML2 | -0.559070605 | 0.040430015 | 0.471819992 | 1.393296098 |
| 100129434 | LOC100129434 | -0.573758014 | 0.027921671 | 0.40468463 | 1.554058594 |
| 55240 | STEAP3 | -0.57396052 | 0.103724777 | 0.498711659 | 0.98411749 |
| 5190 | PEX6 | -0.580474248 | 0.025663673 | 0.389481557 | 1.590681187 |
| 202559 | KHDRBS2 | -0.581393642 | 0.050514784 | 0.493681625 | 1.2965815 |
| 1417 | CRYBB3 | -0.584807867 | 0.112587026 | 0.498711659 | 0.948511653 |
| 10178 | TENM1 | -0.586487087 | 0.110028207 | 0.498711659 | 0.958495964 |
| 114805 | GALNT13 | -0.591275696 | 0.035939495 | 0.447767795 | 1.44442803 |
| 59344 | ALOXE3 | -0.59534521 | 0.028270021 | 0.407062493 | 1.548673869 |
| 101930332 | NA (ENTREZID: 101930332) | -0.596898377 | 0.048321333 | 0.493681625 | 1.315861094 |
| 400221 | FLJ22447 | -0.599642117 | 0.140109212 | 0.498711659 | 0.853533309 |
| 9472 | AKAP6 | -0.600770729 | 0.041469312 | 0.477220859 | 1.38227317 |
| 4739 | NEDD9 | -0.607450307 | 0.010193318 | 0.267175344 | 1.991684427 |
| 33 | ACADL | -0.613846206 | 0.181725547 | 0.498711659 | 0.740584015 |
| 100750225 | PCAT1 | -0.618398349 | 0.017249421 | 0.33387683 | 1.763225478 |
| 220416 | LRRC63 | -0.620675772 | 0.101628071 | 0.498711659 | 0.992986318 |
| 5255 | PHKA1 | -0.621354858 | 0.023490914 | 0.376120459 | 1.629100085 |
| 79627 | OGFRL1 | -0.626669628 | 0.025865533 | 0.391707636 | 1.587278568 |
| 339751 | MAP3K20-AS1 | -0.638984794 | 0.055999435 | 0.498711659 | 1.251816355 |
| 79986 | ZNF702P | -0.648820702 | 0.02061712 | 0.358935079 | 1.685772001 |
| 11075 | STMN2 | -0.651351484 | 0.167841929 | 0.498711659 | 0.775099538 |
| 54622 | ARL15 | -0.675158575 | 0.017411386 | 0.334964225 | 1.759166656 |
| 100507500 | EGFR-AS1 | -0.679737522 | 0.028810566 | 0.410886996 | 1.54044821 |
| 100129550 | LINC02035 | -0.680726243 | 0.05286163 | 0.493681625 | 1.276859449 |
| 257169 | C9orf43 | -0.69655285 | 0.04192092 | 0.478959251 | 1.377569195 |
| 2917 | GRM7 | -0.697166227 | 0.162123761 | 0.498711659 | 0.79015333 |
| 23240 | TMEM131L | -0.722480714 | 0.018332686 | 0.342246301 | 1.7367739 |
| 2201 | FBN2 | -0.737102622 | 0.011687779 | 0.284136254 | 1.932268009 |
| 441172 | FLJ46906 | -0.737932165 | 0.04287999 | 0.482698455 | 1.367745325 |
| 2823 | GPM6A | -0.74590017 | 0.059534734 | 0.498711659 | 1.225229582 |
| 1813 | DRD2 | -0.750376895 | 0.088362542 | 0.498711659 | 1.053731799 |
| 7056 | THBD | -0.752887687 | 0.012389685 | 0.290268247 | 1.906939735 |
| 57528 | KCTD16 | -0.753490181 | 0.042740999 | 0.482057 | 1.369155331 |
| 1490 | CCN2 | -0.761563031 | 0.017659779 | 0.337222211 | 1.753014736 |
| 83478 | ARHGAP24 | -0.76940795 | 0.052893582 | 0.493681625 | 1.276597021 |
| 5067 | CNTN3 | -0.770878161 | 0.248134523 | 0.516075209 | 0.605312808 |
| 361 | AQP4 | -0.783557166 | 0.014929571 | 0.312859893 | 1.825952672 |
| 145837 | DRAIC | -0.80732525 | 0.082832377 | 0.498711659 | 1.081799876 |
| 22915 | MMRN1 | -0.830966478 | 0.026652185 | 0.39747311 | 1.574267181 |
| 23086 | EXPH5 | -0.838888757 | 0.060941987 | 0.498711659 | 1.21508339 |
| 92745 | SLC38A5 | -0.846255983 | 0.03339565 | 0.436387242 | 1.476310099 |
| 4741 | NEFM | -0.863341213 | 0.023667108 | 0.376908324 | 1.625854807 |
| 1734 | DIO2 | -0.865996689 | 0.015071332 | 0.314669456 | 1.821848363 |
| 151354 | LRATD1 | -0.877565807 | 0.132495028 | 0.498711659 | 0.877800419 |
| 79822 | ARHGAP28 | -0.877914262 | 0.040705817 | 0.473084522 | 1.390343524 |
| 81624 | DIAPH3 | -0.912833353 | 0.019368696 | 0.348968345 | 1.712899617 |
| 51314 | NME8 | -0.915492164 | 0.353874027 | 0.63584465 | 0.451151312 |
| 827 | CAPN6 | -0.932862463 | 0.047631258 | 0.493681625 | 1.322107948 |
| 91584 | PLXNA4 | -0.939498795 | 0.065824271 | 0.498711659 | 1.181613942 |
| 414224 | AGAP12P | -0.943339155 | 0.148383555 | 0.498711659 | 0.828614228 |
| 101928773 | LINC01449 | -1.030443453 | 0.035634903 | 0.445969218 | 1.448124419 |
| 2070 | EYA4 | -1.057172757 | 0.018983456 | 0.347092868 | 1.72162472 |
| 3354 | HTR1E | -1.071328309 | 0.097933149 | 0.498711659 | 1.009070281 |
| 100506963 | ELOA-AS1 | -1.120511527 | 0.036357257 | 0.4502243 | 1.43940889 |
| 941 | CD80 | -1.130155276 | 0.047429963 | 0.493681625 | 1.323947214 |
| 338645 | LUZP2 | -1.358062872 | 0.01718969 | 0.333174918 | 1.764731955 |
| 100507096 | GIMD1 | -1.429512111 | 0.018374431 | 0.342774403 | 1.735786101 |
| 101927378 | LOC101927378 | -1.598418457 | 0.017030296 | 0.331670276 | 1.768777804 |
| 100861510 | LRRC3-DT | -1.769689624 | 0.039438624 | 0.468756688 | 1.404078246 |
| 151531 | UPP2 | -1.822091994 | 0.016787399 | 0.331256573 | 1.775016587 |
| 408186 | LOC408186 | -1.856407474 | 0.022810231 | 0.373390802 | 1.641870317 |
| 9982 | FGFBP1 | 3.751493683 | 0.000534459 | 0.064547514 | 3.272085605 |
| 5967 | REG1A | 3.232760281 | 0.003574927 | 0.166409901 | 2.446732822 |
| 7409 | VAV1 | 3.160061417 | 0.002906185 | 0.153571991 | 2.536676743 |
| 191585 | PLAC4 | 2.930531771 | 0.001020799 | 0.087521032 | 2.991059764 |
| 3932 | LCK | 2.929888009 | 0.001005964 | 0.086558625 | 2.997417561 |
| 54857 | GDPD2 | 2.817981727 | 0.001212915 | 0.095887882 | 2.916169633 |
| 64600 | PLA2G2F | 2.757411635 | 0.001926648 | 0.121033552 | 2.715197624 |
| 1048 | CEACAM5 | 2.707422251 | 0.008143582 | 0.240645298 | 2.089184526 |
| 115362 | GBP5 | 2.625987062 | 0.001226115 | 0.096238624 | 2.911468794 |
| 101928372 | LOC101928372 | 2.624327942 | 0.003410124 | 0.161830729 | 2.467229829 |
| 10871 | CD300C | 2.609685531 | 0.004889695 | 0.191341678 | 2.31071823 |
| 91543 | RSAD2 | 2.583585063 | 0.003177729 | 0.157756498 | 2.497883143 |
| 4939 | OAS2 | 2.522368947 | 0.008040484 | 0.24007312 | 2.094717808 |
| 101929623 | LINC01215 | 2.517573831 | 0.004828701 | 0.190939426 | 2.316169686 |
| 27156 | RSPH14 | 2.505769832 | 0.000958168 | 0.083457638 | 3.018558337 |
| 101927489 | TCERG1L-AS1 | 2.435149276 | 0.000408067 | 0.057079117 | 3.389268525 |
| 1014 | CDH16 | 2.415484556 | 0.006374076 | 0.214753 | 2.195582763 |
| 387111 | LINC00222 | 2.333216636 | 0.003360721 | 0.161213936 | 2.47356754 |
| 100820829 | MYZAP | 2.327799404 | 0.000350276 | 0.053890511 | 3.455589618 |
| 6261 | RYR1 | 2.325362322 | 0.004834846 | 0.190939426 | 2.315617355 |
| 84873 | ADGRG7 | 2.276794371 | 0.000385101 | 0.056193108 | 3.414425354 |
| 100861550 | PLUT | 2.218697982 | 0.005598586 | 0.203549087 | 2.251921646 |
| 101926963 | PRKAR1B-AS1 | 2.213029198 | 0.000475492 | 0.061633976 | 3.322856786 |
| 92749 | DRC1 | 2.141643544 | 0.00038449 | 0.056193108 | 3.415114951 |
| 100616314 | MIR4750 | 2.071603258 | 0.002121713 | 0.128289957 | 2.673313363 |
| 63036 | CELA2A | 2.013755523 | 0.000924563 | 0.082183696 | 3.034063491 |
| 119385 | AGAP11 | 1.998690176 | 0.003015601 | 0.155404722 | 2.520626121 |
| 250 | ALPP | 1.968464764 | 0.005121901 | 0.194525898 | 2.29056882 |
| 1812 | DRD1 | 1.967956558 | 0.003518638 | 0.164870848 | 2.453625411 |
| 25878 | MXRA5 | 1.951792007 | 0.006358381 | 0.214753 | 2.196653452 |
| 102724094 | MUC12-AS1 | 1.857400539 | 0.000323236 | 0.051402596 | 3.490480276 |
| 5175 | PECAM1 | 1.839224616 | 0.009822864 | 0.262143076 | 2.007761869 |
| 25791 | NGEF | 1.808564356 | 0.000947843 | 0.082812312 | 3.023263593 |
| 643977 | FLJ32255 | 1.804030326 | 0.003957642 | 0.176310508 | 2.402563494 |
| 360 | AQP3 | 1.765279181 | 0.000682327 | 0.069238034 | 3.166007443 |
| 54854 | FAM83E | 1.755982345 | 0.00387968 | 0.175140732 | 2.411204094 |
| 25825 | BACE2 | 1.717011386 | 0.000594032 | 0.066407614 | 3.226190159 |
| 4486 | MST1R | 1.714708249 | 0.003292631 | 0.159661255 | 2.482456937 |
| 2781 | GNAZ | 1.695238266 | 0.005759041 | 0.206474704 | 2.23964983 |
| 56154 | TEX15 | 1.682460632 | 0.000619299 | 0.067521014 | 3.208099621 |
| 441307 | HRAT92 | 1.661260292 | 0.007019855 | 0.226908729 | 2.153671858 |
| 7134 | TNNC1 | 1.612738491 | 0.000603428 | 0.066670585 | 3.219374542 |
| 3914 | LAMB3 | 1.609879749 | 0.003978837 | 0.176529793 | 2.400243852 |
| 103021165 | CASC19 | 1.605854977 | 0.005453982 | 0.200863582 | 2.2632863 |
| 9313 | MMP20 | 1.604740642 | 0.001982061 | 0.12217869 | 2.702882984 |
| 768220 | MIR765 | 1.568932272 | 0.002705584 | 0.148438915 | 2.567738978 |
| 6262 | RYR2 | 1.56643179 | 0.000394354 | 0.056320771 | 3.40411375 |
| 5414 | SEPTIN4 | 1.506935603 | 0.004022282 | 0.177087614 | 2.395527485 |
| 8938 | BAIAP3 | 1.489879325 | 0.000433605 | 0.059373016 | 3.362905718 |
| 440836 | ODF3B | 1.475835419 | 0.002355305 | 0.135932667 | 2.627952846 |
| 79170 | PRR15L | 1.464514982 | 0.001282411 | 0.097363766 | 2.891972765 |
| 27121 | DKK4 | 1.454847764 | 0.003539854 | 0.1655917 | 2.45101465 |
| 113763 | ZBED6CL | 1.442540682 | 0.003357819 | 0.161213936 | 2.473942718 |
| 8792 | TNFRSF11A | 1.441641481 | 0.002757974 | 0.150278934 | 2.559409832 |
| 120406 | NXPE2 | 1.441516964 | 0.00774473 | 0.234999627 | 2.110993718 |
| 130367 | SGPP2 | 1.431871363 | 0.00556527 | 0.203003874 | 2.254513761 |
| 100874216 | ZNF385D-AS1 | 1.41548333 | 0.003886571 | 0.175173294 | 2.410433394 |
| 2564 | GABRE | 1.412023618 | 0.000361345 | 0.054591567 | 3.44207795 |
| 201799 | TMEM154 | 1.406744908 | 0.000674929 | 0.069238034 | 3.170741911 |
| 126353 | MISP | 1.391207571 | 0.00226389 | 0.132816438 | 2.645144679 |
| 136 | ADORA2B | 1.387896933 | 0.008644084 | 0.249070459 | 2.063281021 |
| 2877 | GPX2 | 1.367607893 | 0.006863616 | 0.222898562 | 2.163447022 |
| 375593 | TRIM73 | 1.364967925 | 0.000656727 | 0.069238034 | 3.182615128 |
| 285513 | GPRIN3 | 1.330853249 | 0.000542057 | 0.064791918 | 3.265955043 |
| 407 | ARR3 | 1.310429574 | 0.007333876 | 0.229547419 | 2.134666437 |
| 93010 | B3GNT7 | 1.283028469 | 0.000668889 | 0.069238034 | 3.174645946 |
| 341208 | HEPHL1 | 1.280678196 | 0.007883629 | 0.237511356 | 2.103273822 |
| 56106 | PCDHGA10 | 1.258367594 | 0.000391049 | 0.056320771 | 3.40776882 |
| 283422 | LINC01559 | 1.245661261 | 0.008118202 | 0.240645298 | 2.090540147 |
| 7049 | TGFBR3 | 1.232090449 | 0.003244267 | 0.158590987 | 2.488883411 |
| 10 | NAT2 | 1.230656644 | 0.003244986 | 0.158590987 | 2.488787173 |
| 101926941 | SPRY4-AS1 | 1.214593685 | 0.001618466 | 0.111261127 | 2.79089642 |
| 1901 | S1PR1 | 1.214359446 | 0.00308499 | 0.1572723 | 2.510746239 |
| 6275 | S100A4 | 1.201175307 | 0.005220135 | 0.196846937 | 2.282318265 |
| 9068 | ANGPTL1 | 1.188332355 | 0.000819383 | 0.076786728 | 3.086513051 |
| 51450 | PRRX2 | 1.173463206 | 0.00203517 | 0.124009999 | 2.691399308 |
| 9220 | TIAF1 | 1.16386453 | 0.001319895 | 0.098111059 | 2.879460616 |
| 10008 | KCNE3 | 1.146002123 | 0.004436178 | 0.185523245 | 2.352991036 |
| 978 | CDA | 1.139540626 | 0.001971941 | 0.121989709 | 2.705106083 |
| 6004 | RGS16 | 1.136942323 | 0.002873125 | 0.153062642 | 2.541645479 |
| 81563 | C1orf21 | 1.135959916 | 0.000729435 | 0.071917685 | 3.137013402 |
| 144455 | E2F7 | 1.122922108 | 0.001178697 | 0.094313502 | 2.928597822 |
| 440556 | PRDM16-DT | 1.11514553 | 0.000575083 | 0.0661113 | 3.24026947 |
| 94015 | TTYH2 | 1.108503878 | 0.00313666 | 0.15735649 | 2.503532554 |
| 6843 | VAMP1 | 1.104750944 | 0.000680075 | 0.069238034 | 3.16744319 |
| 51478 | HSD17B7 | 1.104257283 | 0.002646904 | 0.146223404 | 2.57726181 |
| 6536 | SLC6A9 | 1.099344924 | 0.003293538 | 0.159661255 | 2.482337321 |
| 26240 | FAM50B | 1.093685456 | 0.001995534 | 0.12217869 | 2.699940868 |
| 2487 | FRZB | 1.092369499 | 0.006127285 | 0.210482178 | 2.212731919 |
| 102725191 | LOC102725191 | 1.088483116 | 0.001723211 | 0.115675097 | 2.763661542 |
| 105372417 | NA (ENTREZID: 105372417) | 1.072524413 | 0.001082155 | 0.090047638 | 2.96571053 |
| 8646 | CHRD | 1.06900572 | 0.000687625 | 0.069238034 | 3.162648342 |
| 2591 | GALNT3 | 1.059660866 | 0.000509425 | 0.062971482 | 3.292919746 |
| 2296 | FOXC1 | 1.05275202 | 0.009797803 | 0.261798618 | 2.008871297 |
| 27111 | SDCBP2 | 1.039579852 | 0.002790265 | 0.151162781 | 2.554354548 |
| 285512 | FAM13A-AS1 | 1.035218427 | 0.002957466 | 0.154939588 | 2.529080239 |
| 375719 | AQP7P1 | 1.034514785 | 0.005673526 | 0.20470111 | 2.246146951 |
| 101926962 | ADGRF5-AS1 | 1.029793823 | 0.006858262 | 0.222898562 | 2.163785928 |
| 3638 | INSIG1 | 1.029008444 | 0.003967685 | 0.176310508 | 2.401462814 |
| 375287 | RBM43 | 1.028037358 | 0.000863429 | 0.078832974 | 3.063773369 |
| 23474 | ETHE1 | 1.027787389 | 0.001429687 | 0.103035443 | 2.844759032 |
| 9466 | IL27RA | 1.027530857 | 0.001249632 | 0.096684719 | 2.903217862 |
| 11187 | PKP3 | 1.018844019 | 0.000471674 | 0.061633976 | 3.326358063 |
| 4598 | MVK | 1.0170802 | 0.006093705 | 0.210482178 | 2.215118574 |
| 3433 | IFIT2 | 1.014234352 | 0.002612424 | 0.144599964 | 2.582956335 |
| 26577 | PCOLCE2 | 1.008428557 | 0.006523163 | 0.216525907 | 2.185541769 |
| 25894 | PLEKHG4 | 0.991088331 | 0.001074455 | 0.090047638 | 2.968811769 |
| 11274 | USP18 | 0.974283541 | 0.001731768 | 0.115693396 | 2.76151029 |
| 282679 | AQP11 | 0.942980268 | 0.004274585 | 0.182796435 | 2.369106043 |
| 154 | ADRB2 | 0.940955797 | 0.001026396 | 0.087521032 | 2.988685049 |
| 1848 | DUSP6 | 0.933051668 | 0.00111999 | 0.091913627 | 2.950785855 |
| 101669762 | BLACAT1 | 0.912232848 | 0.004349844 | 0.1835428 | 2.361526318 |
| 84226 | C2orf16 | 0.907463156 | 0.001913857 | 0.120588544 | 2.718090515 |
| 84868 | HAVCR2 | 0.903614581 | 0.006109853 | 0.210482178 | 2.213969239 |
| 55809 | TRERF1 | 0.902088469 | 0.001438562 | 0.103402233 | 2.842071416 |
| 91107 | TRIM47 | 0.897523126 | 0.001951928 | 0.121280066 | 2.709536206 |
| 6347 | CCL2 | 0.893862925 | 0.009631541 | 0.259687407 | 2.016304222 |
| 148932 | MOB3C | 0.886304745 | 0.001310557 | 0.098111059 | 2.882544086 |
| 26873 | OPLAH | 0.885009145 | 0.00283489 | 0.152107164 | 2.547463788 |
| 1909 | EDNRA | 0.875456226 | 0.001662618 | 0.112673108 | 2.779207522 |
| 8876 | VNN1 | 0.874570457 | 0.000620637 | 0.067521014 | 3.207162337 |
| 29800 | ZDHHC1 | 0.873311803 | 0.007696415 | 0.234612665 | 2.113711523 |
| 9627 | SNCAIP | 0.870028401 | 0.002223992 | 0.132113512 | 2.652866779 |
| 57026 | PDXP | 0.852556885 | 0.001278157 | 0.097301017 | 2.893415797 |
| 100505832 | PROX1-AS1 | 0.849271668 | 0.000536475 | 0.064547514 | 3.270450512 |
| 100422737 | LINC02532 | 0.848808829 | 0.000531432 | 0.064487244 | 3.274552298 |
| 1717 | DHCR7 | 0.839654137 | 0.004876783 | 0.191265528 | 2.311866569 |
| 10039 | PARP3 | 0.836251256 | 0.000655341 | 0.069238034 | 3.18353266 |
| 285386 | TPRG1 | 0.830489348 | 0.002594611 | 0.143894516 | 2.585927745 |
| 9572 | NR1D1 | 0.824377359 | 0.00563251 | 0.203739022 | 2.249298029 |
| 4599 | MX1 | 0.820445383 | 0.001933124 | 0.121070141 | 2.713740287 |
| 8535 | CBX4 | 0.814122261 | 0.000513909 | 0.062971482 | 3.289113777 |
| 197021 | LCTL | 0.80868767 | 0.004904635 | 0.191564107 | 2.309393307 |
| 1181 | CLCN2 | 0.800768021 | 0.003177909 | 0.157756498 | 2.497858543 |
| 5329 | PLAUR | 0.784628441 | 0.005176609 | 0.196069811 | 2.285954637 |
| 401232 | LINC01011 | 0.780023638 | 0.004610063 | 0.187673063 | 2.33629314 |
| 80852 | GRIP2 | 0.77015925 | 0.008810812 | 0.250935818 | 2.054984065 |
| 84791 | LINC00467 | 0.769522604 | 0.001057387 | 0.089093483 | 2.975766033 |
| 7076 | TIMP1 | 0.767035156 | 0.003957194 | 0.176310508 | 2.402612658 |
| 10509 | SEMA4B | 0.747157511 | 0.000495027 | 0.062196021 | 3.305371113 |
| 29785 | CYP2S1 | 0.747079549 | 0.000502921 | 0.062909466 | 3.29850023 |
| 378884 | NHLRC1 | 0.743667619 | 0.003811107 | 0.173702055 | 2.418948858 |
| 57552 | NCEH1 | 0.737723754 | 0.008662962 | 0.249070459 | 2.062333591 |
| 56100 | PCDHGB6 | 0.723040461 | 0.00060948 | 0.0668193 | 3.215040541 |
| 7172 | TPMT | 0.712906183 | 0.008225779 | 0.242134364 | 2.084822963 |
| 5064 | PALM | 0.712125961 | 0.000924955 | 0.082183696 | 3.033879396 |
| 104797538 | KCNMB2-AS1 | 0.711053926 | 0.005805188 | 0.207604937 | 2.236183711 |
| 30845 | EHD3 | 0.705825932 | 0.007939178 | 0.238049599 | 2.100224461 |
| 57449 | PLEKHG5 | 0.690098493 | 0.001050563 | 0.088781953 | 2.978577899 |
| 129642 | MBOAT2 | 0.675805895 | 0.00150468 | 0.105755919 | 2.822555852 |
| 100132707 | PAXIP1-AS2 | 0.675371148 | 0.000516723 | 0.062971482 | 3.286742207 |
| 6508 | SLC4A3 | 0.675126104 | 0.0006712 | 0.069238034 | 3.173148052 |
| 9722 | NOS1AP | 0.648602035 | 0.004166607 | 0.18007732 | 2.380217461 |
| 649446 | DLGAP1-AS1 | 0.619627684 | 0.001179126 | 0.094313502 | 2.928439784 |
| 55083 | KIF26B | 0.618310807 | 0.001634243 | 0.111261127 | 2.786683367 |
| 91860 | CALML4 | 0.617921899 | 0.001084567 | 0.090047638 | 2.964743614 |
| 7263 | TST | 0.61326239 | 0.001762933 | 0.116474313 | 2.753764193 |
| 55848 | PLGRKT | 0.604704032 | 0.008108899 | 0.240645298 | 2.091038109 |
| 645676 | ASH1L-AS1 | 0.599092446 | 0.009131643 | 0.254356697 | 2.039451076 |
| 55195 | CCDC198 | 0.59343181 | 0.001257263 | 0.096747942 | 2.900573865 |
| 112770 | GLMP | 0.577591576 | 0.000765775 | 0.073973456 | 3.115898816 |
| 1308 | COL17A1 | 0.575290183 | 0.000853436 | 0.078600248 | 3.068829041 |
| 2920 | CXCL2 | 0.560238852 | 0.005692584 | 0.205128089 | 2.244690552 |
| 5947 | RBP1 | 0.558203011 | 0.000860508 | 0.078819755 | 3.065245088 |
| 149473 | CCDC24 | 0.557041981 | 0.008106659 | 0.240645298 | 2.091158095 |
| 10591 | DNPH1 | 0.554300281 | 0.00083601 | 0.077831198 | 3.077788528 |
| 728769 | SCAMP1-AS1 | 0.553457568 | 0.001269491 | 0.096901046 | 2.896370374 |
| 101 | ADAM8 | 0.542440315 | 0.006031327 | 0.210482178 | 2.219587125 |
| 126375 | ZNF792 | 0.52951388 | 0.003031773 | 0.155955074 | 2.518303319 |
| 399687 | MYO18A | 0.517646348 | 0.00116606 | 0.094313502 | 2.933279102 |
| 4000 | LMNA | 0.517198799 | 0.000368459 | 0.055065291 | 3.433610831 |
| 56521 | DNAJC12 | 0.500501092 | 0.008395314 | 0.244748399 | 2.075963056 |
| 2950 | GSTP1 | 0.497148929 | 0.000494683 | 0.062196021 | 3.305673014 |
| 401261 | LOC401261 | 0.4873815 | 0.00887908 | 0.251546418 | 2.051632031 |
| 85462 | FHDC1 | 0.463835098 | 0.005450947 | 0.200863582 | 2.263528041 |
| 135932 | TMEM139 | 0.463748754 | 0.003963702 | 0.176310508 | 2.401899004 |
| 26167 | PCDHB5 | 0.462960091 | 0.003361108 | 0.161213936 | 2.473517532 |
| 64761 | PARP12 | 0.457055997 | 0.004347634 | 0.1835428 | 2.361747024 |
| 29948 | OSGIN1 | 0.453560473 | 0.009559221 | 0.25887555 | 2.019577498 |
| 57088 | PLSCR4 | 0.4498896 | 0.004097673 | 0.178819783 | 2.387462702 |
| 51310 | SLC22A17 | 0.442194858 | 0.005386393 | 0.199813058 | 2.268701963 |
| 55362 | TMEM63B | 0.436161554 | 0.000400176 | 0.056532267 | 3.397748961 |
| 5324 | PLAG1 | 0.430917195 | 0.009737448 | 0.261090495 | 2.011554849 |
| 7846 | TUBA1A | 0.430426428 | 0.006535049 | 0.216525907 | 2.184751152 |
| 4241 | MELTF | 0.425032915 | 0.000760019 | 0.07390668 | 3.11917555 |
| 54894 | RNF43 | 0.41470595 | 0.007738616 | 0.234999627 | 2.111336703 |
| 9926 | LPGAT1 | 0.411300338 | 0.004580334 | 0.187134642 | 2.339102852 |
| 5274 | SERPINI1 | 0.406881723 | 0.000420901 | 0.058398024 | 3.375820042 |
| 25959 | KANK2 | 0.402659582 | 0.006455224 | 0.215898284 | 2.190088683 |
| 10318 | TNIP1 | 0.392148721 | 0.003062239 | 0.156953567 | 2.513960917 |
| 4052 | LTBP1 | 0.377891491 | 0.003413862 | 0.161830729 | 2.466754039 |
| 5315 | PKM | 0.371115331 | 0.009906433 | 0.262890817 | 2.004082693 |
| 8986 | RPS6KA4 | 0.368664833 | 0.009057023 | 0.254124676 | 2.043014529 |
| 57175 | CORO1B | 0.367616046 | 0.006615509 | 0.218596445 | 2.179436735 |
| 4550 | RNR2 | 0.361185868 | 0.008762333 | 0.250741009 | 2.057380246 |
| 4967 | OGDH | 0.357859379 | 0.004642429 | 0.187812491 | 2.33325473 |
| 8509 | NDST2 | 0.356337744 | 0.002250691 | 0.132413438 | 2.647684126 |
| 10161 | LPAR6 | 0.350881443 | 0.00588081 | 0.207828655 | 2.230562852 |
| 57147 | SCYL3 | 0.349764864 | 0.003323589 | 0.160498843 | 2.478392687 |
| 93611 | FBXO44 | 0.347354327 | 0.004130145 | 0.179324377 | 2.384034701 |
| 100381270 | ZBED6 | 0.342472155 | 0.000749046 | 0.073341937 | 3.125491511 |
| 284129 | SLC26A11 | 0.33906643 | 0.002516373 | 0.140378039 | 2.599224983 |
| 50488 | MINK1 | 0.329129821 | 0.004874521 | 0.191265528 | 2.312068054 |
| 64332 | NFKBIZ | 0.307805334 | 0.005097659 | 0.194292647 | 2.292629219 |
| 2987 | GUK1 | 0.304700236 | 0.009495996 | 0.258330594 | 2.022459477 |
| 226 | ALDOA | 0.29850717 | 0.004746696 | 0.18977162 | 2.323608582 |
| 9520 | NPEPPS | 0.289930191 | 0.006120133 | 0.210482178 | 2.21323914 |
| 8567 | MADD | 0.272218475 | 0.001597964 | 0.110399468 | 2.796433009 |
| 151011 | SEPTIN10 | 0.269616413 | 0.000475656 | 0.061633976 | 3.322707021 |
| 7533 | YWHAH | 0.26516489 | 0.00892688 | 0.251716736 | 2.049300303 |
| 2260 | FGFR1 | 0.257164249 | 0.006721011 | 0.220118925 | 2.172565394 |
| 8079 | MLF2 | 0.251424018 | 0.000652934 | 0.069238034 | 3.185130716 |
| 387921 | NHLRC3 | 0.236018605 | 0.00450681 | 0.185633035 | 2.346130751 |
| 4779 | NFE2L1 | 0.233196846 | 0.004631945 | 0.187812491 | 2.334236606 |
| 83637 | ZMIZ2 | 0.227376999 | 0.0088059 | 0.250935818 | 2.055226251 |
| 5364 | PLXNB1 | 0.223017058 | 0.006057708 | 0.210482178 | 2.217691665 |
| 6272 | SORT1 | 0.221023263 | 0.003684997 | 0.169862817 | 2.433562861 |
| 2108 | ETFA | 0.217707159 | 0.000321252 | 0.051402596 | 3.49315416 |
| 1718 | DHCR24 | 0.204477304 | 0.008101266 | 0.240645298 | 2.091447108 |
| 9414 | TJP2 | 0.201609064 | 0.001883591 | 0.11965225 | 2.725013393 |
| 9887 | SMG7 | 0.192031602 | 0.002834679 | 0.152107164 | 2.547496114 |
| 9877 | ZC3H11A | 0.189921853 | 0.002478033 | 0.139312281 | 2.605892914 |
| 9746 | CLSTN3 | 0.177492108 | 0.008704182 | 0.249651752 | 2.060272037 |
| 375 | ARF1 | 0.176118196 | 0.008864181 | 0.251446959 | 2.052361385 |
| 6746 | SSR2 | 0.17033054 | 0.004235607 | 0.181740357 | 2.373084343 |
| 4316 | MMP7 | 3.222852586 | 0.023288112 | 0.375753064 | 1.632865719 |
| 284021 | MILR1 | 2.524621563 | 0.016249383 | 0.32760294 | 1.789163125 |
| 56147 | PCDHA1 | 2.371359358 | 0.036329583 | 0.450077884 | 1.439739587 |
| 90226 | UCN2 | 2.251383254 | 0.041225887 | 0.47547078 | 1.384829991 |
| 1510 | CTSE | 2.223952615 | 0.013911641 | 0.30503555 | 1.856621638 |
| 85477 | SCIN | 2.155316073 | 0.010906014 | 0.274535696 | 1.962333949 |
| 100131551 | LINC00887 | 2.084835518 | 0.014911544 | 0.312712928 | 1.826477386 |
| 100505817 | LINC02582 | 2.046991413 | 0.024558263 | 0.381055674 | 1.609802354 |
| 154064 | RAET1L | 2.022019515 | 0.025857503 | 0.391707636 | 1.587413416 |
| 5794 | PTPRH | 1.890450157 | 0.011797315 | 0.285134068 | 1.928216824 |
| 10103 | TSPAN1 | 1.847881097 | 0.021427084 | 0.362587642 | 1.669036928 |
| 10562 | OLFM4 | 1.847791777 | 0.076032953 | 0.498711659 | 1.118998142 |
| 101929505 | LINC02150 | 1.783263591 | 0.019346426 | 0.348968345 | 1.713399253 |
| 5268 | SERPINB5 | 1.697957463 | 0.103376153 | 0.498711659 | 0.985579634 |
| 256536 | TCERG1L | 1.664692252 | 0.024488827 | 0.380810646 | 1.611032017 |
| 390874 | ONECUT3 | 1.653730427 | 0.026782544 | 0.398565301 | 1.572148173 |
| 154860 | FEZF1-AS1 | 1.624368393 | 0.010508904 | 0.269762522 | 1.978442575 |
| 246777 | SPESP1 | 1.612054962 | 0.010275358 | 0.267187549 | 1.988203038 |
| 23166 | STAB1 | 1.610500095 | 0.02019825 | 0.356832556 | 1.694686257 |
| 9048 | ARTN | 1.601804715 | 0.067932894 | 0.498711659 | 1.167919884 |
| 644165 | BCRP3 | 1.583283218 | 0.03678556 | 0.453532125 | 1.434322628 |
| 64919 | BCL11B | 1.526954954 | 0.019478116 | 0.350051327 | 1.710453052 |
| 7425 | VGF | 1.521105817 | 0.041032938 | 0.474959858 | 1.386867386 |
| 105376068 | NA (ENTREZID: 105376068) | 1.518563944 | 0.043651137 | 0.486449782 | 1.36000444 |
| 875 | CBS | 1.501413843 | 0.039125636 | 0.468074463 | 1.40753859 |
| 574432 | IBA57-DT | 1.433319665 | 0.01581673 | 0.323365671 | 1.800883299 |
| 54997 | TESC | 1.414640821 | 0.011798997 | 0.285134068 | 1.928154909 |
| 101929768 | OSMR-AS1 | 1.405535341 | 0.017321426 | 0.334131729 | 1.761416357 |
| 100287036 | LOC100287036 | 1.402989842 | 0.022590356 | 0.372096128 | 1.646076925 |
| 56000 | NXF3 | 1.402350901 | 0.051384963 | 0.493681625 | 1.289163952 |
| 4907 | NT5E | 1.367467761 | 0.014910632 | 0.312712928 | 1.826503948 |
| 101926933 | LOC101926933 | 1.357342419 | 0.039961218 | 0.470279473 | 1.398361283 |
| 104326052 | NRIR | 1.344038078 | 0.031864211 | 0.430644586 | 1.496696831 |
| 105372412 | LOC105372412 | 1.333861358 | 0.076990052 | 0.498711659 | 1.113565387 |
| 220980 | TMEM72-AS1 | 1.33057133 | 0.017101905 | 0.332608623 | 1.76695551 |
| 56103 | PCDHGB2 | 1.326295901 | 0.012369211 | 0.290268247 | 1.907658002 |
| 1381 | CRABP1 | 1.306583744 | 0.029878339 | 0.417048939 | 1.52464355 |
| 4065 | LY75 | 1.261174682 | 0.014724638 | 0.311899748 | 1.831955373 |
| 222256 | CDHR3 | 1.235350451 | 0.030140432 | 0.41791598 | 1.520850527 |
| 56105 | PCDHGA11 | 1.220140826 | 0.019284597 | 0.348703239 | 1.714789432 |
| 3910 | LAMA4 | 1.217558099 | 0.05347773 | 0.493681625 | 1.271827036 |
| 10071 | MUC12 | 1.154696396 | 0.024911857 | 0.383668644 | 1.603593898 |
| 8538 | BARX2 | 1.151571624 | 0.029391126 | 0.414029782 | 1.531783775 |
| 440900 | LINC01191 | 1.148471762 | 0.062781319 | 0.498711659 | 1.202169564 |
| 79173 | BRME1 | 1.133909986 | 0.013089155 | 0.297030712 | 1.883088389 |
| 124961 | ZFP3 | 1.106231067 | 0.019058301 | 0.347348646 | 1.719915818 |
| 4060 | LUM | 1.102383549 | 0.016991697 | 0.331600848 | 1.769763245 |
| 51666 | ASB4 | 1.100478158 | 0.015927049 | 0.323728391 | 1.797864684 |
| 7099 | TLR4 | 1.084365152 | 0.011302253 | 0.279067358 | 1.946834975 |
| 83593 | RASSF5 | 1.068198163 | 0.025725072 | 0.390204824 | 1.589643401 |
| 64093 | SMOC1 | 1.05986236 | 0.010228914 | 0.267177232 | 1.990170473 |
| 54742 | LY6K | 1.041985934 | 0.07597113 | 0.498711659 | 1.119351414 |
| 677781 | SCARNA16 | 1.04059816 | 0.118960757 | 0.498711659 | 0.924596281 |
| 728441 | GGT2 | 1.035915092 | 0.012385445 | 0.290268247 | 1.907088385 |
| 677797 | SNORA7B | 1.032997689 | 0.115722095 | 0.498711659 | 0.936583713 |
| 2170 | FABP3 | 1.031841741 | 0.010657202 | 0.271648091 | 1.972356802 |
| 9796 | PHYHIP | 1.030721752 | 0.126068741 | 0.498711659 | 0.899392584 |
| 554210 | MIR429 | 1.029793823 | 0.051159047 | 0.493681625 | 1.291077554 |
| 5724 | PTAFR | 1.029431061 | 0.029512469 | 0.415060206 | 1.529994456 |
| 56121 | PCDHB15 | 1.009941429 | 0.025938868 | 0.391981984 | 1.586048981 |
| 960 | CD44 | 1.006861914 | 0.031191475 | 0.425126334 | 1.505964088 |
| 29986 | SLC39A2 | 0.95447507 | 0.17624372 | 0.498711659 | 0.753886349 |
| 57717 | PCDHB16 | 0.942827176 | 0.03217374 | 0.433178454 | 1.492498452 |
| 943 | TNFRSF8 | 0.930776631 | 0.161093277 | 0.498711659 | 0.792922584 |
| 731656 | LINC01348 | 0.928313444 | 0.022271099 | 0.369385425 | 1.652258352 |
| 3918 | LAMC2 | 0.920190292 | 0.021640183 | 0.36479653 | 1.664739071 |
| 4856 | CCN3 | 0.917474403 | 0.024155598 | 0.379578428 | 1.616982207 |
| 100874091 | TM4SF1-AS1 | 0.903217382 | 0.039532649 | 0.469284935 | 1.403044084 |
| 53405 | CLIC5 | 0.90085217 | 0.047835653 | 0.493681625 | 1.320248293 |
| 79094 | CHAC1 | 0.888025506 | 0.030656946 | 0.421546786 | 1.513471111 |
| 2634 | GBP2 | 0.882758003 | 0.029925235 | 0.417146318 | 1.52396243 |
| 4634 | MYL3 | 0.875120596 | 0.027185594 | 0.400439957 | 1.565661173 |
| 170591 | S100Z | 0.858595913 | 0.091754672 | 0.498711659 | 1.037371813 |
| 80765 | STARD5 | 0.855938841 | 0.018319045 | 0.342246301 | 1.737097171 |
| 3640 | INSL3 | 0.851933522 | 0.178678834 | 0.498711659 | 0.74792689 |
| 654 | BMP6 | 0.835157473 | 0.021848993 | 0.366451364 | 1.660568574 |
| 84225 | ZMYND15 | 0.832246173 | 0.05691531 | 0.498711659 | 1.244770894 |
| 80342 | TRAF3IP3 | 0.823486754 | 0.016349975 | 0.32809721 | 1.786482907 |
| 4608 | MYBPH | 0.818573664 | 0.135053107 | 0.498711659 | 0.86949542 |
| 339942 | H1-10-AS1 | 0.815113806 | 0.079844492 | 0.498711659 | 1.097755038 |
| 57689 | LRRC4C | 0.812452245 | 0.057610174 | 0.498711659 | 1.239500813 |
| 4597 | MVD | 0.811092007 | 0.011057912 | 0.276399141 | 1.956326871 |
| 7482 | WNT2B | 0.810149553 | 0.030143122 | 0.41791598 | 1.520811769 |
| 6558 | SLC12A2 | 0.798827281 | 0.012726884 | 0.293804784 | 1.895277914 |
| 2086 | ERV3-1 | 0.798263667 | 0.024389347 | 0.380516496 | 1.612799827 |
| 57822 | GRHL3 | 0.795333604 | 0.056123867 | 0.498711659 | 1.250852413 |
| 117157 | SH2D1B | 0.789689592 | 0.028522955 | 0.40863234 | 1.544805483 |
| 101928100 | KLRK1-AS1 | 0.784747998 | 0.023002999 | 0.374689142 | 1.638215539 |
| 3695 | ITGB7 | 0.782277842 | 0.142834396 | 0.498711659 | 0.845167197 |
| 23149 | FCHO1 | 0.771600817 | 0.074197311 | 0.498711659 | 1.129611834 |
| 5125 | PCSK5 | 0.769467829 | 0.010989808 | 0.276155403 | 1.959009895 |
| 4248 | MGAT3 | 0.762441785 | 0.015255583 | 0.317582309 | 1.816571191 |
| 10811 | NOXA1 | 0.751660648 | 0.011529103 | 0.282458046 | 1.938204481 |
| 284348 | LYPD5 | 0.734468618 | 0.034777388 | 0.441836216 | 1.458703039 |
| 131034 | CPNE4 | 0.72982452 | 0.047391904 | 0.493681625 | 1.324295843 |
| 728233 | PI4KAP1 | 0.72486306 | 0.055264608 | 0.498487467 | 1.257552906 |
| 101927668 | LOC101927668 | 0.718182225 | 0.053644601 | 0.49394054 | 1.270473981 |
| 130612 | TMEM198 | 0.710330427 | 0.049185428 | 0.493681625 | 1.308163545 |
| 6035 | RNASE1 | 0.707708587 | 0.023631695 | 0.376752455 | 1.626505127 |
| 3656 | IRAK2 | 0.687648428 | 0.020464314 | 0.35878954 | 1.689002809 |
| 794 | CALB2 | 0.675797173 | 0.185825557 | 0.498711659 | 0.730894557 |
| 257044 | CATSPERE | 0.674261979 | 0.026795603 | 0.398565301 | 1.571936465 |
| 3487 | IGFBP4 | 0.66973993 | 0.015980735 | 0.324355224 | 1.79640325 |
| 283152 | CCDC153 | 0.661771052 | 0.123253912 | 0.498711659 | 0.909199288 |
| 7483 | WNT9A | 0.654768859 | 0.02708363 | 0.400439957 | 1.567293128 |
| 406985 | MIR200C | 0.652210251 | 0.201877077 | 0.498711659 | 0.694912992 |
| 101926975 | LINC01844 | 0.652210251 | 0.235344814 | 0.500356689 | 0.628295367 |
| 54441 | STAG3L1 | 0.650184317 | 0.013204833 | 0.297462033 | 1.879267087 |
| 101927132 | LINC02133 | 0.645503042 | 0.112677278 | 0.498711659 | 0.948163653 |
| 1437 | CSF2 | 0.645503042 | 0.132143999 | 0.498711659 | 0.878952555 |
| 4511 | TRNC | 0.642852614 | 0.015627883 | 0.321255914 | 1.806099849 |
| 10252 | SPRY1 | 0.641286836 | 0.011947828 | 0.287263813 | 1.922711038 |
| 284578 | MFSD4A-AS1 | 0.637884876 | 0.06115151 | 0.498711659 | 1.213592815 |
| 11211 | FZD10 | 0.637884876 | 0.26248957 | 0.532423126 | 0.580887949 |
| 90019 | SYT8 | 0.630380675 | 0.037738306 | 0.459707935 | 1.423217598 |
| 11247 | NXPH4 | 0.629551316 | 0.242996765 | 0.510302888 | 0.614399508 |
| 130271 | PLEKHH2 | 0.629184052 | 0.048346182 | 0.493681625 | 1.315637817 |
| 100873982 | ABCC5-AS1 | 0.626709097 | 0.2630471 | 0.533097024 | 0.579966482 |
| 574407 | OBSCN-AS1 | 0.624088951 | 0.023784725 | 0.377366068 | 1.623701866 |
| 3428 | IFI16 | 0.621925364 | 0.015818099 | 0.323365671 | 1.800845711 |
| 1291 | COL6A1 | 0.617280175 | 0.028219453 | 0.406746893 | 1.549451409 |
| 439931 | THAP7-AS1 | 0.615821371 | 0.026783082 | 0.398565301 | 1.572139449 |
| 286122 | C8orf31 | 0.614825776 | 0.038587611 | 0.465675598 | 1.413552108 |
| 50512 | PODXL2 | 0.604413046 | 0.029392079 | 0.414029782 | 1.531769694 |
| 7804 | LRP8 | 0.600811486 | 0.023733098 | 0.377366068 | 1.624645567 |
| 192668 | CYS1 | 0.600113386 | 0.206449512 | 0.498711659 | 0.685186139 |
| 7498 | XDH | 0.599023898 | 0.019122773 | 0.347625567 | 1.71844913 |
| 100132352 | FRG1HP | 0.59277155 | 0.017172208 | 0.333063425 | 1.76517386 |
| 3177 | SLC29A2 | 0.590473925 | 0.011271712 | 0.278798139 | 1.948010116 |
| 54855 | TENT5C | 0.587583176 | 0.027429654 | 0.402306312 | 1.561779671 |
| 256472 | TMEM151A | 0.586219634 | 0.274341875 | 0.545807075 | 0.561707897 |
| 961 | CD47 | 0.581430355 | 0.041941275 | 0.478959251 | 1.377358371 |
| 91010 | FMNL3 | 0.577184721 | 0.04243014 | 0.480765251 | 1.372325535 |
| 39 | ACAT2 | 0.571332281 | 0.012682188 | 0.293011164 | 1.896805813 |
| 105369971 | FIGNL2-DT | 0.567914104 | 0.024339648 | 0.380365601 | 1.613685707 |
| 10718 | NRG3 | 0.565792379 | 0.016276443 | 0.327779859 | 1.788440498 |
| 54626 | HES2 | 0.563760562 | 0.035862457 | 0.447524535 | 1.445359959 |
| 101929490 | NA (ENTREZID: 101929490) | 0.561096253 | 0.100302797 | 0.498711659 | 0.998686956 |
| 54507 | ADAMTSL4 | 0.559077408 | 0.011526901 | 0.282458046 | 1.938287437 |
| 100616403 | MIR4691 | 0.553946517 | 0.187062144 | 0.498711659 | 0.728014092 |
| 283848 | CES4A | 0.552170188 | 0.051223418 | 0.493681625 | 1.290531446 |
| 84969 | TOX2 | 0.547840312 | 0.36421387 | 0.646156072 | 0.438643519 |
| 11076 | TPPP | 0.538169149 | 0.042929165 | 0.482761842 | 1.367247559 |
| 10863 | ADAM28 | 0.525348998 | 0.039275527 | 0.468074463 | 1.405877979 |
| 94005 | PIGS | 0.517727389 | 0.046454391 | 0.493681625 | 1.332973229 |
| 4322 | MMP13 | 0.511686587 | 0.130448188 | 0.498711659 | 0.884561949 |
| 64135 | IFIH1 | 0.506414005 | 0.032611723 | 0.435178498 | 1.486626255 |
| 4319 | MMP10 | 0.505601568 | 0.017512929 | 0.335951946 | 1.756641213 |
| 3557 | IL1RN | 0.505453624 | 0.018385828 | 0.342774403 | 1.735516807 |
| 90271 | OLMALINC | 0.499537095 | 0.117376702 | 0.498711659 | 0.930418097 |
| 53827 | FXYD5 | 0.492979939 | 0.016071088 | 0.325657354 | 1.793954721 |
| 80301 | PLEKHO2 | 0.489492907 | 0.021236628 | 0.361302608 | 1.67291444 |
| 29937 | NENF | 0.486180432 | 0.089360592 | 0.498711659 | 1.048853963 |
| 5420 | PODXL | 0.483754511 | 0.068234944 | 0.498711659 | 1.165993161 |
| 23138 | N4BP3 | 0.481280888 | 0.029028591 | 0.412044825 | 1.537174044 |
| 79616 | CCNJL | 0.470810701 | 0.010867252 | 0.273802674 | 1.963880262 |
| 79844 | ZDHHC11 | 0.468317934 | 0.17719902 | 0.498711659 | 0.751538684 |
| 127294 | MYOM3 | 0.46807784 | 0.050894884 | 0.493681625 | 1.293325871 |
| 440456 | PLEKHM1P1 | 0.465246345 | 0.03310718 | 0.435203051 | 1.48007781 |
| 3625 | INHBB | 0.461509561 | 0.025581835 | 0.388863063 | 1.592068307 |
| 9052 | GPRC5A | 0.46042261 | 0.044381085 | 0.491498014 | 1.352802085 |
| 3156 | HMGCR | 0.454712143 | 0.01653071 | 0.329165154 | 1.781708493 |
| 91544 | UBXN11 | 0.453943757 | 0.150891463 | 0.498711659 | 0.821335331 |
| 29964 | PRICKLE4 | 0.453097478 | 0.075671579 | 0.498711659 | 1.121067204 |
| 3783 | KCNN4 | 0.450940643 | 0.403381263 | 0.682761741 | 0.394284279 |
| 79411 | GLB1L | 0.432055735 | 0.023301796 | 0.375753064 | 1.632610604 |
| 399473 | SPRED3 | 0.430466315 | 0.05085302 | 0.493681625 | 1.293683251 |
| 65999 | LRRC61 | 0.430132859 | 0.013648127 | 0.30182132 | 1.864926945 |
| 200150 | PLD5 | 0.428814862 | 0.051764864 | 0.493681625 | 1.285964923 |
| 79891 | ZNF671 | 0.417488896 | 0.019191289 | 0.34814644 | 1.716895855 |
| 2944 | GSTM1 | 0.416962089 | 0.12265141 | 0.498711659 | 0.911327455 |
| 7180 | CRISP2 | 0.415386204 | 0.570066698 | 0.793557519 | 0.244074329 |
| 254531 | LPCAT4 | 0.414374051 | 0.046947137 | 0.493681625 | 1.328390887 |
| 24138 | IFIT5 | 0.412824852 | 0.099326845 | 0.498711659 | 1.002933359 |
| 3417 | IDH1 | 0.408745407 | 0.034119416 | 0.439382009 | 1.466998411 |
| 101928079 | SLC44A3-AS1 | 0.405295716 | 0.039734325 | 0.469324524 | 1.40083416 |
| 126567 | C2CD4C | 0.404645864 | 0.040807758 | 0.473552509 | 1.389257265 |
| 27101 | CACYBP | 0.401994451 | 0.018705929 | 0.344643459 | 1.728020718 |
| 100463486 | MTRNR2L8 | 0.401241477 | 0.01121497 | 0.278528703 | 1.950201884 |
| 148304 | C1orf74 | 0.399487522 | 0.096498534 | 0.498711659 | 1.015479284 |
| 200942 | KLHDC8B | 0.398752463 | 0.014479643 | 0.308670759 | 1.839242146 |
| 912 | CD1D | 0.396407373 | 0.273641513 | 0.545038847 | 0.562818017 |
| 91316 | GUSBP11 | 0.389865245 | 0.208918013 | 0.498711659 | 0.680024113 |
| 5002 | SLC22A18 | 0.389706125 | 0.159277775 | 0.498711659 | 0.79784482 |
| 79605 | PGBD5 | 0.389434579 | 0.182908202 | 0.498711659 | 0.737766819 |
| 217 | ALDH2 | 0.388128758 | 0.049491589 | 0.493681625 | 1.305468602 |
| 10379 | IRF9 | 0.387232384 | 0.058361701 | 0.498711659 | 1.233872059 |
| 728975 | LOC728975 | 0.384391779 | 0.026547723 | 0.396720545 | 1.575972722 |
| 440279 | UNC13C | 0.377982564 | 0.036274061 | 0.449586194 | 1.440403821 |
| 351 | APP | 0.377481071 | 0.055587664 | 0.498711659 | 1.255021576 |
| 1292 | COL6A2 | 0.374295831 | 0.021185731 | 0.361085741 | 1.673956546 |
| 7442 | TRPV1 | 0.373845114 | 0.030120172 | 0.41791598 | 1.521142552 |
| 11078 | TRIOBP | 0.372880997 | 0.019285727 | 0.348703239 | 1.714763985 |
| 311 | ANXA11 | 0.368054375 | 0.06503737 | 0.498711659 | 1.186837029 |
| 101929395 | LINC01752 | 0.367687746 | 0.461464667 | 0.727175637 | 0.335861546 |
| 84033 | OBSCN | 0.364880869 | 0.029124185 | 0.412252339 | 1.535746219 |
| 114787 | GPRIN1 | 0.364094881 | 0.014457118 | 0.308670759 | 1.839918274 |
| 2707 | GJB3 | 0.363360109 | 0.135701465 | 0.498711659 | 0.867415464 |
| 2207 | FCER1G | 0.362419437 | 0.057809959 | 0.498711659 | 1.237997339 |
| 152519 | NIPAL1 | 0.36090747 | 0.109331625 | 0.498711659 | 0.961254197 |
| 1894 | ECT2 | 0.360381703 | 0.014977134 | 0.313625167 | 1.824571285 |
| 124222 | PAQR4 | 0.354113372 | 0.06507013 | 0.498711659 | 1.186618326 |
| 4233 | MET | 0.352930243 | 0.012633023 | 0.292441787 | 1.898492713 |
| 10625 | IVNS1ABP | 0.350522783 | 0.044403139 | 0.491550538 | 1.352586327 |
| 11118 | BTN3A2 | 0.3482241 | 0.100409871 | 0.498711659 | 0.998223591 |
| 10654 | PMVK | 0.345093911 | 0.016855838 | 0.331482705 | 1.773249651 |
| 84674 | CARD6 | 0.343519556 | 0.023592129 | 0.376535449 | 1.627232866 |
| 55224 | ETNK2 | 0.331455906 | 0.018611412 | 0.34408973 | 1.730220677 |
| 23385 | NCSTN | 0.331389868 | 0.040869312 | 0.473860392 | 1.388602673 |
| 3949 | LDLR | 0.329409261 | 0.012434432 | 0.290898875 | 1.905374048 |
| 57658 | CALCOCO1 | 0.32886482 | 0.115260467 | 0.498711659 | 0.938319625 |
| 2827 | GPR3 | 0.324637252 | 0.29155374 | 0.565953002 | 0.535281383 |
| 7294 | TXK | 0.32068607 | 0.272674387 | 0.543912134 | 0.564355655 |
| 3655 | ITGA6 | 0.317447124 | 0.057666338 | 0.498711659 | 1.239077627 |
| 129790 | NA (ENTREZID: 129790) | 0.314382787 | 0.082191584 | 0.498711659 | 1.08517265 |
| 4166 | CHST6 | 0.314010545 | 0.457985789 | 0.724536279 | 0.339147998 |
| 55827 | DCAF6 | 0.312265652 | 0.069018799 | 0.498711659 | 1.161032602 |
| 9266 | CYTH2 | 0.310142546 | 0.047195487 | 0.493681625 | 1.326099528 |
| 64855 | NIBAN2 | 0.308383559 | 0.010844063 | 0.273681251 | 1.964807968 |
| 4179 | CD46 | 0.303861564 | 0.058775478 | 0.498711659 | 1.23080383 |
| 4643 | MYO1E | 0.302838726 | 0.010617026 | 0.271249333 | 1.973997119 |
| 3622 | ING2 | 0.301919048 | 0.037733601 | 0.459707935 | 1.423271747 |
| 286410 | ATP11C | 0.300073943 | 0.073896692 | 0.498711659 | 1.131375002 |
| 9334 | B4GALT5 | 0.298738175 | 0.100578493 | 0.498711659 | 0.997494876 |
| 3993 | LLGL2 | 0.296837621 | 0.022710461 | 0.373184913 | 1.64377405 |
| 9580 | SOX13 | 0.295410715 | 0.052713517 | 0.493681625 | 1.278078007 |
| 1604 | CD55 | 0.293234765 | 0.011242631 | 0.278528703 | 1.949132043 |
| 57228 | SMAGP | 0.290041358 | 0.023053722 | 0.374920061 | 1.637258948 |
| 3673 | ITGA2 | 0.28790864 | 0.011973721 | 0.28728883 | 1.921770866 |
| 11066 | SNRNP35 | 0.287263657 | 0.018904766 | 0.346170149 | 1.723428694 |
| 90634 | N4BP2L1 | 0.286801897 | 0.080688502 | 0.498711659 | 1.093188347 |
| 81544 | GDPD5 | 0.28554029 | 0.03034301 | 0.419226436 | 1.51794134 |
| 124565 | SLC38A10 | 0.284238495 | 0.055565885 | 0.498711659 | 1.255191764 |
| 23015 | GOLGA8A | 0.281636489 | 0.033001755 | 0.435178498 | 1.481462964 |
| 100128770 | LOC100128770 | 0.281354855 | 0.097626561 | 0.498711659 | 1.010432009 |
| 4690 | NCK1 | 0.280337256 | 0.071668793 | 0.498711659 | 1.14466991 |
| 6844 | VAMP2 | 0.279707566 | 0.027743512 | 0.403781146 | 1.556838563 |
| 442582 | STAG3L2 | 0.279328064 | 0.218012661 | 0.498711659 | 0.661518284 |
| 9605 | VPS9D1 | 0.278967863 | 0.032984096 | 0.435178498 | 1.481695414 |
| 5546 | PRCC | 0.278399916 | 0.030781286 | 0.422323463 | 1.51171324 |
| 84777 | DLGAP1-AS2 | 0.27603853 | 0.392252746 | 0.671976876 | 0.406434007 |
| 9388 | LIPG | 0.275573009 | 0.13342144 | 0.498711659 | 0.874774376 |
| 23170 | TTLL12 | 0.274036913 | 0.023928124 | 0.37830683 | 1.621091349 |
| 342184 | FMN1 | 0.272613287 | 0.559181336 | 0.786582484 | 0.252447333 |
| 5639 | PRRG2 | 0.272117482 | 0.085154066 | 0.498711659 | 1.06979461 |
| 259173 | ALS2CL | 0.271600702 | 0.047274538 | 0.493681625 | 1.325372707 |
| 3797 | KIF3C | 0.270191248 | 0.064730048 | 0.498711659 | 1.188894071 |
| 105370548 | NA (ENTREZID: 105370548) | 0.269413043 | 0.273081657 | 0.544227516 | 0.563707471 |
| 146439 | BICDL2 | 0.267482281 | 0.126260404 | 0.498711659 | 0.898732825 |
| 3032 | HADHB | 0.266451997 | 0.015139269 | 0.315510963 | 1.819895094 |
| 4513 | COX2 | 0.266337955 | 0.08869204 | 0.498711659 | 1.052115356 |
| 4878 | NPPA | 0.264983672 | 0.438207279 | 0.712447507 | 0.358320413 |
| 253980 | KCTD13 | 0.26285502 | 0.046301297 | 0.493681625 | 1.334406843 |
| 11328 | FKBP9 | 0.261240724 | 0.059469483 | 0.498711659 | 1.225705837 |
| 860 | RUNX2 | 0.259749701 | 0.250109759 | 0.517515603 | 0.601869362 |
| 84807 | NFKBID | 0.259391655 | 0.096150193 | 0.498711659 | 1.01704984 |
| 79083 | MLPH | 0.258738347 | 0.523297169 | 0.76364923 | 0.281251615 |
| 56063 | TMEM234 | 0.256684264 | 0.146912328 | 0.498711659 | 0.832941759 |
| 81606 | LBH | 0.25388364 | 0.207854298 | 0.498711659 | 0.682240991 |
| 54344 | DPM3 | 0.250412397 | 0.102088624 | 0.498711659 | 0.99102265 |
| 83986 | FAM234A | 0.250344333 | 0.015780309 | 0.323291405 | 1.801884497 |
| 53 | ACP2 | 0.248958977 | 0.020677249 | 0.359184257 | 1.684507242 |
| 4549 | RNR1 | 0.248559063 | 0.085164987 | 0.498711659 | 1.069738916 |
| 54972 | TMEM132A | 0.24810384 | 0.198522752 | 0.498711659 | 0.702189713 |
| 55733 | HHAT | 0.247411374 | 0.181551141 | 0.498711659 | 0.741001017 |
| 25994 | HIGD1A | 0.247311589 | 0.117890052 | 0.498711659 | 0.928522841 |
| 85407 | NKD1 | 0.247290477 | 0.674894781 | 0.848951784 | 0.17076393 |
| 9191 | DEDD | 0.246249776 | 0.021397819 | 0.362308335 | 1.66963049 |
| 1592 | CYP26A1 | 0.245249961 | 0.418650215 | 0.696138525 | 0.378148681 |
| 55972 | SLC25A40 | 0.245010833 | 0.046447384 | 0.493681625 | 1.333038741 |
| 100130418 | CECR7 | 0.244930496 | 0.303357834 | 0.580213236 | 0.518044785 |
| 353149 | TBC1D26 | 0.244930496 | 0.306406664 | 0.583216064 | 0.513701794 |
| 152065 | C3orf22 | 0.244930496 | 0.445758083 | 0.718779791 | 0.350900773 |
| 102464835 | NA (ENTREZID: 102464835) | 0.244930496 | 0.445758083 | 0.718779791 | 0.350900773 |
| 408187 | SPINK14 | 0.244930496 | 0.504885472 | 0.752970137 | 0.296807126 |
| 105369175 | NA (ENTREZID: 105369175) | 0.244930496 | 0.529442571 | 0.767841918 | 0.276181141 |
| 285084 | LINC01305 | 0.244930496 | 0.552461895 | 0.782808711 | 0.257697671 |
| 105369728 | LOC105369728 | 0.244930496 | 0.555192796 | 0.783993452 | 0.255556178 |
| 84066 | TEX35 | 0.244930496 | 0.592521902 | 0.806589933 | 0.227295592 |
| 163404 | PLPPR5 | 0.244930496 | 0.597279185 | 0.810075586 | 0.22382262 |
| 102465428 | MIR6717 | 0.244930496 | 0.60217725 | 0.812906901 | 0.220275656 |
| 151790 | WDR49 | 0.244930496 | 0.61070336 | 0.816043383 | 0.214169691 |
| 64838 | FNDC4 | 0.244298374 | 0.075569133 | 0.498711659 | 1.12165556 |
| 6925 | TCF4 | 0.2435885 | 0.102881144 | 0.498711659 | 0.987664215 |
| 54332 | GDAP1 | 0.243246154 | 0.028074808 | 0.405844274 | 1.551683205 |
| 574412 | MIR452 | 0.242418316 | 0.594899313 | 0.8085471 | 0.225556533 |
| 26778 | SNORA70 | 0.242258476 | 0.32032868 | 0.598711161 | 0.494404176 |
| 90231 | KIAA2013 | 0.241278032 | 0.016079296 | 0.325657354 | 1.79373297 |
| 9138 | ARHGEF1 | 0.239709204 | 0.017316488 | 0.334131729 | 1.761540184 |
| 6667 | SP1 | 0.238698935 | 0.053200682 | 0.493681625 | 1.2740828 |
| 5045 | FURIN | 0.236065996 | 0.067801241 | 0.498711659 | 1.168762357 |
| 3455 | IFNAR2 | 0.231270031 | 0.042132273 | 0.479689616 | 1.37538511 |
| 10956 | OS9 | 0.231236709 | 0.023166081 | 0.375753064 | 1.635147429 |
| 197370 | NSMCE1 | 0.229709041 | 0.092153642 | 0.498711659 | 1.035487496 |
| 3675 | ITGA3 | 0.229436273 | 0.015566352 | 0.320526871 | 1.807813153 |
| 6810 | STX4 | 0.229397149 | 0.06976108 | 0.498711659 | 1.156386805 |
| 201283 | AMZ2P1 | 0.227839242 | 0.030817194 | 0.422323463 | 1.511206908 |
| 285 | ANGPT2 | 0.22521032 | 0.628329233 | 0.825685328 | 0.201812734 |
| 23162 | MAPK8IP3 | 0.224959703 | 0.066432688 | 0.498711659 | 1.177618175 |
| 6549 | SLC9A2 | 0.224678765 | 0.188624266 | 0.498711659 | 0.724402437 |
| 57216 | VANGL2 | 0.223659754 | 0.067917898 | 0.498711659 | 1.168015764 |
| 440926 | H3P6 | 0.22349239 | 0.089147646 | 0.498711659 | 1.04989012 |
| 4259 | MGST3 | 0.223245556 | 0.014077992 | 0.306540571 | 1.851459286 |
| 7832 | BTG2 | 0.221225977 | 0.012532417 | 0.292165831 | 1.901965163 |
| 4242 | MFNG | 0.217325859 | 0.29244573 | 0.567057942 | 0.533954715 |
| 3853 | KRT6A | 0.205454 | 0.333257074 | 0.613832033 | 0.477220623 |
| 7391 | USF1 | 0.198354365 | 0.123750742 | 0.498711659 | 0.907452188 |
| 90780 | PYGO2 | 0.197402118 | 0.020271013 | 0.357271699 | 1.693124548 |
| 5993 | RFX5 | 0.196873404 | 0.013123507 | 0.297030712 | 1.881950093 |
| 9863 | MAGI2 | 0.194332648 | 0.091213955 | 0.498711659 | 1.039938713 |
| 58 | ACTA1 | 0.194002093 | 0.827686591 | 0.922449201 | 0.082134081 |
| 23432 | GPR161 | 0.193133755 | 0.44465083 | 0.71778626 | 0.351980893 |
| 6768 | ST14 | 0.19306404 | 0.359718171 | 0.641595319 | 0.444037623 |
| 51332 | SPTBN5 | 0.191465772 | 0.448332718 | 0.720616299 | 0.348399566 |
| 105371697 | NA (ENTREZID: 105371697) | 0.189137842 | 0.703757309 | 0.862471085 | 0.152577082 |
| 9261 | MAPKAPK2 | 0.187216817 | 0.084009272 | 0.498711659 | 1.075672779 |
| 84614 | ZBTB37 | 0.186271646 | 0.21545836 | 0.498711659 | 0.66663665 |
| 121274 | ZNF641 | 0.185398352 | 0.063052086 | 0.498711659 | 1.200300541 |
| 10865 | ARID5A | 0.18446034 | 0.257877072 | 0.526944405 | 0.58858727 |
| 84070 | FAM186B | 0.182365674 | 0.596857012 | 0.809685795 | 0.2241297 |
| 116254 | GINM1 | 0.180627551 | 0.055126103 | 0.498253138 | 1.258642708 |
| 84451 | MAP3K21 | 0.178133703 | 0.08904232 | 0.498711659 | 1.050403533 |
| 51003 | MED31 | 0.177509342 | 0.145039015 | 0.498711659 | 0.838515158 |
| 100507458 | ZNF213-AS1 | 0.175075199 | 0.399159541 | 0.678568831 | 0.398853485 |
| 9910 | RABGAP1L | 0.173647035 | 0.023498422 | 0.376120459 | 1.628961301 |
| 100505678 | STARD4-AS1 | 0.173324036 | 0.265666778 | 0.536347087 | 0.575662751 |
| 2132 | EXT2 | 0.167410866 | 0.200255104 | 0.498711659 | 0.698416406 |
| 25780 | RASGRP3 | 0.165104372 | 0.554634035 | 0.783774136 | 0.255993484 |
| 9708 | PCDHGA8 | 0.163933277 | 0.438180032 | 0.712444 | 0.358347417 |
| 9322 | TRIP10 | 0.162229876 | 0.071724448 | 0.498711659 | 1.144332785 |
| 5792 | PTPRF | 0.161707825 | 0.045359639 | 0.493681625 | 1.34333041 |
| 638 | BIK | 0.161447855 | 0.42554038 | 0.70218614 | 0.371059223 |
| 23365 | ARHGEF12 | 0.156516804 | 0.010070696 | 0.26477537 | 1.996940514 |
| 153090 | DAB2IP | 0.152307186 | 0.027145364 | 0.400439957 | 1.56630433 |
| 376497 | SLC27A1 | 0.152267319 | 0.473044162 | 0.734386978 | 0.325098313 |
| 23324 | MAN2B2 | 0.151933803 | 0.010353872 | 0.268064651 | 1.984897208 |
| 16 | AARS1 | 0.15049541 | 0.205518246 | 0.498711659 | 0.687149615 |
| 23607 | CD2AP | 0.149425593 | 0.087983517 | 0.498711659 | 1.055598682 |
| 54914 | FOCAD | 0.148809458 | 0.107686021 | 0.498711659 | 0.96784067 |
| 57149 | LYRM1 | 0.148726951 | 0.023555476 | 0.376408354 | 1.627908115 |
| 4358 | MPV17 | 0.148257332 | 0.069035813 | 0.498711659 | 1.160925556 |
| 79085 | SLC25A23 | 0.148210176 | 0.401254549 | 0.680744632 | 0.396580031 |
| 427 | ASAH1 | 0.147564038 | 0.106443489 | 0.498711659 | 0.972880899 |
| 6840 | SVIL | 0.146612916 | 0.170941623 | 0.498711659 | 0.767152177 |
| 220906 | WAC-AS1 | 0.143860222 | 0.177106299 | 0.498711659 | 0.751765992 |
| 2274 | FHL2 | 0.140545007 | 0.141655029 | 0.498711659 | 0.848768003 |
| 285989 | ZNF789 | 0.136418443 | 0.325687816 | 0.605071025 | 0.487198488 |
| 5352 | PLOD2 | 0.135520676 | 0.059830645 | 0.498711659 | 1.223076315 |
| 51278 | IER5 | 0.135447622 | 0.243491318 | 0.510932306 | 0.61351652 |
| 823 | CAPN1 | 0.134890765 | 0.056034365 | 0.498711659 | 1.251545545 |
| 4123 | MAN2C1 | 0.133341224 | 0.287032088 | 0.560640169 | 0.54206955 |
| 81844 | TRIM56 | 0.12950052 | 0.023263645 | 0.375753064 | 1.633322238 |
| 4301 | AFDN | 0.129287822 | 0.073325431 | 0.498711659 | 1.134745376 |
| 29911 | HOOK2 | 0.127465093 | 0.220746309 | 0.498711659 | 0.656106549 |
| 4540 | ND5 | 0.126064247 | 0.269770958 | 0.54090217 | 0.569004806 |
| 51341 | ZBTB7A | 0.125324452 | 0.07745851 | 0.498711659 | 1.110930861 |
| 8916 | HERC3 | 0.122500549 | 0.084056891 | 0.498711659 | 1.075426677 |
| 58527 | ABRACL | 0.120770761 | 0.118483017 | 0.498711659 | 0.926343896 |
| 54751 | FBLIM1 | 0.115626679 | 0.158861332 | 0.498711659 | 0.7989818 |
| 23252 | OTUD3 | 0.11562385 | 0.094106976 | 0.498711659 | 1.026378182 |
| 6282 | S100A11 | 0.112407027 | 0.221015565 | 0.498711659 | 0.65557714 |
| 100289678 | ZNF783 | 0.112183285 | 0.248861626 | 0.51609888 | 0.604042066 |
| 23353 | SUN1 | 0.109982035 | 0.195679721 | 0.498711659 | 0.70845418 |
| 1435 | CSF1 | 0.109245765 | 0.376552027 | 0.654317043 | 0.42417501 |
| 51022 | GLRX2 | 0.107621548 | 0.212733802 | 0.498711659 | 0.672163498 |
| 9260 | PDLIM7 | 0.107444655 | 0.46047353 | 0.726669879 | 0.33679533 |
| 51141 | INSIG2 | 0.104658229 | 0.400945648 | 0.680462317 | 0.396914496 |
| 4043 | LRPAP1 | 0.102357724 | 0.272817863 | 0.54401267 | 0.564127197 |
| 60313 | GPBP1L1 | 0.099679632 | 0.087998 | 0.498711659 | 1.055527198 |
| 22874 | PLEKHA6 | 0.097995291 | 0.319542572 | 0.59779616 | 0.495471273 |
| 56257 | MEPCE | 0.094648605 | 0.221466879 | 0.498711659 | 0.654691215 |
| 399665 | FAM102A | 0.093380012 | 0.343183923 | 0.623621049 | 0.464473066 |
| 2686 | GGT7 | 0.09311591 | 0.480495362 | 0.740169927 | 0.3183108 |
| 23268 | DNMBP | 0.089378129 | 0.13703762 | 0.498711659 | 0.863160193 |
| 825 | CAPN3 | 0.089278952 | 0.551891776 | 0.782501782 | 0.258146078 |
| 5826 | ABCD4 | 0.085764682 | 0.259840096 | 0.529545649 | 0.585293832 |
| 5366 | PMAIP1 | 0.080572203 | 0.358651235 | 0.640859721 | 0.445327669 |
| 788 | SLC25A20 | 0.077000118 | 0.342138329 | 0.622358606 | 0.46579827 |
| 56965 | PARP6 | 0.0738285 | 0.42516925 | 0.701831141 | 0.371438153 |
| 7871 | SLMAP | 0.066274604 | 0.19429204 | 0.498711659 | 0.711544992 |
| 117143 | TADA1 | 0.064042896 | 0.407153514 | 0.686364524 | 0.390241813 |
| 316 | AOX1 | 0.057057154 | 0.6454055 | 0.833126753 | 0.190167338 |
| 6470 | SHMT1 | 0.056685309 | 0.431459053 | 0.70702215 | 0.365060414 |
| 11282 | MGAT4B | 0.056070413 | 0.323263691 | 0.602182806 | 0.490443073 |
| 9870 | AREL1 | 0.054593034 | 0.353807489 | 0.635765325 | 0.451232979 |
| 1163 | CKS1B | 0.051734909 | 0.531545161 | 0.769003151 | 0.274459831 |
| 5087 | PBX1 | 0.049249007 | 0.702926416 | 0.86200334 | 0.153090136 |
| 284370 | ZNF615 | 0.046508298 | 0.662486609 | 0.843008955 | 0.178822896 |
| 2043 | EPHA4 | 0.046476044 | 0.841098919 | 0.929013338 | 0.075152925 |
| 255812 | SDHAP1 | 0.045180381 | 0.759831499 | 0.888671536 | 0.119282707 |
| 282974 | STK32C | 0.044071277 | 0.726195312 | 0.872817603 | 0.138946559 |
| 647024 | C6orf132 | 0.040525858 | 0.753067157 | 0.885768689 | 0.123166293 |
| 3419 | IDH3A | 0.039653137 | 0.527740009 | 0.766348448 | 0.27757998 |
| 54868 | TMEM104 | 0.03071339 | 0.613276096 | 0.817736962 | 0.212343963 |
| 8612 | PLPP2 | 0.027677225 | 0.854085167 | 0.935566249 | 0.068498821 |
| 283932 | FBXL19-AS1 | 0.025982449 | 0.7821897 | 0.899930168 | 0.106687907 |
| 170384 | FUT11 | 0.0174014 | 0.852712857 | 0.934646138 | 0.069197189 |
| 26258 | BLOC1S6 | 0.015368141 | 0.88190083 | 0.947754677 | 0.054580249 |
| 3688 | ITGB1 | 0.009319947 | 0.914111738 | 0.963918497 | 0.039000714 |
| 144165 | PRICKLE1 | 0.007832414 | 0.959809293 | 0.982933049 | 0.017815049 |
| 124995 | MRPL10 | 0.003664423 | 0.960052147 | 0.983061756 | 0.017705177 |
| 205251 | MTLN | 0.003361039 | 0.975550185 | 0.988570982 | 0.010750384 |
| 151393 | RMDN2 | 0.002816238 | 0.972522023 | 0.987135463 | 0.012100555 |
| 7991 | TUSC3 | 0.002630007 | 0.967004379 | 0.985361708 | 0.014571559 |

**Supplementary table 4: List of TaqMan gene expression assays used for RT-qPCR**

| Gene symbol | Corresponding TaqMan Assay ID |
| --- | --- |
| NANOG | Hs04399610_g1 |
| SOX2 | Hs1053049_s1 |
| POU5F1P | Hs01895061_u1 |
| Albumin | Hs00609411_m1 |
| AFP | Hs00173490_m1 |
| HNF4A | Hs002230853_m1 |
| GAPDH | Hs99999905_m1 |
| FAH | Hs00908451_m1 |
| ASGR1 | Hs01005019_m1 |
| TAT | Hs00944626_m1 |
| HPD | Hs00157976_m1 |
| GSTZ1 | Hs01041668_m1 |

**Supplementary table 5: List of primary antibodies used for immunofluorescence**

| Name of the antibody | Dilution | Product information |
| --- | --- | --- |
| Rabbit polyclonal anti human Albumin | 1:200 | Dako DK-A0001 |
| Rabbit polyclonal anti-Nanog | 1:200 | Novus Biologicals NB100-58842 |
| Rabbit polyclonal anti-human Oct3/4 | 1:100 | Novus Biologicals NBP3-05882 |
| Mouse monoclonal anti-human TRA-1-81 | 1:100 | Fisher Scientific 41-1100 |
| Mouse monoclonal anti-SSEA4 | 1:100 | Fisher Scientific 41-4000 |

**Supplementary table 6: List of secondary antibodies used for immunofluorescence**

| Name of the antibody | Product information |
| --- | --- |
| Alexa Fluor 555 goat anti-rabbit IgG | Invitrogen A21428 |
| Alexa Fluor 488 goat anti-mouse IgG | Invitrogen A11001 |
| Alexa Fluor 488 goat anti-rabbit IgG | Invitrogen A11008 |

**Supplementary table 7: List of fluorochrome-conjugated primary antibodies for FACS**

| Name of the antibody | Product information |
| --- | --- |
| PerCP-Cy 5.5 anti-human SOX17 | BD Bioscience 562387 |
| APC anti-human CD184 (CXCR4) | BD Bioscience 560936 |
| PE anti-human FOXA2 | BD Bioscience 561589 |
| PE anti-human EpCAM | BD Bioscience 347198 |
| FITC anti-human TRA1-60 | BD Bioscience 560876 |
| Alexa 647 anti-human Nanog | BD Bioscience 561300 |
| PE anti-human SSEA4 | BD Biosciences 560128 |
| APC anti-human Albumin | R&D Systems IC1455A |

**Supplementary table 8: List of primary antibodies used for western blot**

| Name of the antibody | Dilution | Product information |
| --- | --- | --- |
| Rabbit polyclonal anti-Fumarylacetoacetase (H-42) | 1:200 | Santa Cruz sc-67288 |
| Rabbit Polyclonal anti-beta actin | 1:10000 | Abcam ab8227 |

**Supplementary table 9: List of growth factors for HLC differentiation**

| Name of the growth factor | Product information |
| --- | --- |
| Activin A | R&D Systems 338-AC |
| CHIR99021 | Stem Cell Technologies 72054 |
| BMP4 | Peprotech 120-05ET |
| bFGF | Gibco PHG0266 |
| IWP-2 | Selleckchem S7085 |
| A83-01 | Reprocell 04-0014 |
| HGF | Peprotech 100-39H |
| Oncostatin M (OSM) | R&D Systems 295OM010 |
